# Reshaping epistatic network modules enhances wheat yield potential

**DOI:** 10.1101/2025.04.01.646715

**Authors:** Mou Yin, Haiming Han, Weihua Liu, Lingrang Kong, Xingpu Li, Wanquan Ji, Jizhong Wu, Guangbing Deng, Haibin Dong, Jinpeng Zhang, Shenghui Zhou, Zhimeng Zhang, Yuexuan Hou, Haojie Wang, Caihong Zhao, Xuanzhao Li, Qiaoling Luo, Xiaomin Guo, Xinming Yang, Hong-Qing Ling, Fei He, Lihui Li

**Author notes:** Correspondence (F.H.), (L.L.). These authors contributed equally: M.Y., H.H.

## Abstract

Increasing wheat yield is a key approach to ensuring global food security and an enduring focus in crop breeding. Here, we assembled a panel of 3,030 wheat lines to dissect the genetic mechanisms of yield improvement. We conducted large-scale field trials across five ecological environments over two consecutive years to evaluate yield performance and constructed a haplotype atlas using the Wheat 660K genotype array. 234 quantitative trait loci (QTLs) and 522 haplotype blocks (HBs) were identified for 14 traits through genome-wide association studies (GWAS), including 10 QTLs and 35 HBs associated with grain yield. Genomic analysis revealed that wheat yield improvement is primarily driven by changes in the epistatic network rather than the introduction of new haplotype segments. SNPs or HBs explain 53.3%-62.9% of the phenotypic variation in yield, whereas epistasis explains 70.4%, highlighting the role of epistasis in yield improvement. Moreover, yield improvement is significantly correlated with the accumulation of favorable epistatic modules. Pedigree-based analysis revealed that the modules remained relatively stable under the pressure of breeding selection, while new favorable modules were also created during hybrid breeding, highlighting that crop design breeding should be based on genetic modules. This study provides insights into genetic mechanism of wheat yield improvement and guidance for designing future wheat in the era of artificial intelligence.

## Introduction

As one of the three staple crops, wheat serves as a primary food source for more than 4 billion people worldwide, contributing approximately 20% of the calories and protein in the human diet^1^. With an increasing global population and the coming climate change^2–3^, improving wheat yield is a critical strategy for ensuring global food security and remains an enduring focus in crop breeding. Yield improvement is a complex optimization of compositive traits, resulting from the interaction and coordinated expression of numerous genes. Although many studies claim to have discovered genes or alleles that enhance yield, these genes often target single-plant yield, greenhouse yield, yield in a single environment, or yield component traits^4–5^. Thus, the practical deployment of these genes has limited yield gains in agricultural production^4–5^. The complex genetic mechanisms underlying crop yield improvement remain largely unknown, representing the most significant constraint in the era of intelligent design breeding.

China is a secondary origin center of wheat and has diverse ecological environments^6^. Its cultivation areas are widely distributed across 10 ecological zones, with each characterized by different climates, soils, and pest and disease risks^6^. Consequently, breeders have focused on developing region-specific wheat varieties adapted to different ecological conditions^6–7^. Through a century of breeding efforts, significant progress has been made in wheat improvement in China. From 1949 to 2024, the yield increased from 0.6 t ha^−1^ to 5.9 t ha^−1^, and the annual production rose from 13.8 million tons to 138 million tons (National Bureau of Statistics and FAO) (Supplementary Fig. 1). Currently, China is the world’s largest wheat producer, with an annual planting area exceeding 22 million hectares and a total production of over 135 million tons, accounting for approximately 17.5% of global wheat production. Therefore, modern wheat breeding in China represents a typical model for deciphering the genetic basis and mechanisms of yield improvement. To date, apart from the semi-dwarf genes^8–11^ and T1BL.1RS^6–7,12^, we know little about the genetic basis behind this sustained and significant yield increase. The genetic dissection of this case has a broad impact on future wheat yield improvement.

To dissect the genetic mechanisms of yield improvement, we constructed a panel of 3,030 wheat lines representing 70 years of wheat yield improvement history in China. To evaluate the yield potential of these varieties in practical production conditions, we conducted large-scale field trials across five ecological locations over two consecutive years. 14 agronomic traits, including yield per plot and yield component traits, were investigated. Our germplasm panel and field trials established the prerequisite conditions for uncovering the complex genetic mechanisms underlying yield improvement. Based on comprehensive and scientific experimental designs, we dissected the genetic basis of yield at the level of SNP, haplotype, epistasis, and network.

Our analysis revealed that yield improvement is controlled by a genetic epistatic network formed by thousands of alleles, rather than by the aggregation of a few alleles or haplotypes. Yield enhancement is essentially the result of the expansion of epistatic networks, the accumulation of superior modules, and the refinement of module components. Epistatic modules can be transmitted, aggregated, optimized, and formed through genetic recombination, and favorable modules maintain dynamic stability under the breeding selection. This is the first time that the genetic mechanisms of crop yield improvement have been revealed at the level of genetic network modules, emphasizing that future design breeding should focus on comprehensive improvement at the level of genetic modules. This landmark study uncovers the hidden principles already practiced in modern wheat breeding and provides critical guidance for genomic design breeding in the era of artificial intelligence.

## Results

### The genetic diversity of Chinese wheat cultivars

We assembled a panel of 3,030 wheat lines to investigate the mechanism of yield improvement (Supplementary Table 1). Among them, 2,772 modern Chinese cultivars were mainly from the four major wheat-growing zones, which account for over 90% of China’s wheat production^6^. These cultivars represent 70 years of breeding history in China, which began in the 1910s for selection breeding with landraces, and hybridization breeding in the 1950s with cultivar lines from other countries^6^. Since the pedigrees of Chinese cultivars were based on the landraces and introduced germplasms^6^, our panel included 65 Chinese landraces and 193 introduced lines from 25 other countries to quantify the genetic contributions of the two sources. Those landraces and introduced lines were selected because they played a critical role during the wheat breeding history in China. Cultivars of this panel included 1,170 landmark and elite lines that were widely cultivated from the 1950s to the 2010s. Besides, our panel also contained 1,602 advanced breeding lines from various national breeding programs, referred to as innovative germplasms, as they exhibit unique advantages in certain traits, such as yield, yield component traits, disease resistance, or stress tolerance. In summary, this germplasm panel not only contained diverse cultivars in China but also the donor populations of landraces and introduced lines.

The 3,030 wheat lines were genotyped with the Wheat 660K SNP array^13^, generating 201,740 variable SNP loci with a minor allele frequency of at least 0.01 and missing rate of no more than 0.4 (Supplementary Fig.2 and Supplementary Table 2). Wheat lines of different groups were largely clustered together on the phylogenetic tree, with certain degrees of admixtures between groups (Fig. 1a), which demonstrated wide germplasm exchanges between different breeding programs in China^6^. Most (70%) of the landraces were clustered near the root of the tree, while the rest were located in different cultivar groups, suggesting the majority of landraces have distinct genetic backgrounds compared to the cultivars. This phenomenon was also observed on the population PCA plot, where most landraces formed a separate cluster (Fig. 1b,c). Introduced wheat lines were predominantly located at the center of the germplasm panel (Fig. 1b), possibly reflecting their important genetic contribution to the cultivars. Wheat lines of different zones or subzones tended to cluster together, but there was no complete separation between the groups (Fig. 1b,c). Genetic assignment analysis with ADMIXTURE^14^ showed high mixing levels between groups, as well as again the unique genetic background of the landraces (Fig. 1d). Those results consistently demonstrated that there was a distinct population structure between the landraces and the cultivars and a high degree of genetic mixing between cultivar populations.

**Fig. 1.**
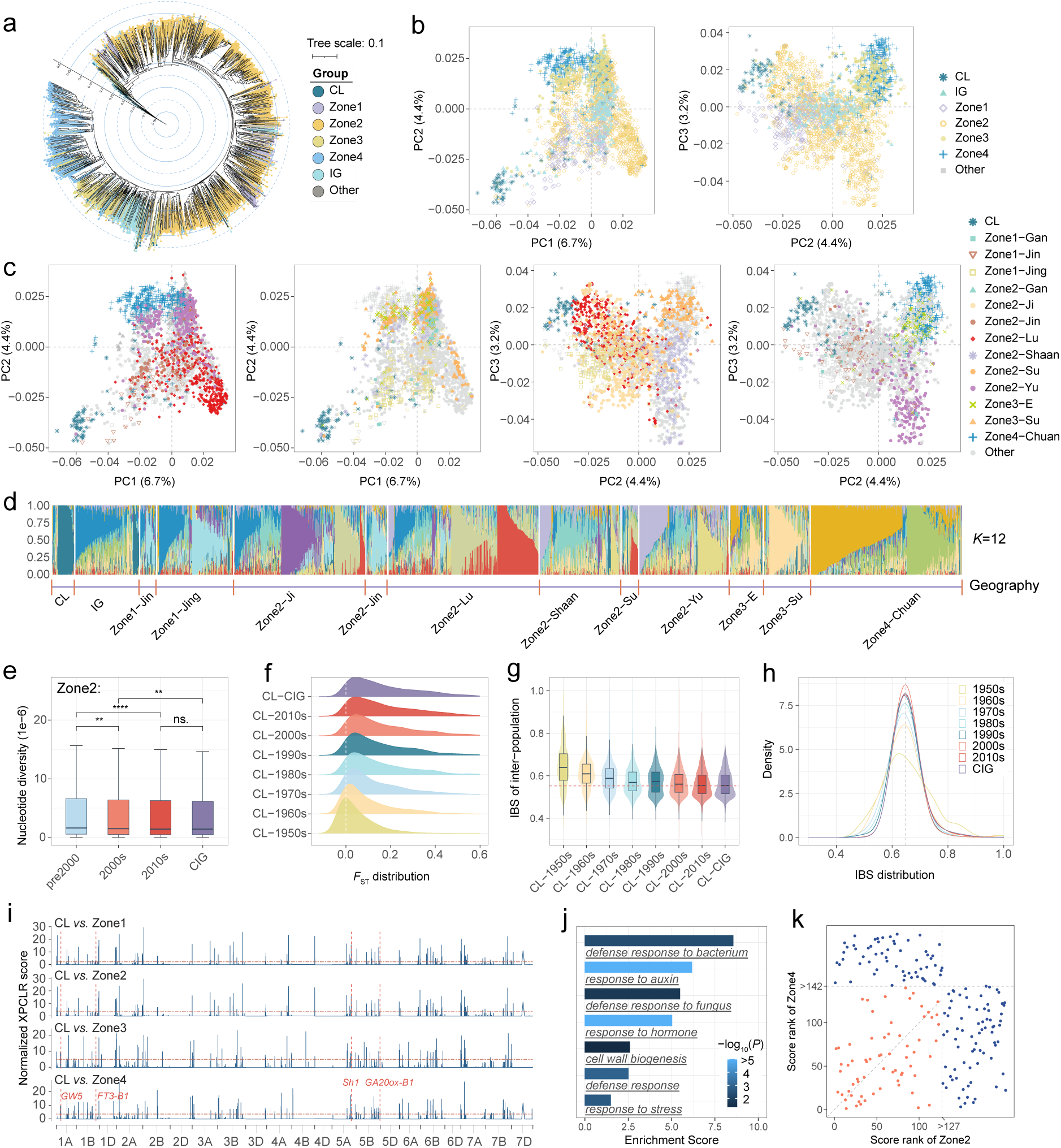
Population genetic characteristics. **a,** Neighbor-Joining tree of all accessions. **b,c,** Principal component analysis. CL: Chinese landrace; IG: introduced germplasms; Zone 1: Northern Winter Wheat Zone; Zone 2: Yellow and Huai River Valleys Winter Wheat Zone; Zone 3: Middle and Low Yangtze Valleys Winter Wheat Zone; Zone 4: Southern Winter Wheat Zone; Chuan: Sichuan; E: Hubei; Gan: Gansu; Ji: Hebei; Jin: Shanxi; Jing: Beijing; Lu: Shandong; Shaan: Shaanxi; Su: Jiangsu; Yu: Henan. **d,** Maximum likelihood estimation of individual ancestry coefficients using ADMIXTURE with *K*=12. **e,** The mean nucleotide diversity of different stages in Zone 2. Statistical significance for the different era groups within the Zone 2 group was determined using a one-tail t-test (ns: non-significant; ^⁎^*P*<0.05; ^⁎⁎^*P*<0.01; ^⁎⁎⁎^*P*<0.001; ^⁎⁎⁎⁎^*P*<0.0001). **f,g,** Distributions of fixation index (*F*_ST_) (f) and pairwise identity-by-state (IBS) (g) between CL and cultivars, respectively. CIG: Chinese innovative germplasm (advanced breeding lines). **h,** Distributions of pairwise IBS within populations. **i,** Identification of genetic improvement targets for different ecological zones using XP-CLR statistics. The Chinese landraces were considered as the reference population. The red dashed horizontal line represents the cut-off level of top 1% candidate genomic regions. Some loci were under selection across four ecological zones, with four genes associated with important agronomic traits labeled. **j**, Gene ontology (GO) enrichment of genes located in top 1% candidate regions of XPCLR. **k,** The ranking comparison of XP-CLR scores between Zone 2 and Zone 3. Higher rankings represent stronger selection intensity. The gray dashed line represents the top 1% threshold. Each dot represents a genomic region of 1 Mb. Red dots indicate loci under selection in both ecological zones, while dark blue dots indicate loci under selection in only one ecological zone.

For lines of Zone 2, nucleotide diversity (π) decreased from 5.5×10^−6^ in lines released before 2000 to 5.1×10^−6^ in those released during the 2010s (Fig. 1e and Supplementary Table 3). The population divergence index (*F*_ST_) between landraces and cultivars continued to increase from 0.083 in the 1950s to 0.194 in the 2010s (Fig. 2f and Supplementary Table 4). This was consistent with the decreasing trend of the genetic similarity index between the landraces and cultivars, with identity-by-state (IBS) decreasing continuously from 0.646 to 0.563 (Fig. 2g). Meanwhile, the IBS within the population kept increasing from early lines to current lines (Fig. 2h). Those results supported the notion that the cultivars tended to have decreased genetic diversity and homogenized genetic background probably because of wide germplasm exchange and common breeding practices. Furthermore, we used the XP-CLR statistic^15^ to detect genomic loci impacted by breeding selection, which was calculated by comparing each cultivar population from four ecological zones with the reference population of landraces (Fig. 1i and Supplementary Table 5). Genes involved in response to auxin or hormone, defense response to fungus and bacterium, and cell wall biogenesis were enriched among the top 1% genomic regions (250.0 million bases (Mb)) identified as being under selection in at least one zone (Fig. 1j and Supplementary Table 6). This revealed genomic improvements in wheat growth, morphogenesis, and disease defense in modern breeding. A total of 143.0 Mb genomic regions have been subjected to selection in at least two ecological zones (Supplementary Table 5), which represented the common targets of different breeding programs. For example, some genomic loci that are associated with important agronomic traits, such as *TaGW5*^16^, *FT3-B1*^17^, *Sh1*^18^ and *GA20ox-B1*^19–20^, were selected in four ecological regions. However, there are significant differences in selection targets among different ecological zones overall (Fig. 1k and Supplementary Fig. 2c). For example, a total of 65.5 Mb genomic regions were subjected to selection in both Zone 2 and Zone 4, but 60.0 MB and 74.0 Mb regions were under selection in Zone 2 and Zone 4, respectively, suggesting that the geography-specific trait improvements had shaped distinct genomic landscapes.

**Fig. 2.**
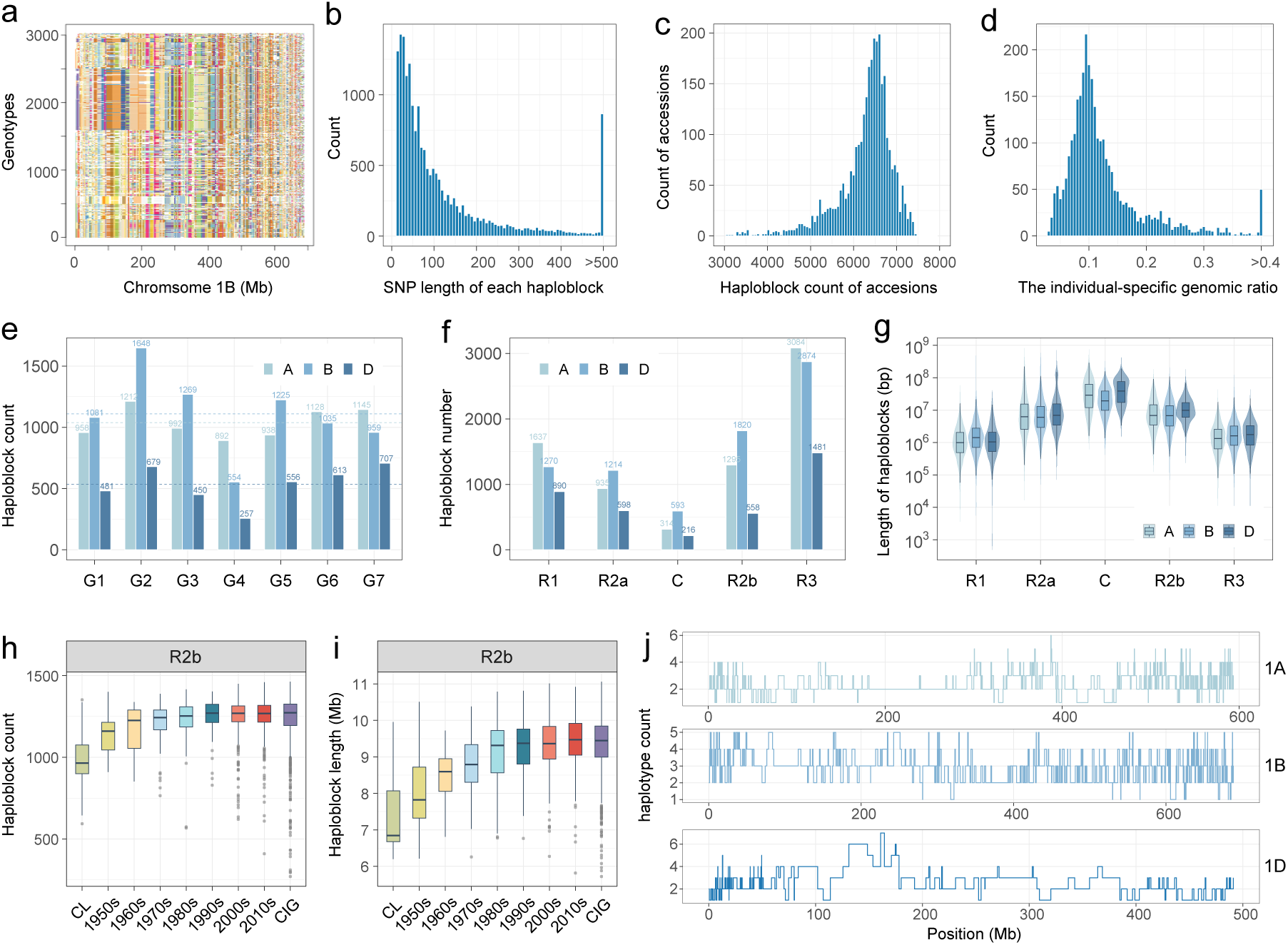
Characteristics of haplotype block library. **a,** Haplotype block (HB) plot of chromosome 1B across all accessions. The heatmap clearly highlights the short arm of chromosome 1R derived from rye. **b,** SNP number (bp) for each HB. **c,** Distributions of the number of HBs carried by each accession. **d,** Distributions of the genomic ratio not covered by HBs in each accession. The ratio is calculated as uncovered genomic region divided by reference genomic size. These uncovered regions correspond to individual-specific or rare haplotype segments with a frequency of less than 5. **e,** The number of HBs per chromosome. The dot lines represent the average HB numbers of A, B and D genome. **f,g,** Number (**f**) and length (Mb) (**g**) of HBs in five chromosomal regions of the A, B, and D genomes, respectively. R1 and R3 represent the terminal regions of the short arm and long arm of the chromosome, respectively; R2a and R2b represent the middle region of the short arm and long arm, respectively; C represents the centromere and its adjacent regions. The partitioning of these genomic regions is based on the previous study^31^. **h,i,** The number (**h**) and length (Mb) (**i**) of HBs carried by cultivars from different groups on region R2b. **j,** Number of HB along the chromosome. This result was calculated using HaploBlocker with the parameters ‘overlap_remove = TURE’.

### The genomic landscape defined by haplotype blocks

The selection unit during the breeding process is a chromosome segment rather than a single polymorphism site^21–24^. To better capture the genomic features shaped by breeding, we inferred the haplotype block (HB) from linked alleles in sub-population^25^. HB represents a genomic feature that manifests exclusively at the population level. We used HB as the unit in the following analysis to better dissect the genomic dynamics during breeding. As shown in Fig. 2a, HBs can be considered as genomic segments carried by each of the 3,030 wheat lines. A total of 991 wheat lines carried longer HBs due to the 1BL.1RS translocation^26^ (Fig. 2a), which has been the most widely deployed distant hybridization in China as well as in the world^27^. HB-based PCA showed consistent results with SNP-based PCA (Supplementary Fig. 3a), indicating that the inferred HBs were accurate. In total, 18,779 HBs were detected in this panel (Supplementary Data 1), with haplotype length ranging from 0.94 kilobases (kb) to 332.5 Mb with an average length of 2.57 Mb (Supplementary Table 7). Of those, 91.5% of HB was defined by more than 20 continuous SNPs with an average of 129.8 SNPs (Fig. 2b). A wheat line carried 6,263.4 HBs on average (Fig. 2c), covering 86.6% of the genome (Fig. 2d). Each HB was carried by at least 21 lines and at most 2,982 lines, with an average of 1,012.9 lines (Supplementary Fig. 3b). 95.7% of HBs was detected in more than 5% of lines (150) in the panel (Supplementary Fig. 3b). In other terms, HBs represent common chromosome segments in breeding but do not include those individual-specific or rare segments. Randomly sampling 12 accessions could approximately cover 90% of the entire HBs (Supplementary Fig. 3c). There were 7,265, 7,771, and 3,743 HBs for the A, B, and D subgenome, respectively (Fig. 2e). More HBs were detected near the arm end of the chromosome, while the fewest HBs were found in the centromeric regions (Fig. 2f and Supplementary Table 8). HBs located near the telomere are 2.3 Mb in length on average, while the average length of HBs in the centromere region was 37.0 Mb, with some exceeding 100 Mb (Fig. 2g and Supplementary Table 8). Compared to the cultivars, the landraces carried fewer and shorter HBs (Fig. 2 h,i and Supplementary Fig. 3d,e). Meanwhile, a significant increasing trend in both the number and length of HBs was observed from the 1950s to the 2010s, indicating that modern breeding has fixed favorable alleles into longer HBs. The average HB number in a sliding chromosome window was only 2.9 (ranging from 1 to 9) (Fig. 2j and Supplementary Table 9), suggesting that the HB diversity was limited in modern wheat. Those results supported that HB is a better unit for the study of the dynamic process of recombination and selection than SNP.

### Genomic composition of modern wheat cultivars

#### Landraces contributed the majority of genetic materials for modern cultivars

HB is defined as a series of linked identity-by-descent alleles and represents the chromosome segment that was selected in breeding^25^. Genomic regions covered by HBs in each line have been gradually increasing from 75.4% for landrace to 89.3 % for cultivars of the 2010s (Fig. 3a), suggesting modern breeding keeps fixing alleles into haplotype blocks. Conversely, genomic regions not covered by HBs indicate alleles not targeted by modern breeding (Fig. 3a), which highlighted the untapped genetic diversity among landraces. Our subsequent analysis focused on HBs rather than these genomic regions that have not been targeted by modern breeding. The concept of HB is naturally advantageous for tracking the ancestral donor population of chromosome segments. HB analysis revealed that a single landrace contributed between 52.1% and 92.2% of its genomic content to the modern cultivar population (Supplementary Table 10). For example, 91.4% of the genome for ‘Dazibai’ exists among the cultivar population, while 57.8% of the genome for ‘Banjiemang’. This high variability demonstrated the unequal genomic utilization of different landraces in breeding programs. For all the HBs carried by the cultivars, 93.2% (17,498/18,779) were preexisted in the landrace population, indicating that the majority of the genomic compositions of the cultivars were likely inherited from the landraces (Fig. 3b,c). Meanwhile, randomly sampling 42 accessions (64.6%) from landraces could approximately cover 90% of the entire HBs (Supplementary Fig. 4a). The magnificent proportion of modern HBs existing in old if not ancient germplasms inspired us to check the number of newly introduced HBs detected among germplasms released from the 1950s to 2010s. The result showed that a log_10_-scale decrease of new HBs was observed during the past six decades (Fig. 3b). Although our panel contains 2,348 lines released in the 21^st^ century, only 28 HBs were absent in germplasms released before 2000 (Fig. 3b), suggesting that breeding was primarily shuffling and assembling existing alleles rather than using new alleles.

**Fig. 3.**
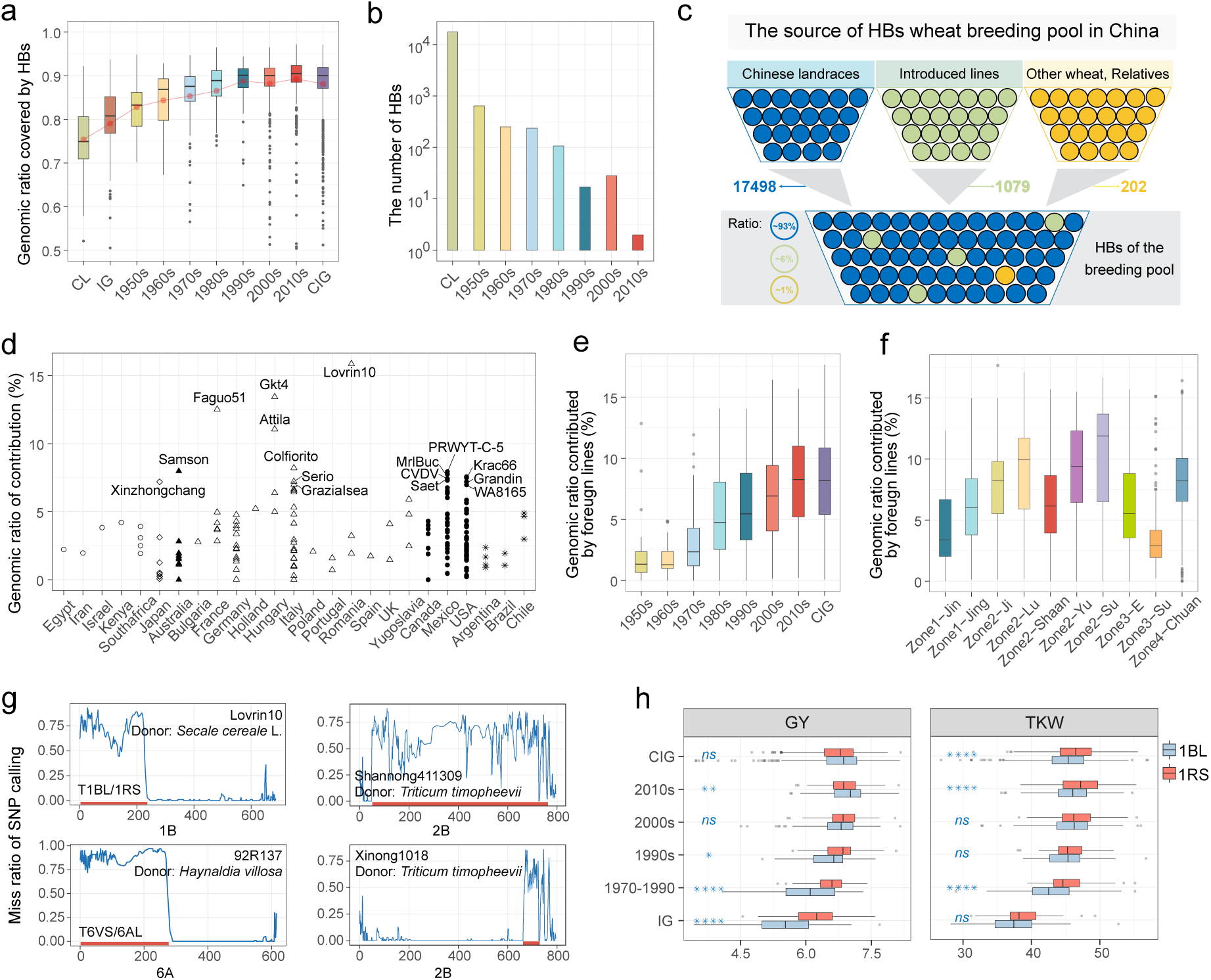
Quantitative analysis of genomic composition for modern Chinese cultivars. **a,** Genomic ratio covered by haplotype blocks (HBs) across different eras. **b,** Number of novel HBs in the haplotype pool across different eras. **c,** Schematic diagram depicting the sources of HBs of Chinese breeding pool. **d,** Ratio of HBs contributed to Chinese cultivars by each introduced line. The ratio was calculated using the total length of introduced-derived HBs divided by the IWGSC RefSeq v1.0 genome size. **e,f,** Ratio of HBs derived from introduced lines across historical stages (**e**) and different subzones (**f**). **g,** Four examples of alien chromosome translocations that were identified based on the ratio of SNP missing calls, including an 1BL.1RS translocation, a 6VS.6AL translocation and two wheat-*T. timopheevii* translocations. A continuous high missing ratio of SNPs indicates a putative alien segment. These translocations are relative to the positions in IWGSC RefSeq v1.0, rather than their own lengths. **h,** Boxplots comparing the grain yield (GY, t ha^−1^) and thousand grain weight (TKW) between the 991 T1B.1R lines and 2046 non-T1B.1R lines. Statistical significance was determined by a one-tail t-test (ns: non-significant; ^⁎^*P*<0.05; ^⁎⁎^*P*<0.01; ^⁎⁎⁎^*P*<0.001; ^⁎⁎⁎⁎^*P*<0.0001).

#### Introduced germplasms contributed 5.7% new HBs to the cultivar population

The introduced lines are believed to have historically played a significant role in improving the yield and quality of Chinese wheat cultivars^6–7,28^. For the 6.8% HBs (1,281/18,779) that were not detected in the landraces, 5.7% (1,079/18,779) were detected in introduced lines, indicating that they are likely inherited from these lines and therefore categorized as introduced-derived HBs (Fig. 3c and Supplementary Fig. 4b). Among the 1,079 HBs, 518 (48.0%) were from at least 10 introduced lines while 101 HBs (9.4%) were from only a single introduced line (Supplementary Fig. 4c). On average, each introduced line contributed 3.4% of its genome to the cultivar population. (Fig. 3d and Supplementary Table 10). Among the 193 introduced lines, 27 lines had contributed less than 1% of their genomes. For example, ‘Nanda2419’ and ‘Early Piemium’ commonly used by breeders^6–7^ contributed very little HBs, less than 1% of their genomes. Conversely, ‘Lovrin10’ has the highest ratio (15.9%) of introduced-derived HBs as it is the source of the T1B.1R lines in China, which is consistent with the historical role of this famous line^6–7,27^. Those results revealed the genomic footprints of introduced germplasms in the Chinese wheat improvement.

The proportion of the genome covered by introduced-derived HBs in each Chinese cultivar rose from 2.3% in the 1950s to 8.1% in the 2010s (Fig. 3e), indicating that alleles from introduced lines were continuously favored by breeders (Supplementary Fig. 4d). Strong geographical bias was observed for the frequency of introduced-derived HBs among different breeding programs (Fig. 3f and Supplementary Fig. 4e). For example, Zone2-Lu and Zone2-Yu, which are two neighboring provinces, displayed differential utilization for introduced-derived HBs. Zone4-Chuan, characterized by climate and pest risk profiles that differ significantly from other ecological zones, exhibited a unique frequency spectrum of introduced-derived HBs. Those geographical biases demonstrated the differential breeding usage of introduced-derived alleles in different breeding programs.

#### Genomic contribution from wild relatives

Some ultralong chunks, such as the HBs on 1B and 6A, might be wild relative introgressions, as their recombination tends to be suppressed (Supplementary Fig. 4f). The HB-based heatmap characterized the chromosome segments originating from 1RS of rye (T1BL.1RS^26^), two types of durum wheat 1B^29^, 6VS of *Haynaldia villosa* (T6VS.6AL^30^). Additionally, about 1.1% of the HBs (202) among the cultivars that could not be detected in either the landraces or introduced lines were likely originated from wild relatives (Fig. 3c). However, it is difficult to infer extra unreported alien segments by HB method, especially many alien segments are low frequency and cannot be captured by HBs. Alien translocations tend to result in continuous and high but not all missing genotype calls due to sequence divergence between wild relative and the hexaploid wheat when the reference genome (IWGSC RefSeq v1.0^31^) was used for genotyping^32–33^. Therefore, we used the SNP missing ratio to infer the genomic segments from wild relatives to complement the HB-based method (see Methods). For example, the SNP missing ratio was average of 0.75 on 1BS for ‘Lovrin10’, which donated the widely used 1BL.1RS translocation^26^ (Fig. 3g). The 6VS.6AL translocation^30^ was also uncovered for ‘92R137’ line^34^ based on continuous and high missing genotype calls on 6AS (Fig. 3g). In total, we identified 276 putative alien translocations with a length at last 10 Mb, including genomic segments from rye, *H. villosa*, *Triticum timopheevii*, *Aegilops markgrafii*, and *Aegilops ventricose* (Supplementary Table 11).

An ultralong introgression came from *T. timopheevii* and was approximately located on 79-761 Mb of 2B^35^ (Fig. 3g), which was carried by 11 lines, including three innovative germplasms and eight introduced lines (Supplementary Table 11). Another *T. timopheevii* introgression carried by 154 lines was also discovered on approximately 664-749 Mb of 2B^36^ (Fig. 3g). This alien segment was detected in seven introduced lines and 147 cultivars released after 1970 (Supplementary Table 11). Those two introgressions might represent the utilization of wild relatives for disease resistance in breeding^37^. The most frequent introgression event comes from *Ae. markgrafii* and was located on approximately 592-623 Mb of 2D^38^ (Supplementary Table 11), which was carried by more than one-third in this panel (1,185 lines), including 86 introduced lines, 1095 Chinese cultivars, and four landraces, suggesting that this alien segment was an ancient translocation. Among all introgressions, the short arm of 1R from rye genome was the most famous. In our panel, 32.7% of lines (991) carried this translocation (Fig. 3g and Supplementary Table 11). Previous studies showed that the translocation improved disease resistance and yield^39–43^. However, phenotypic analysis revealed that, although the thousand kernel weight (TKW) of T1BL.1RS lines was significantly higher, the grain yield did not increase for lines released after 2000, and even significantly decreased in lines released during the 2010s (Fig. 3h). This result suggested that the yield improvement advantage of 1BL.1RS translocation tended to disappear after 30 years of utilization.

Unlike traditional views that introduced germplasms greatly contributed novel favorable alleles, our HB-based analysis highlighted the dominant role of local landraces in the genomic makeup of modern cultivars. Our results estimated that the introduced lines only contributed 5.7% of the total HBs of the wheat breeding pool. Those genetic segments are widely distributed across the wheat genome with strong geographical bias, demonstrating distinct breeding preferences of different ecological environments. Meanwhile, we also revealed the extensive use of many alien segments from wild relatives in modern breeding, and some alien segments that are not widely used but may have potential in breeding.

### The genetic basis of trait improvement from landraces to modern cultivars

#### Agronomic traits and narrow-sense heritability

As the most important trait in cereal crops, grain yield is highly influenced by environment^44–46^. To accurately assess the genetic traits of wheat varieties, the 3,030 wheat lines were tested in field trials of five different ecological sites in China. A bunch of 14 traits, including yield and yield component traits, were collected over two consecutive years (Supplementary Fig. 5). In this study, we investigated the yield per plot for each accession through large-scale field trials, ensuring compliance with the standard definition of yield^5^. Significant increases in heritability (*h*²) were observed when the best linear unbiased prediction (BLUP) was used to fit multi-batch phenotypes to remove environmental impact (Supplementary Fig. 6a,b and Supplementary Table 12). The yield heritability was less than 0.4 in a single environment, but it increased to 0.58 when BLUP values were used. Most traits, including lodging resistance, heading date, and spike length, exhibited high heritability (even above 0.7), whereas effective tiller number and cold resistance showed the lowest heritability (below 0.4). The difference in heritability between BLUP and single-environment phenotypes indicated that traits such as flowering date, plant height, lodging resistance, spike length, and TKW were more environmentally stable compared to grain number per spike, kernel number per spikelet, spikelet number per spike, and grain yield. These results demonstrated the robustness of the field trial design in investigating the genetic basis of trait improvement.

#### The routes of yield improvement

The average grain yield has increased from 5.025 ± 0.516 t ha^−1^ in landraces to 6.945 ± 0.428 t ha^−1^ in the 2010s cultivars, indicating that the yield increased by about 37.6% through genetic improvement (Fig. 4a and Supplementary Table 13). The most significant yield increases (Δ_GY_) were observed in wheat lines from the 1950s-1960s (Δ_GY_ = 0.762 t ha^−1^), followed by the 1970s-1980s (Δ_GY_ = 0.279 t ha^−1^) and the 1980s-1990s (Δ_GY_ = 0.284 t ha^−1^), coinciding with the deployment of semi-dwarf genes^6,8–9^ and the utilization of distant hybrids^27,30^. However, the rate of yield increase slowed after the 1990s (Δ_GY=_ 0.174 t ha^−1^), indicating that the yield potential has been constrained due to genetic homogenization of cultivars. Similarly, TKW exhibited the highest increase during the1950s-1960s (Δ_TKW_ = 4.3 g) and 1970s-1980s (Δ_TKW_ = 2.6 g), followed by a recent slowdown (Fig. 4a and Supplementary Table 13). Overall, breeding improvements have significantly contributed to wheat with larger and more grains, shorter height, fewer single-plant tillers, and earlier flowering (Fig. 4a and Supplementary Fig. 7a). From the spatial perspective, significant differences in agronomic traits were observed among cultivars from different ecological zones or subzones (Supplementary Fig. 7b). For instance, cultivars from Zone 1 exhibited the greatest effective tiller number, the tallest plant height, the fewest spikelet number per spike, the strongest cold tolerance, and the latest flowering time. Cultivars from Zone 2 had the highest TKW, Zone 3 showed the longest spike length, and Zone 4 displayed the greatest grain number per spike, spikelet number per spike and kernel number per spikelet, and the latest maturity date. These significant trait differences reflect the regionalized breeding strategies implemented in China.

**Fig. 4.**
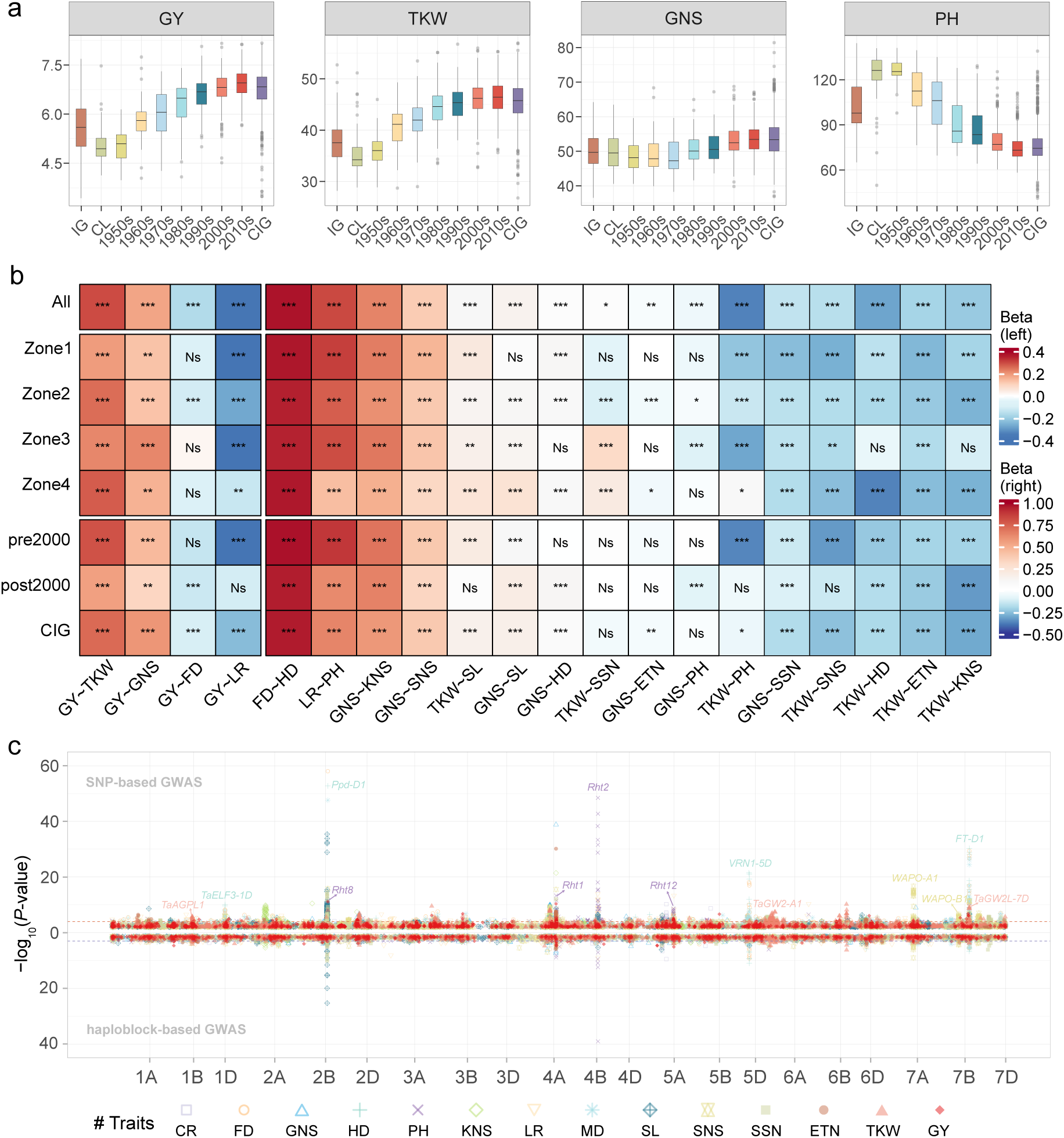
The routes and genetic basis of agronomic trait improvement. **a,** Comparison of agronomic traits of wheat lines across different historical stages. GY: grain yield (t ha^−1^), TKW: thousand grain weight. GNS: grain number per spike, PH: plant height. **b,** Quantification of yield component effects on yield across different ecological zones or historical stages using structural equation models. The values of each row are comparable, but the values of each column are not. The color represents the standardized partial regression coefficients (*β*). Statistical significance is denoted as follows: Ns: non-significant; ^⁎^*P*<0.05; ^⁎⁎^*P*<0.01; ^⁎⁎⁎^*P*<0.001. **c,** Manhattan plot of GWAS of 14 traits. The upper panel represents SNP-based GWAS, and the lower panel shows HB-based GWAS. For SNP-based GWAS, the multiple-test corrected significance threshold is denoted with red horizontal dashed lines. For HB-based GWAS, the significance threshold is considered to be 1×10^−3^. Some known genes are labeled in corresponding QTLs. CR: cold resistance; ETN: effective tiller number; FD: flowering date; HD: heading date; KNS: kernel number per spikelet; LR: lodging resistance; MD: maturation date; SL: spike length; SNS: spikelet number per spike; SSN: sterile spikelet number per spike.

Yield increase can be achieved through the increase of different yield components, as well as the cooperative improvement of traits^47^. Significant high correlations were observed between yield and lodging resistance (|PCC| = 0.62), plant height (|PCC| = 0.55), TKW (|PCC| = 0.53), and flowering date (|PCC| = 0.45) (Supplementary Fig. 6c). To elucidate breeding strategies from different historical periods and ecological zones, we employed a formal structural equation model^48^ (SEM) to quantify the direct and indirect effects of yield components on yield (Supplementary Note 1). Based on the collected trait dataset, SEM analysis across all Chinese landraces and cultivars revealed that wheat yield improvement in China has primarily relied on enhancing lodging resistance (|*β* _LR_| = 0.39), followed by increasing TKW, grain number per spike, and earlier flowering time (|*β* _TKW_| = 0.30, |*β* _GNS_| = 0.18, |*β* _FD_| = 0.17) (Fig. 4b and Supplementary Table 14). However, the approaches for yield improvement have shifted significantly over different historical periods. For example, yield improvement was predominantly driven by enhanced lodging resistance during the 1950s–1990s, while increasing TKW became the dominant contributor to yield improvement after 2000. This reflects the breeding history: the ‘Green Revolution’ in the 1960s initiated the extensive use of semi-dwarf genes^11^, which markedly enhanced lodging resistance, thereby minimizing yield losses to increase substantially yield. Nearly all cultivars released after 2000 have achieved lodging resistance (Supplementary Fig. 7a), owing to the widespread deployment of semi-dwarf and lodging-resistant genes^6,8–9^. As a result, optimizing spike traits, such as TKW and grain number per spike, has become essential for further improvements (Fig. 4a and Supplementary Fig. 7a). Moreover, TKW, grain number per spike, and lodging resistance showed roughly equivalent effects on yield improvement in the advanced breeding lines (Supplementary Table 14).

From the spatial perspective, the routes to achieving yield improvement exhibited distinct patterns across different ecological zones. For instance, enhancing lodging resistance had the most substantial positive effect on yield improvement in Zone 1 and Zone 3 (Fig. 4b and Supplementary Table 14). In Zone 2, yield increases were predominantly driven by enhanced lodging resistance and increased TKW, with both traits contributing almost equally to the observed improvements (|*β* _LR_| = 0.28, |*β* _TKW_| = 0.25). Notably, in Zone 4, yield gains were mainly driven by improvements in TKW and grain number per spike (|*β* _TKW_| = 0.27, |*β* _GNS_| = 0.16), while lodging resistance played a relatively minor role in yield enhancement (|*β* _LR_| = 0.13). In the southwestern wheat zone, particularly Sichuan, where the climate is warm with limited sunshine, and winds during the heading to maturity stage are weaker and less frequent that in other zones^6^, enhancing lodging resistance does not directly contribute to more yield gains. In summary, these differences in trait improvements reflected the ecology-specific breeding routes tailored by Chinese breeders to develop specialized cultivars for distinct ecological environments. The SEM analysis revealed the breeding routes of wheat across different historical stages and ecological zones, providing valuable insights for future adjustments in breeding strategies.

#### Identifying QTLs and HBs associated with traits through GWAS

To uncover the genetic basis of trait evolution, we performed GWAS analysis to identify genomic loci and HBs associated with 14 agronomic traits. In total, 234 quantitative trait loci (QTLs) (*P* < 10^−5^) were identified through GWAS, including 10 QTLs associated with grain yield (Fig. 4c, Supplementary Fig. 8, and Supplementary Table 15). The highest number of QTLs were identified for flowering date (31), whereas the lowest number were mapped for cold tolerance (4) (Supplementary Fig. 9a). The total phenotypic variance explained (PVE) by the QTLs mapped by GWAS for each trait ranged from 3.9% to 38.9%, with 9.4% for grain yield (Supplementary Fig. 9b). The PVE by the ten QTLs of yield ranged from 0.6% to 1.5% (Supplementary Table 15). Only 12 QTLs explained more than 3% of the phenotypic variance, while 159 QTLs explained less than 1%, reflecting the polygenic nature of quantitative traits. Two QTLs (*QTL-AX-89703298* and *QTL-AX-111956072*) colocalized with *Rht2* and *Ppd-D1* explained the highest phenotypic variation for plant height (PVE = 8.4%) and flowering date (PVE = 7.9%), respectively. Many QTLs were colocalized with reported genes or homologous genes of known function. For examples, four semi-dwarf genes (*Rht1, Rht2*, *Rht8* and *Rht12*), seven flowering or heading genes (*Ppd-D1*, *TaELF3-1D*, *TaTFL2*, *PhyC*, *VRN1-5D*, *VRN2-5A* and *VRN3-7D*), four grain weight genes (*TaAGPL1*, *TGW1*, *TaGW2-A1* and *TaGW2L-7D*), and six genes associated with grain number or spikelet (*TaTB1, WAPO-A1*, *WAPO-B1*, *TaSPL14*, *SVP-B1* and *TaLOG*) were confirmed (Supplementary Fig. 8 and Supplementary Table 15).

Many genomic loci are associated with multiple traits, as a result of genetic linkage or pleiotropy^49^. For example, *QTL*-*AX-111956072* was associated with yield and eleven yield components, while *QTL-AX-89323611* was associated with yield and seven yield components (Supplementary Table 16). These two loci are located near *Ppd-D1* and *Rht1*, respectively. To better reveal linked gene clusters or pleiotropic haplotypes associated with traits, we performed HB-based GWAS and identified 522 HBs associated with 14 traits (*P* < 10^−3^), including 35 HBs associated with grain yield (Fig. 4c and Supplementary Table 17). Among these 35 HBs, 19 HBs exhibited yield-enhancing effects. GWAS showed that 88 HBs were significantly associated with at least two traits (Supplementary Fig. 10 and Supplementary Table 18). For example, four HBs associated with reduced plant height (4B-HB117[*Rht1*] and 4B-HB119) and early flowering or maturing (2D-HB93[*Ppd-D1*] and 7B-HB443) showed yield-increasing effect. Two HBs (4A-HB371 and 7A-HB857) showed positive effects on spikelet number per spike, but significantly negative effects on TKW. Two HBs (2D-HB368 and 4B-HB116) could increase the grain number per spike, but reduce TKW and effective tiller number, respectively. Because these HBs have multi-effects on different traits, the selection in breeding may encounter trade-offs between traits. Uncoupling these trade-offs and making reasonable use of the multi-effect haplotype segments may help further enhance yield^50^.

Frequency dynamic analysis can reveal the utilization of HBs by breeders, allowing us to review breeding history at the genomic level and broaden our understanding of the selection during breeding^51^. The frequency analysis showed that eleven HBs with yield-enhancing effects, such as 2A-HB102, 4B-HB190, and 5A-HB304, had low frequencies in landraces but significantly increased frequencies in cultivars, indicating they have undergone artificial selection (Supplementary Fig. 11 and Supplementary Table 17). In contrast, four yield-enhancing HBs, such as 1B-HB410 and 5B-HB205, were found at high frequencies in landraces but were not adequately utilized in cultivars. Among all yield-enhancing HBs, 2D-HB93, carrying *Ppd-D1*, was the most widely utilized in breeding. In cultivars released in the 2010s, 89% of lines carried this HB. Conversely, 2B-HB1352 and 3B-HB1025 have yet to be widely utilized in breeding, indicating that they were overlooked in breeding programs and could serve as valuable genetic resources for future improvement. Some HBs carrying *Rht* genes, such as 2D-HB85[*Rht8*], 4B-HB113[*Rht1*], 4D-HB37[*Rht2*], and 5A-HB909[*Rht12*], showed significantly increased frequencies during breeding. Among these, *Rht1*, *Rht8*, and *Rht12* were widely employed before the 1960s, while *Rht2* was extensively utilized after the 1960s, revealing the history of dwarfing breeding in Chinese wheat. Strong selection pressure has targeted other agronomic trait-associated HBs, such as 1B-HB977 and 5B-HB800, which increased TKW; 7A-HB447, which increased effective tiller number; and 2D-HB85, which reduced sterile spikelet number per spike (Supplementary Fig. 11 and Supplementary Table 17), highlighting their importance in breeding.

From the spatial perspective, the utilization of many HBs exhibited significant geographic biases. For example, two yield-enhancing HBs (5B-HB205 and 7B-HB443) have been widely utilized in northern China (Zone 1 and Zone 2) but are less frequently used in southern China (Zone 3 and Zone 4) (Supplementary Fig. 11). On chromosome 5A, a major QTL associated with cold tolerance spans the 512– 524 Mb region (Supplementary Fig. 8) and contains more than 20 *C-repeat binding transcription factor*^52^ (CBF) homologous genes. This QTL includes six cold-resistance HBs whose frequency distributions also displayed distinct north-south differences; they were extensively utilized in Zone 1 and Zone 2 (except Zone 2-Shaan) (Supplementary Fig. 11), where ensuring winter wheat survival is a critical breeding objective^6^. Similarly, five flowering-related HBs exhibited the same north-south differentiation (Supplementary Fig. 11). For instance, 5D-HB334[*VRN1-5D*], 6A-HB760, and 6D-HB358 were widely utilized in southern zones (Zone 3 and Zone 4), whereas their frequencies were significantly lower in northern populations. Additionally, some HBs associated with spike traits also showed geographic specificity (Supplementary Fig. 11). For example, 5D-HB80, 5A-HB793, and 6D-HB237, which increased grain number per spike, have been strongly selected in Zone 2, Zone 3, and Zone 4, respectively. These geographically specific frequency distributions result from long-term natural selection and ecologically specific breeding targets.

In summary, frequency dynamics analysis revealed the spatiotemporal specificity of the selection on these trait-associated HBs, providing a scientific basis for understanding the historical trajectory of agronomic trait selection in modern breeding. Many HBs have gradually been fixed during breeding due to common breeding targets. At the same time, we have identified numerous HBs that have yet to be widely utilized in breeding, which could be prioritized in future breeding programs to prevent the loss of genetic diversity.

### Deciphering epistatic network modules to improve wheat yield potential

#### Epistasis and its association with yield

Over 90% of the genomes in the modern cultivars are composed of pre-existing HBs in the landraces (Fig. 3a-c), indicating that past breeding efforts mainly involved the diverse assembling of HBs already present in the breeding pool. It is widely accepted that wheat lines carrying more favorable haplotypes will have significantly higher yields, a principle supported by some breeders. However, the correlation between yield and the number of favorable HBs was only 0.29 (Fig. 5a), suggesting that yield improvement involves more complex genetic mechanisms and cannot be attributed solely to the aggregation of favorable alleles. Only 10 QTLs and 35 HBs were identified to have significant associations with grain yield through GWAS, explaining a total of 9.4% of the phenotypic variation in yield, with each QTL explaining only 0.6%-1.5% (Supplementary Fig. 9). Furthermore, the heritability of yield, estimated based on all HBs or SNPs, was only 0.56 or 0.58, respectively. These limitations prompted us to focus on epistasis—the phenotypic effects of genetic interactions across two or multiple loci^53–58^. While the effect of a single allele on complex traits may be minor and non-significant, these alleles can have a significant impact on traits through their epistatic interactions^56^. Together, these alleles may form a genetic epistatic network that regulates yield and other complex traits^59–60^.

**Fig. 5.**
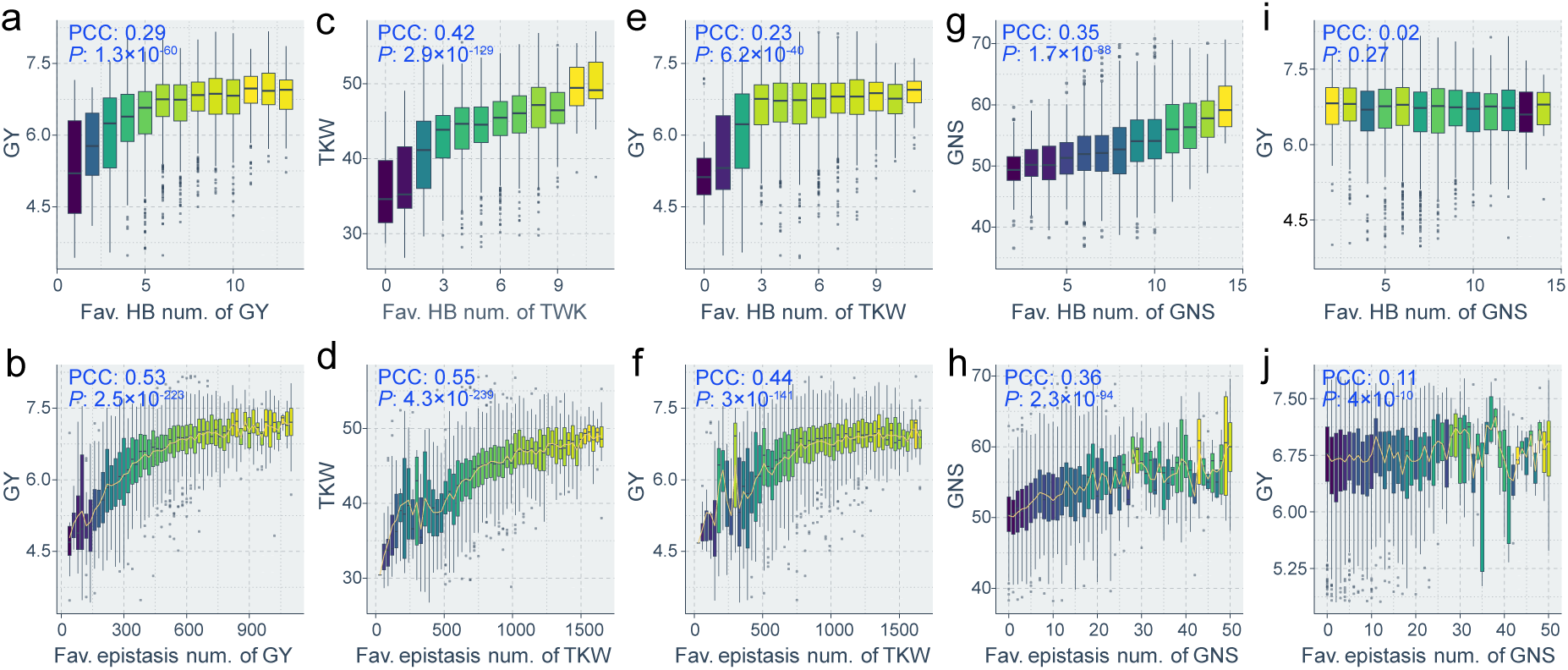
Pearson correlation coefficients between grain yield and favorable HB or favorable epistasis. **a**, Correlations between grain yield and the number of favorable HB of yield. **b**, Correlations between grain yield and the number of favorable epistasis of yield. **c**, Correlations between TKW and the number of favorable HB of TKW. **d**, Correlations between TKW and the number of favorable epistasis of TKW. **e**, Correlations between grain yield and the number of favorable HB of TKW. **f**, Correlations between grain yield and the number of favorable epistasis of TKW. **g**, Correlations between GNS and the number of favorable HB of GNS. **h**, Correlations between GNS and the number of favorable epistasis of GNS. **i**, Correlations between grain yield and the number of favorable HB of GNS. **j**, Correlations between grain yield and the number of favorable epistasis of GNS. The *P*-values were provided. Each point represents a sample, and boxplots are plotted based on bins of HB number or epistasis number. The yellow line represents the variation in the average value of each bin. The median of each bin is used to color the boxes in the boxplot.

In statistical genetics, epistasis is the deviation of the genotypic value of two or more loci from the sum of the genotypic values contributed by each locus individually, defined as the departure from additive effects in the linear model^53,55–56,58^ (Supplementary Note 2). To elucidate the role of epistasis in yield improvement, we conducted an exhaustive two-dimensional epistatic detection (HB-HB interactions)^61^. A total of 13,119 HB-HB interactions were found to significantly affect grain yield, 23,935 interactions affected TKW, and 1,161 interactions affected grain number per spike (*P* < 10^−16^) (Supplementary Fig. 12a, Supplementary Data 2 and Supplementary Note 3). Notably, 96.8% of the HBs (4495/4643) showed no significant associations with yield or yield component traits, but they significantly impacted yield through epistasis, which highlights the complex polygenic nature of yield and the indirect effects of numerous small-effect loci. Based on the effect on traits, these significant HB-HB interactions were categorized into favorable, inferior, or moderate epistasis (Supplementary Fig. 12b, see Methods). For yield-associated epistasis, 27.7% (3,636/13,119), 21.2% (2,782/13,119), and 51.2% (6,721/13,119) HB-HB interactions were classified as favorable, inferior, and moderate epistasis, respectively (Supplementary Table 19). By comparing the correlation between the number of favorable HBs or epistasis carried by each line and its yield, it was found that the correlation between epistasis and yield was significantly stronger than the correlation between HBs and yield (PCC: 0.53 vs. 0.29) (Fig. 5a, b). Furthermore, the heritability was calculated to assess the robustness of the observed correlations. The results showed that the heritability for yield based on SNPs or HBs was only 0.53-0.63, but the genome-wide epistasis explained significantly higher phenotypic variation (*h*² = 0.70) (Supplementary Table 20). These results suggest a closer link between yield and epistasis.

Similarly, the analysis of both TKW and grain number per spike revealed the same trend, showing significantly stronger correlations with epistasis than with HBs (PCC_TKW_: 0.55 vs. 0.42; PCC_GNS_: 0.36 vs. 0.35) (Fig. 5c,d,g,h). In breeding, yield can be improved by targeting different yield components. To explore this, we examined the relationship between grain yield and the number of favorable HBs that were significantly associated with TKW or grain number per spike. The results showed a relatively low correlation between yield and the number of favorable HBs associated with TKW (PCC = 0.23) and no correlation between yield and the number of favorable HBs associated with grain number per spike (PCC = 0.02) (Fig. 5e, i). However, when epistasis was taken into account, the correlations became significantly stronger (Fig. 5f, j), especially the correlation between yield and the number of favorable epistasis associated with TKW, which reached 0.44 (Δ_PCC_ = 0.21), indicating that epistasis associated with TKW plays a key role in driving yield improvement in China.

#### Remodeling and optimization of epistatic network modules in wheat breeding

The improvement of complex polygenic traits may be driven by higher-order and complex genetic networks and modules composed of these epistases. To explore the relationship between yield and epistasis at the network level, we used a network graph to visualize these epistatic interactions. The network graph revealed a tightly connected network and a series of modules (Supplementary Fig. 13a), which might reflect complex biological processes. To reveal the dynamic changes of epistatic networks during modern breeding, we next analyzed the networks at individual and population levels.

The trends across different historical stages revealed that the number of favorable HBs carried by wheat lines has shown only a slight increase over time (Supplementary Fig. 13d), whereas the number of favorable epistasis carried by these lines has increased continuously and significantly (Supplementary Fig. 13e). For example, the epistatic networks of landraces (e.g., ‘Chengduguangtou’) and early improved lines (e.g., ‘Bima1’) were not only smaller in scale but also loosely connected (Supplementary Fig. 14). In contrast, the epistatic networks of current elite cultivars (e.g., ‘Chuanmai42’ and ‘Jimai22’) expanded significantly, forming larger and more complex modules, suggesting that yield improvement may be driven by remodeling and expanding the epistatic networks. Given the distinct trends in the epistatic networks of wheat lines with different improvement statuses, we further analyzed correlations between grain yield and network topological properties. The results showed that yield was significantly correlated with many topological parameters, including the number of nodes, average neighbor count, graph diameter, maximum component size, average module size, and number of modules (Fig. 6a, Supplementary Fig. 15, and Supplementary Table 21). Among these, the correlation between yield and the number of modules was the strongest (PCC = 0.58). The topological properties, such as the number and size of modules carried by wheat lines, have increased significantly from landraces to current cultivars (Fig. 6b and Supplementary Fig. 16). On average, each landrace carried three modules, while cultivars released after 1990 carried an average of 12 modules. In addition, we analyzed the correlations between different network parameters to characterize the patterns of network changes. The results indicated that as the number of nodes or edges in the network expanded, both the number and size of modules also increased significantly (Supplementary Fig. 17). This suggested that the epistatic network changes driven by selection in breeding are governed by certain regularities, where the aggregation of epistasis involves continuous expansion and refinement of modules rather than random accumulation of unrelated epistasis.

**Fig. 6.**
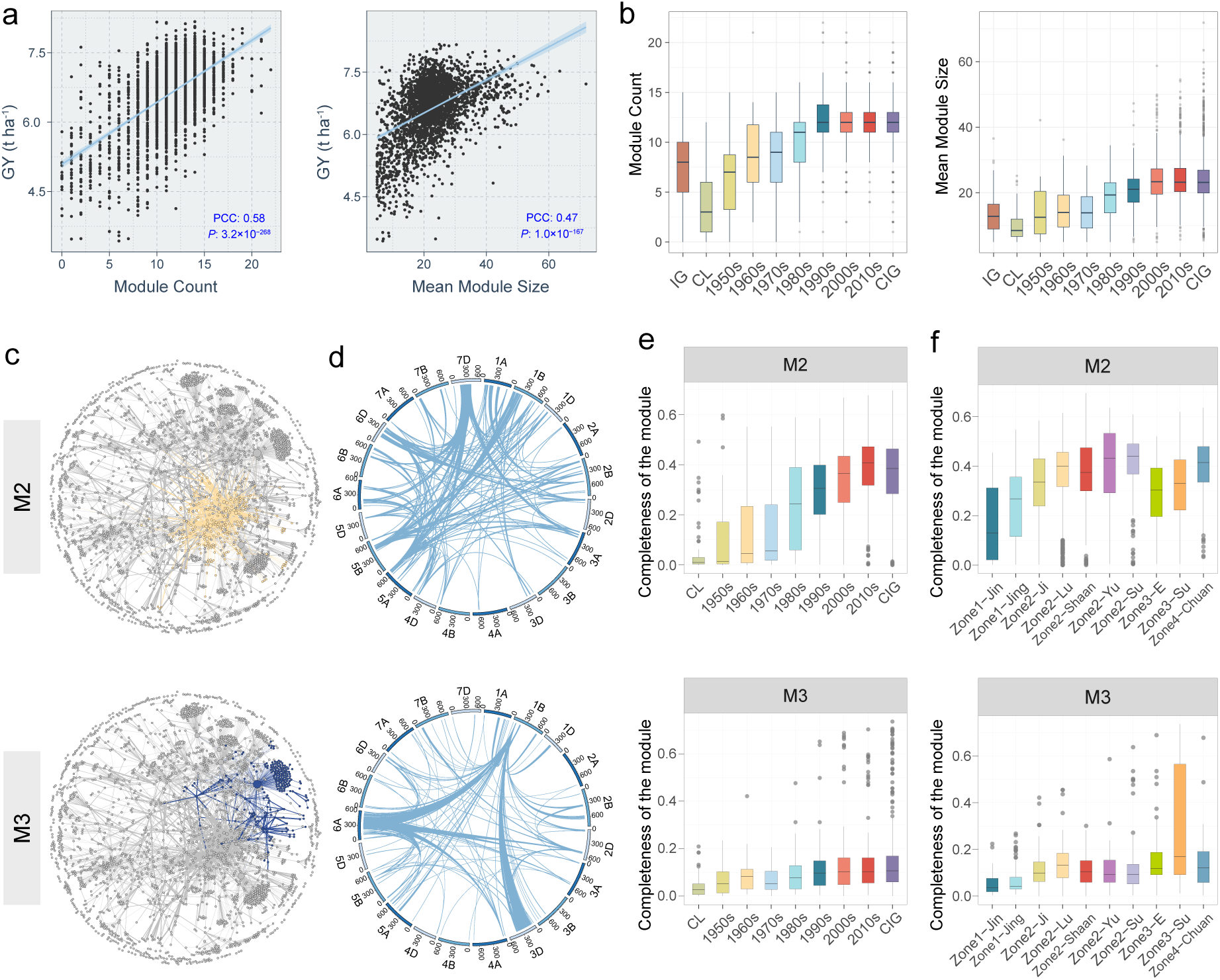
Remodeling and optimizing of epistatic network modules. **a,** Pearson correlation coefficients (PCC) between the module number and size of epistatic network and grain yield. Each point represents a sample. The *P*-value of correlation was calculated. A linear method was used for fitting. **b,** The module number and size of wheat lines across different era groups. Each point represents a sample, and boxplots are plotted based on groupings. **c,** Visualization of favorable epistatic module M2 (yellow) and Module M3 (dark blue). Each point represents an HB, and each edge indicates a favorable epistatic interaction between two HBs. Modules M2 and M3 are highlighted in yellow and dark blue, respectively. **d**, Circular link plots of modules M2 and M3 on genome. This plot reflects the genomic locations of these epistasis and modules. Epistasis between HBs is represented by links. Since an HB corresponds to a genomic segment, all links are shown with a width. **e**, The completeness of modules M2 and M3 changes over time. Each point represents a sample, and boxplots are plotted based on era groupings. The completeness of each module is calculated by dividing the number of edges in the module carried by each wheat line by the total number of edges in the module. **f**, Comparison of module completeness of wheat lines across different subzones. The calculation of module completeness is the same as that for **e**. Each point represents a sample, and boxplots are plotted based on subzone groupings.

The epistatic network patterns of wheat populations from different ecological zones or eras showed significant differences. For example, wheat populations from different ecological zones exhibited significant differences in utilization frequency of epistasis (Supplementary Fig. 13b and Supplementary Fig. 18), which may be linked to breeding targets and environmental selection pressures during long-term breeding processes. On the other hand, the frequency of epistasis significantly increased from the landrace population to the 2010s cultivar population (Supplementary Fig. 13c and Supplementary Fig. 19), suggesting that the selection in breeding primarily focused on numerous epistatic interactions, rather than just a few alleles or haplotypes. Furthermore, we analyzed the dynamic changes of 15 major modules over time (Fig. 6c-e, Supplementary Fig. 13a and Supplementary Fig. 20-21). For instance, the completeness of modules M2 and M3 significantly increased from the landrace population to the 2010s cultivar population, reflecting the ongoing optimization and refinement of these modules in the breeding process. In addition, the extent of module optimization and utilization differed significantly across populations of ecological zones (Fig. 6f and Supplementary Fig. 22). For example, module M2 showed the highest level of optimization and utilization in populations of Zone 2 and Zone 4, while module M3 was most refined and utilized in populations of Zone 3. The dynamic changes of modules uncovered the evolutionary process of the genome and epistatic network during breeding.

#### Inheritance and formation of favorable epistatic modules in breeding lineages

To test the transmission and forming of favorable epistatic modules during yield improvement, we analyzed the completeness of modules associated with yield in specific breeding lineages. For example, the high-yielding cultivar ‘Chuanmai 104’ was developed through the hybridization between ‘Chuanmai 42’ and ‘Chuannong16’. The module analysis of this lineage showed that ‘Chuanmai 104’ inherited modules M4, M6, M7, and M10 from ‘Chuanmai 42’, module M12 from ‘Chuannong16’, and created modules M2, M11, and M14 (Supplementary Fig. 23). Although hybridization led to significant genomic changes in the progeny, the modules remaining stable after recombination indicated that favorable modules were preserved by breeders. The second representative case is the ‘Bimamai’ lineage. In the 1940s, six lines were bred by hybridizing ‘Quality’ from the United States with the landrace ‘Mazhamai’^6^. The analysis of this lineage revealed that the progeny line, ‘Bima1’, expanded modules M7, M8, M10, M14, and M15, while ‘Bima4’ inherited M11 and M15 from ‘Quality’ and expanded M7, M10, and M14 (Supplementary Fig. 24). As a matter of fact, ‘Bima1’ was the most widely cultivated variety in the 1950s and ‘Bima4’ became a backbone breeding parental line^6^. Another representative case is the development of ‘Chang 6878’ through the hybridization between ‘Linhan 5175’ and ‘Jinmai 63’ (Supplementary Fig. 25). The parent ‘Linhan 5175’ exhibited a high level of completeness in module M5, and the progeny ‘Chang 6878’ inherited this superior module. ‘Linhan 5175’ is an innovative germplasm, which reflects its potential value as an excellent parent for breeding. Furthermore, module analysis of a series of other lineages revealed the same patterns (Supplementary Fig. 26-29 and Supplementary Note 4), suggesting that the effective transmission, integration, and formation of epistatic modules during breeding play a crucial role in wheat yield improvement.

In summary, these findings indicated that modern breeding has significantly remolded, expanded, and optimized epistatic networks and modules. A series of modules remained stable during breeding, suggesting that breeding targeted the entire module rather than individual or a few alleles or haplotypes. Moreover, new favorable epistatic modules can arise, indicating that modules can not only be inherited but also reshaped and refined. These modules likely represent specific biological processes underlying yield and its component traits, meaning that genetic improvement essentially involves reshaping, expanding, and optimizing these biological processes. This highlights that future wheat design breeding should be based on genetic modules rather than focusing solely on one or a few alleles.

## Discussion

### A representative germplasm panel and precise identification of yield

Previous population genomic studies on wheat yield were typically based on recombinant inbred lines^62^ or natural populations, such as diverse germplasms, landraces, and modern cultivars^28,63–66^. Although these studies identified some yield-related QTLs, they were not suitable for deciphering the process of sustained yield improvement. Unlike previous research, our wheat population represents a 70-year history of yield improvement, making it highly suitable for analyzing genomic changes during breeding and the genetic mechanisms underlying yield enhancement. These germplasms include main cultivated varieties from different ecological zones and historical periods, as well as landraces and introduced varieties that potentially contributed genetically to these cultivars.

Recent reports indicated that although hundreds of genes with the potential to increase crop yields have been identified over the past two decades, few of these findings have translated into actual yield increases in production^4–5^. Because the yield effects of these genes are often assessed in greenhouses or small-scale field trials, few field trials have been designed to assess crop performance in different real-world environments or to define yield using only individual plant yield or yield component traits rather than grain-weight harvested per unit area^5^. Thus, we designed field trials that were consistent with actual production conditions across five ecological locations over two years. To the best of our knowledge, few previous studies measured yield data at a similar scale for wheat. This dataset allowed us to accurately assess the yield potential and to analyze the genetic basis for sustaining yield improvement.

### Homogenization and genetic diversity of modern cultivars

This germplasm panel, together with the corresponding phenotypic and genotypic data, provides an opportunity to trace the evolutionary history of wheat improvement in China and to dissect the genomic basis for the sustained increase in yield. The comprehensive and systematic analyses of this germplasm panel also provided novel and valuable insights for global wheat improvement efforts. Our results indicated that wheat improvement has largely reshaped the genomes of Chinese cultivars, which are facing challenges of narrow genetic diversity and severe homogeneity (Fig. 1). It is a key objective of future breeding to broaden the genetic diversity of modern wheat by capturing and introducing the diversity of landraces, introduced germplasms and wild relatives^64,67^. This requires strengthening the conservation and collection of germplasm resources and implementing protective measures for rare genetic materials to ensure their preservation during the breeding process. In addition, wheat from different zones or subzones exhibit distinct genetic structures, resulting from the differences in ecological environments and the implementation of targeted breeding by breeders. More precise deployment of specialized cultivars to local environments can increase yield^68^.

We conducted the first quantitative evaluation of the genetic contributions of Chinese landraces, introduced germplasms, and wild relatives to Chinese cultivars (Fig. 3 and Supplementary Fig. 4). Firstly, each landrace carries an average of 24.6% of rare haplotype segments compared to Chinese cultivar (10.7%) (Fig. 3a), indicating the untapped genetic diversity in landrace. The genomes of Chinese cultivars were largely shaped by genetic components from local landraces, which was consistent with the historical fact that early wheat breeding efforts in China primarily focused on the systematic reselection of local landraces^6–7^.Landraces exhibit optimal adaptation to local ecological conditions and production practices, along with corresponding production potential, serving as a foundational genetic resource for wheat improvement across local regions^6,64^. Directly exploring and utilizing the genetic diversity of landraces will be a more direct and effective approach^64^. Second, introduced germplasms provided 5.7% of HBs to Chinese breeding pool, which improved genetic diversity of Chinese cultivar population (Fig. 3c-f). Many introduced lines, such as ‘Quality’ and ‘Zhengyin 1’, were the direct or indirect parents of many cultivar^6^. Many HBs carried by Chinese cultivars may have originated directly from introduced lines, although they pre-existed in Chinese landraces. Meanwhile, introduced lines also contributed 11.6% in epistasis to Chinese cultivar population. Therefore, the introduced germplasms not only contributed some novel HBs to the cultivar population, but also epistasis. Third, we detected many alien chromosome segments in our germplasm panel (Fig. 3g-h and Supplementary Table 11), indicating that wild relatives have played an important role in Chinese wheat breeding^67,69–70^. For example, the germplasm panel contains 991 lines with 1BL.1RS translocations and 99 lines with 6VS.6AL translocations (Supplementary Fig. 4f). Alien segments such as 1RS and 6VS confer wheat with unique beneficial traits, particularly disease resistance^26,34,40,71^. In conclusion, future wheat breeding should focus on exploiting the genetic diversity of landraces, supplemented by the diversity of introduced germplasms and wild relatives.

### The potential of innovative germplasms in breeding

Since precisely identifying high-yielding and well-adapted varieties is of great significance for future sustainable development, we included 1,602 advanced breeding lines (i.e., innovative germplasms) in this study. Those lines were mostly developed after 2000 and were selected based on their potential in breeding programs in the next 10-20 years, such as high yield, high quality, disease resistance, or wide adaptability.

The results showed that the post-2000 cultivar group exhibited comparable agronomic performance, including yield, TKW, and spike grain number with innovative germplasms. Among the 150 most productive lines (top 5%) in the panel, 90 lines of various ecological regions belong to innovative germplasms (Supplementary Table 12). With larger-scale field trials, these high-yield lines may be directly utilized in actual agricultural production. Innovative germplasms carry many superior haplotype segments and epistatic modules (Fig. 6 and Supplementary Fig. 21), such as ‘Linhan 5175’ and ‘Xinong 4211’ (Supplementary Fig. 25-26). This highlights the importance of utilizing innovative germplasms in breeding programs to enhance the yield potential. In addition, although some innovative germplasms do not demonstrate similar performance in field trials, they exhibit unique advantages in certain traits. The superior alleles carried by these germplasms are high-value targets to be investigated to unlock their potential in breeding. It is worth mentioning that this panel was designed in 2016, and our classification of germplasm types is based on information up to 2023. As a result, more than 140 lines of them were identified as innovative germplasm at the time of collection but have now passed national approval and are classified in the 2010s group. In other words, the innovative germplasms have already demonstrated their potential. Thus, we believe that more innovative germplasms will be approved or widely utilized in future breeding programs.

### Epistatic modules and yield improvement

One perspective is that crop yield is controlled by a network of genetic interactions formed by thousands of loci, with only a limited number of core genes exerting significant direct effects on traits^59–60^. However, little solid scientific evidence supported this view. Our analysis provides strong support for this perspective, demonstrating that the epistatic effects between many haplotypes with insignificant effects on traits are significant (Supplementary Fig. 12). In addition, we found a significantly high positive correlation between yield and the number of favorable epistasis (Fig. 5), revealing that that yield is primarily driven by genetic interactions between a large number of small-effect haplotypes^53,55–58^. Moreover, these favorable epistases formed network modules (Supplementary Fig. 13a). The favorable epistatic network of current elite cultivars has expanded significantly compared with landraces and older cultivars (Supplementary Fig. 14). Further, high correlations between grain yield and the topology properties such as the number of modules were observed (Fig. 6 and Supplementary Fig. 15). These findings indicate that crop yield has been impacted by epistatic interaction networks formed by a large number of loci, and yield improvement is likely related to remodeling, expanding and optimizing epistasis networks.

To better understand the genetic mechanisms of yield improvement and to apply this theory in future wheat breeding, we propose a new conceptual framework (Supplementary Fig. 30). Combined GWAS with genetic interaction analysis can better reveal the genetic basis of quantitative traits. Our view highlights that future wheat breeding should be designed at the module level rather than focusing solely on a few alleles or haplotypes. Our findings also answer some outstanding questions in breeding. For example, some breeders have an intuitive understanding that crossbreeding between two elite wheat lines from different countries or ecological zones can sometimes make it difficult to breed high-yielding and widely adaptable cultivars. Conversely, a cross combination of two moderately or poorly performing lines may produce excellent cultivars. Our work may explain those phenomena - hybridization broke the originally favorable epistatic modules due to the differences between the genetic backgrounds of parents. When epistatic modules of the two lines are complementary, more favorable modules can be formed in the progeny. Therefore, our conclusions provide a theory for the selection of parental combinations in breeding. With the rapid development of artificial intelligence and molecular biotechnology, we can purposefully and efficiently breed varieties carrying targeted epistatic modules in the future.

## Materials and Methods

### Wheat material and genotyping

A total of 3030 wheat accessions and 7 controls were used in this study (Supplementary Table 1). Plant materials were available at Chinese Crop Germplasm Resource Information System (CCRIS) and more detailed information of accessions were queried from literatures^6–7^ and the website (http://202.127.42.47:6010/SDSite/Home/).

Genomic DNA was extracted from seedling leaves of each accession using a cetyl-trimethyl ammonium bromide method^72^ with minor modifications. The DNA concentration and quality were estimated using a Nano-Drop-2000 spectrophotometer (Thermo Fisher Scientific, Wilmington, DE) and electrophoresis on 1% agarose gels with a DNA marker. Genomic DNA was hybridized on Axiom Wheat660K Genotype Array by the China Golden Marker Biotechnology Company (Beijing, China). The genotyping was performed on the genotype array according to the Affymetrix Axiom 2.0 Assay Manual Workflow protocol. Variant quality was initially assessed according to Affymetrix best practices. The scores for the probes were classified into one of the following six categories according to the cluster patterns produced by the Affymetrix software: (i) Poly High Resolution (PHR), representing codominant and polymorphic SNPs, with at least two examples of the minor allele; (ii) No Minor Homozygote (NMH), representing polymorphic and dominant SNPs, where with two clusters were observed; (iii) Mono High Resolution (MHR), representing monomorphic SNPs; (iv) Call Rate Below Threshold (CRBT), where the SNP call rate was below the threshold but other cluster properties were above the threshold; (v) Off-Target Variant (OTV), which included four clusters, one of which represented a null allele; and (vi) other, where one or more cluster properties were below the threshold. A total of 233,422 markers, including 171,912 Poly High Resolution markers and 61,510 Off-Target Variant markers, were considered high quality, accounting for 35.4% of the Wheat660K array probes. The probe sequences were aligned to the reference (IWGSC RefSeq v1.0^31^) using BLASTN^73^ to obtain the position information of markers on chromosomes. A variation atlas that contained 201,740 SNPs were generated with a minor allele frequency at least 0.01 and missing rate no more than 0.4. The categories of identified SNPs were annotated using SnpEff^74^ (v5.0e).

### Field phenotyping

All accessions were planted in five different sites representing diverse wheat production regions of China. These locations can represent the environmental conditions of the major production regions: Shijiazhuang (38.05°N, 114.52°E) in Hebei Province, Tai’an (36.20°N, 117.08°E) in Shandong Province, Yangling (34.28°N, 108.07°E) in Shaanxi Province, Nanjing (32.07°N, 118.78°E) in Jiangsu Province, and Chengdu (30.67°N, 104.07°E) in Sichuan Province. Plantings were conducted across five ecological locations over two years (2017–2019), encompassing a total of 10 environments. Each accession was spaced and planted in a 2-m 5-row plot with 5 cm between plants and 25 cm between rows. Plots are spaced 1 row apart. Five local controls (Hebei: Shi4185; Shandong: Jimai22; Shaanxi: Xiaoyan22; Jiangsu: Yangmai20; Sichuan: Chuanmai42) and a unified control (Zhengmai7698) served as internal controls and were planted every 50 accessions. In each cropping season, fields were meticulously managed following common practices of local wheat production.

In total, 14 agronomic traits were evaluated, namely grain yield (GY), thousand kernel weight (TKW), grain number per spike (GNS), effective tiller number (ETN), spikelet number per spike (SNS), spike length (SL), sterile spikelet number per spike (SSN), kernel number per spikelet (KNS), plant height (PH), lodging resistance (LR), heading date (HD), flowering date (FD), maturation date (MD), and cold resistance (CR). Ten individual plants were randomly selected from the middle of each plot for phenotypic evaluation of GNS, ETN, PH, SL, SNS, SSN, and KNS for each accession. The yield of each plot was measured and converted to yield per hectare. CR was divided into five levels, with higher values indicating poorer cold tolerance. LR was divided into four levels, with higher values indicating poorer lodging tolerance.

### Population structure and genetics parameters

To visualize the genetic structure, PCA was performed using the smartPCA^75^ (v16000) based on all SNP sites. The genetic distance matrix was calculated based on all SNPs using PLINK^76^ (v1.90b6.24), and the neighbor-joining phylogenetic tree was built using PHYLIP^77^ (v 3.697) based on the matrix. Interactive Tree of Life^78^ (iTOL) was used to visualize the phylogenetic trees. The maximum likelihood estimation of individual ancestry for each possible group number (*K* = 2 to 20) was inferred using ADMIXTURE^14^ (v1.3.0) with five-fold cross-validations and 100 bootstrap replicates. Given that cross-validation showed decreasing errors for modeling scenarios with higher *K* values, and thus the delta of cross-validation error was calculated to consider an appropriate ancestry population number. The models with *K* > 12 showed little improvement in accuracy, and thus the output individual ancestry coefficient with *K*=12 was plotted using R package Pophelper^79^.

Genetic differentiation index (*F*_ST_) between groups was calculated using VCFtools^80^ (v0.1.16), with the parameters ‘--fst-window-size 1000000 --fst-window-step 500000’, and nucleotide diversity (π) of each group was calculated with the parameters ‘--window-pi 200000’. The IBS between individual pairs was calculated using --distance command in PLINK.

### Selection sweep

We used landraces as a reference population to detect improvement targets for cultivars of different groups. The XP-CLR statistic^15^ (https://github.com/hardingnj/xpclr) was used to identify selective sweeps between cultivars and landraces. XP-CLR was run with a sliding window size of 1Mb, a step size of 500 kb, a window containing at least six SNPs but less than 500 SNPs, and a correlation level of 0.95. In total, 7.10 Gb of genomic regions were successfully analyzed, covering 48.8% of the reference genome. Based on the XP-CLR scores for each window, the *splineAnalyze* function (smoothness=50,000, method=4) in GenWin (https://cran.r-project.org/web/packages/GenWin) was used to detect the boundary of genomic regions, resulting in a total of 3584 regions (N = 3584). We considered the score of the 36th region (top 1%; n ≈ 36, where n = N × 1%) as the threshold for selection sweeps. The significance thresholds for normalized XP-CLR scores were 2.17, 3.28, 4.98, and 3.45 for Zone 1, Zone 2, Zone 3, and Zone 4, respectively. Genomic regions with scores exceeding the respective thresholds were extracted for each population. Using IWGSC RefSeq v1.0 and its gene annotations, we extracted genes within the target genomic regions. GO enrichment analysis of these target genes was conducted with TBtools-II^81^ (v2.069).

### Inferring subgroup-specific haplotype blocks

The subgroup specific haplotype blocks were infer using the R package HaploBlocker^25^ (v1.6.06) based on the 3030 genotypes. The adaptive mode of block calculation function was used to create a haplotype library with the parameters ‘adaptive_mode = TRUE, window_size = c(5,6,7,8), node_min = 5, overlap_remove = FASLE, bp_map’. The parameter ‘node_min’ was used to control the number of haplotypes per node. The overlapping blocks were identified by setting parameter overlap_remove = FASLE. The parameter ‘bp_map’ was used to maintain the SNP position in the HB library. The presence (1) and absence (0) variation of HB for each chromosome were exported by *block_matrix_construction* function. Results of HB for 21 chromosomes were merged and generated a final presence-absence matrix. The HB position on chromosomes were calculated using custom Perl scripts according to the rank of allele sequences. Based on the previous division of wheat chromosome^31^ (R1, R2a, C, R2b and R3), the number and length of HB on each region were compared separately. Moreover, we ran HaploBlocker with the same parameters except ‘overlap_remove = TURE’ to calculate the amount of HB along the chromosome.

To evaluate the reliability of HB dataset, the HB -based PCA was performed based on the presence-absence matrix using *prcomp* function in the R Stats package^82^ with the parameters ‘center = T and scale. = T’. HB-based PCA was highly consistent with SNP-based PCA, indicating the high accuracy of inferred HBs. The HB heatmaps were plotted using *plot_block* function of HaploBlocker. The chromosome 1RS from rye was characterized by HBs (Fig. 2a), further suggesting the reliability of this HB.

### Phenotypic BLUP

Pearson correlation coefficients of phenotypic data from different environments were calculated using *cor* function in the R Stats package. To remove environmental effects and obtain stable genetic phenotypes of individuals, best linear unbiased prediction (BLUP) is used to fit phenotypic data of all environments. BLUP was calculated using lme4 packages^83^ (v3.2.2) in R. The formula was as follows:

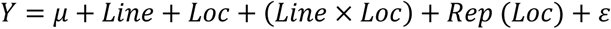

Where *Y*, *μ*, Line, and Loc represent phenotype, intercept, variety effects, and environmental effects, respectively. Rep means different replications, and *ε* represents random effects. *Line* × *Loc* is used to display the interaction between variety and environment, and *Rep* (*Loc*) shows the nested effect of replication within the environment. Using the BLUP, Pearson correlation coefficients was calculated between fourteen agronomic traits.

### Heritability analysis

The narrow sense heritability of 14 traits was calculated for the phenotypic data of each environment and the BLUP using GCTA^84^ (v1.93.3beta2). Based on all SNPs, genetic relationship matrix^85^ (GRM) was estimated using GCTA with the parameters: ‘--make-grm-inbred --make-grm-alg = 0’. The first five PCs based on all SNPs were used as a covariate to control the population structure. The heritability of 14 traits were calculated based on the GRM, covariate and phenotypes. We compared the heritability of different genetic information, including (1) all SNPs, (2) all HBs, (3) SNPs of GWAS hit (*P* < 0.01), (4) HBs of GWAS hit (*P* < 0.05), and (5) epistasis (*P* < 1×10^−16^). For HBs and epistasis, the presence-absence matrix (Supplementary Data 1 and Supplementary Data 2) was used to calculate heritability.

### GWAS and QTL analysis

The SNP-based GWAS and HB-based GWAS were performed to identify SNPs and HBs that have significant associations with 14 agronomic traits. A total of 183,440 biallelic SNP sites with minor allele frequency at least 0.01, missing genotype ratio per site less than 0.4 and heterozygous genotype ratio per site less than 0.03 were used for SNP-based GWAS. A total of 18,779 HBs with frequency at least 20 (≥ 0.007) were used for HB-based GWAS. Associations based on the presence and absence of HB were performed for HB-based GWAS. To reduce false positives, the GRM was provided to consider the kinship between samples, and the first five PCs were used to control the population structure. The mixed linear model (MLM) implemented in GCTA program was used to perform genotype-phenotype associations. The formula of MLM was as follows:

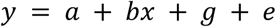

Where *y* is the phenotype, *a* is the mean term, *b* is the additive effect (fixed effect) of the candidate SNP to be tested for association, *x* is the SNP genotype indicator variable coded as 0, 1 or 2, *g* is the polygenic effect (random effect) i.e. the accumulated effect of all SNPs (as captured by the GRM calculated using all SNPs) and *e* is the residual^84^.

We defined QTLs and its intervals using a linkage disequilibrium (LD) clumping approach^76^. LD clumping was performed using the --clump command in PLINK. The analysis was conducted with the following parameters: a primary variant significance threshold of *P* ≤ 1×10^−5^ (--clump-p1) to identify candidate lead SNPs, a secondary variant significance threshold of *P* ≤1×10^−3^ (--clump-p2) to include additional variants for LD analysis, an LD threshold of *r*^2^ ≤ 0.2 (--clump-r2) to define LD relationships, a clumping window size of 5000 kb (--clump-kb) to restrict analysis to a ±5 Mb region, and the --clump- allow-overlap option to permit overlapping regions between different clumps. Using the *clump* function, lead SNPs representing QTLs were identified. For each QTL, the QTL interval was determined based on the positions of the lead SNP and the SNPs linked to it that met both the --clump-p2 significance threshold and the --clump-r2 LD threshold. For genome-wide multiple-test correction, the nominal significance level 0.05 was divided by the number of independent SNPs (N) pruned using PLINK with the parameter ‘--indep-pairwise 100 1 0.2’ (0.05/N ≈ 10^−5^). We then consider a lead SNP with *P* ≤ 1×10^−5^ as a robust QTL signal.

The proportion of variance explained (PVE) by the QTLs (lead SNPs) was calculated using the following formula^86^:

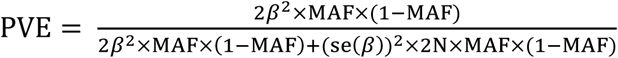

where MAF is the minor allele frequency of the lead SNP, N is the number of individuals that participated in the analysis of the SNP in GWAS, *β* is the effect value in GWAS, and se is the standard error of the effect value in GWAS.

### Structural equation modeling

We used the structural equation model (SEM) to quantify the multivariate causal network of the GY and GY components. SEM was performed using lavaan^48^ (v0.6-18) package in R. TKW, grain number per spike (GNS), lodging resistance (LR) and flowering date (FD) were considered direct factors, while effective tiller number (ETN), spikelet number per spike (SNS), kernel number per spikelet (KNS), sterile spikelet number per spike (SSN), spike length (SL), plant height (PH) and heading date (HD) were considered indirect factors. The model includes the following pathways: GY is regressed on TKW, GNS, LR, and FD (GY∼TKW+GNS+LR+FD); Covariance relationships are specified between TKW and GNS, LR, and FD (TKW∼∼GNS; TKW∼∼LR; TKW∼∼FD), as well as between GNS and LR (GNS∼∼LR); TKW is regressed on SNS, ETN, KNS, SL, SSN, HD, and PH (TKW∼SNS+ETN+KNS+SL+SSN+HD+PH); Similarly, GNS is regressed on SNS, ETN, KNS, SL, SSN, HD, and PH (GNS∼SNS+ETN+KNS+SL+SSN+HD+PH); FD is regressed on HD (FD∼HD); LR is regressed on PH (LR∼PH). Samples containing missing phenotype values (NA) were removed from the dataset using listwise deletion before the SEM analysis. To make coefficients of different pathways comparable, all selected variables were standardized to have a mean of 0 and a standard deviation of 1. We used standardized path coefficients (std.all) from the structural equation model estimated using the lavaan package. These coefficients standardize all variables (both latent and observed) to have a mean of 0 and a standard deviation of 1, allowing for direct comparison of the relative magnitudes of effects across all paths in the model. Goodness-of-fit index (GFI), adjusted goodness-of-fit statistic (AGFI), standardized root mean square residual (SRMR) and comparative fit index (CFI) were used to assess the global fit of SEM model.

### Identifying large alien chromosome translocations

Since Axiom Wheat 660K Genotyping Array are designed based on wheat DNA sequences^13^, the probes can only be hybridized with homologous sequences that are consistent with wheat and relatives. Thus, genotyping of alien chromosome segments using the array results in a continuous and high missing call but not all. Based on this characteristic, we detected the large chromosome exchanges by calculating the SNP missing ratio of each genomic region.

First, for each genotype, we calculated the missing ratio of each slip window by taking 100 SNPs as a sliding window and 20 SNPs as a step. Then, we merged adjacent windows with the missing ratio greater than 0.2. We also merged windows that are less than 2 Mb apart on R1 and R3, 3Mb apart on R2a and R2b and 5 Mb apart on C region, and then the position of alien segments relative to the reference genome was calculated for each genotype. Finally, based on the consistency of breakpoints and the corresponding genotype heatmap, it was determined whether an alien segment carried by different genotype is the same source. Therefore, the frequency of alien segments can be calculated by this method. We speculated the possible donors of several alien segments based on the pedigrees and literatures.

### Epistasis calculation and network topology analysis

HB instead of SNP was used to detect epistasis, because HBs inherently model local epistasis, and thereby solves some of the general problems of markers being correlated but not causal individually^25,87–88^. Additionally, HBs represent a series of physically linked and co-inherited alleles, making them an effective genetic unit for breeding selection^21,24–25^. We use MatrixEpistasis^61^ (v0.1.0) to detect two-dimensional epistasis (HB-HB interaction). MatrixEpistasis can ultrafast and exhaustively scans all pairwise epistatic interactions for quantitative traits with covariate adjustment. The statistical model with covariate adjustment was:

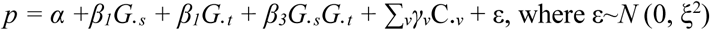

where *α* is the overall mean of the quantitative phenotype; The regression coefficients *β_1_*/*β_2_*, *β_3_* and *γ_v_* are the main genetic additive effect, interaction effect, and covariates, respectively; ε is a normal variable with zero mean and ξ^2^ variance^61^. The regression coefficient *β_3_* gives the size of the effect that the interaction term is having on the phenotype with both main genetic additive effects and covariates adjusted. Therefore, the hypothesis is defined as *H*_0_: *β_3_* = 0 and *H*_1_: *β_3_* ≠ 0, where testing for interaction effects involves assessing whether *β_3_* equals zero^61^. To improve computing efficiency, MatrixEpistasis calculates only *β_3_* and exhaustively scans all pairwise epistasis interactions, identifying only the significant ones^61^.

To reduce false positives, the first five PCs were used as a covariate adjustment to control population structure. We calculated epistasis for GY, TKW, and GNS, and considered HB-HB combinations with *P* less than 1×10^−16^ as candidate epistasis interactions that are associated with traits. This threshold for the *P* is rigorous and robust^58^ and lower than the level of genome-wide significance (0.001/N = 2.8×10^−12^, where N is the number of HB-HB combinations). Based on the presence-absence matrix of HBs, we extracted the epistasis carried by each sample and constructed the presence-absence matrix of epistasis (Supplementary Data 2). We defined favorable, inferior, and moderate epistasis for each HB-HB interaction. For example, when the yield of accessions carrying a certain HB-HB interaction (AABB) is significantly higher than that of accessions without the combination (AA and BB), this HB-HB interaction (AABB) is considered favorable epistasis, and conversely inferior or moderate. (Supplementary Fig. 12b).

We analyze epistatic networks at the group and individual levels. We use the favorable epistasis carried by all samples as a global layout. For group-level network analysis, we extract all epistasis (i.e. edges) contained by each group and then calculate the frequency of each edge. For the favorable epistasis network of each individual, the igraph^89^ (v1.2.11) in R was used to calculate the topological properties of the epistasis network. The properties include node count, edge count, mean neighbor, graph diameter, mean component size, component count, node count of maximum component, edge count of maximum component, a mean neighbor of maximum component, maximum module size, mean module size, max module size, and module count. For individual-level network module analysis, modules were counted only if they contained at least 5 nodes. The Gephi^90^ (v0.10) was used to visualize the networks.

## RESOURCE AVAILABILITY

### Lead contact

Further requests and information concerning this study should be addressed to the lead contact, Lihui Li.

### Materials availability

All the plant materials are available on request to the lead contact, Lihui Li.

### Data and code availability

The variant call format (VCF) file and haplotype block atlas of 3030 accessions based on Axiom Wheat 660K Genotype Array have been deposited at Genome Variation Map - National Genomics Data Center (GVM000959, [GVM000972](https://ngdc.cncb.ac.cn/gvm/submit/sub001828/accession) - PRJCA035838). Information of all accessions were provided in Supplementary Table 1. The phenotypic data was released in Supplementary Table 12.

## ACKNOWLEDGEMENT

This work was supported by the National Key Research and Development Program of China (2021YFD1200605 and 2016YFD0100102). The authors thank all the students, researchers and employed workers for their contribution to field trails and collecting agronomic trait data.

## AUTHOR CONTRIBUTION

M.Y., H.H., and F.H.—bioinformatical and statistical analyses of data; Z.Z., Y.H., H.W., C.Z., X.Z.L., and Q.L.—helped with data analyses; M.Y., H.H., F.H., H.-Q.L., and L.L.—data interpretation; L.L., H.H, W.L., L.K., X.P.L., W.J., J.W., G.D., H.D., J.Z., S.Z., X.G., and X.Y.—assembling the diversity panel and conducted field trials, H.H.—array-based genotyping and data quality control; M.Y., F.H., H.H., and L.L.— manuscript writing; L.L.—conceived idea and supervised the project.

## Supplementary Materials

### Supplementary Notes

#### Supplementary Note 1

##### Indirect effects of yield-related traits on yield

In addition to analyzing the direct effects of thousand kernel weight (TKW), grain number per spike (GNS), lodging resistance, and flowering date on yield, we also examined the indirect effects of other yield-related traits, including spikelet number per spike, kernel number per spikelet, sterile kernel number, spike length, effective tiller number, plant height, and heading date (Fig. 4b and Supplementary Table 14). These traits indirectly influence yield through their effects on TKW or GNS. Structural equation modeling (SEM) analysis across all Chinese landraces and cultivars revealed that TKW was primarily influenced by plant height, followed by flowering date, kernel number per spikelet, and effective tiller number. GNS was mainly affected by kernel number per spikelet, followed by spikelet number per spike, sterile kernel number, and spike length. Additionally, flowering date was strongly influenced by heading date, while lodging was primarily determined by plant height. Specifically, reducing plant height and advancing heading date had positive effects on TKW. Increasing spikelet number per spike, kernel number per spikelet, and spike length while reducing sterile kernel number contributed to an increase in GNS. However, notable trade-offs were observed between certain traits. For instance, a reduction in spikelet number per spike and kernel number per spikelet could enhance TKW but would consequently reduce GNS. Overcoming these trade-offs remains a critical goal in future wheat breeding efforts.

The effects of yield-related traits on TKW and GNS have undergone significant shifts across different historical stages. From the 1950s to the 1990s, TKW was primarily influenced by plant height, followed by spikelet number per spike, and GNS was mainly affected by kernel number per spikelet, followed by spikelet number per spike and sterile kernel number. However, after the 2000s, TKW became predominantly influenced by kernel number per spikelet, followed by effective tiller number, and GNS remained largely determined by kernel number per spikelet, followed by spikelet number per spike and spike length. In innovative germplasms (i.g. CIGs), TKW was primarily influenced by kernel number per spikelet, followed by spikelet number per spike and effective tiller number, while GNS was mainly affected by kernel number per spikelet, followed by spikelet number per spike and sterile kernel number.

Spatially, different wheat ecological zones exhibited distinct patterns for routes of increasing TKW. In Zone 1, TKW was primarily influenced by spikelet number per spike, followed by effective tiller number and plant height. In Zone 2, TKW was mainly affected by kernel number per spikelet, followed by effective tiller number and spikelet number per spike. In Zone 3, TKW was predominantly determined by plant height and sterile kernel number, followed by spikelet number per spike and effective tiller number. In Zone 4, TKW was mainly influenced by heading date, followed by kernel number per spikelet, effective tiller number, spikelet number per spike, and spike length. However, the primary strategy to increase GNS across all ecological zones was to increase spikelet number per spike and kernel number per spikelet. The effects of spike length and sterile kernel number on GNS varied among ecological zones. For example, in Zone 1, reducing sterile kernel number increased GNS, while spike length had no significant effect. In Zone 2 and Zone 3, reducing sterile kernel number had a greater positive impact on GNS than increasing spike length. In contrast, in Zone 4, spike length played a more substantial role in GNS increase than sterile kernel number.

#### Supplementary Note 2

##### Definition of epistasis

Epistasis is defined differently in physiological and statistical genetics. Distinguishing between these definitions can deepen our understanding of epistatic interactions. In physiological genetics, epistasis occurs when phenotypic differences among individuals with different genotypes at one locus depend on their genotypes at another locus^1,2^. In statistical genetics, epistasis is the deviation of the genotypic value of two or more loci from the sum of the genotypic values contributed by each locus individually, defined as the departure from additive effects in the linear model^1–3^. Consequently, the effect of a two-locus genotype cannot be predicted from the average effects of each locus separately^4^.

There are two key inherent distinctions in these definitions. First, statistical epistasis is a population-level phenomenon depending on allele frequencies in a specific population, whereas physiological epistasis is a genotype-level phenomenon, independent of allele frequencies^2^. Thus, when the minor allele frequency is low, the magnitude of epistatic effects may exist but cannot be detected in the epistatic variance. Second, statistical epistasis deviations associated only with the variance component of interaction, whereas physiological epistasis not only can contribute the epistatic variance component but also additive and dominance variance components^2^. Thus, the magnitude of physiological epistatic effects cannot be inferred from the variance components of statistical epistasis^4^. Current theories on statistical epistasis primarily focus on interactions between two loci (SNP-SNP interactions). For two loci (A and B) affecting the same quantitative trait, the effect of any two-locus genotype can be expressed as the sum of the additive (*a*_A_ and *a*_B_), dominance (*d*_A_ and *d*_B_) of the constituent genotypes, and the epistatic effects (*i*_AB_)^4^. These genetic effects can be estimated by measuring the average phenotypes of individuals with each of the nine causal two-locus genotypes^4,5^, as detailed by Mackay *et al*.

#### Supplementary Note 3

##### Statistics of epistatic test results

Using yield as an example, we described the epistasis calculation results. A total of 13,119 HB-HB interactions (epistasis) significantly associated with yield were identified through genome-wide epistasis mapping. Among these epistatic interactions, 83.4% (10,937/13,119) pre-existed in Chinese landraces, whereas the remaining (2,182/13,119) were novel interactions that emerged in modern breeding (Supplementary Fig. 9a). Of the epistatic interactions absent in landraces, 90.7% (1,518/1,682) were detectable in introduced germplasms, suggesting that introduced germplasms have contributed new interactions. In other words, introduced varieties contributed not only 5.7% HBs but also 11.6% epistasis (1,518/13,119) to Chinese cultivated population. Although these new interactions are absent in Chinses landraces, most of the HBs involved in these interactions pre-existed in landraces, meaning that the two HBs were not simultaneously present in the same landrace. Thus, these epistases can be generated through reassembly of HBs.

27.7% (3,636/13,119), 21.2% (2,782/13,119), and 51.2% (6,721/13,119) of the HB-HB interactions were classified as favorable, inferior, and moderate epistasis, respectively (Supplementary Table 19). For epistasis pre-existing in Chinese landraces, 22.2% (2,431/10,937), 21.1% (2,304/10,937), and 56.7% (6,202/10,937) of the HB-HB interactions were classified as favorable, inferior, and moderate epistasis, respectively, while for novel epistasis, 54.3% (1,185/2,182), 21.9% (478/2,182), and 23.8% (519/2,182) of the HB-HB interactions were classified as favorable, inferior, and moderate epistasis, respectively.

#### Supplementary Note 4

##### Epistatic module analyses

We also examined the transmission, refinement and generation of favorable epistatic modules in other pedigrees.

‘Xinong 619’ was bred through the hybridization between ‘Zhoumai 16’ and ‘Xinong 4211’ (Supplementary Fig. 26). Module analysis of this pedigree showed that ‘Xinong 619’ primarily inherited and integrated modules M2 and M9 from its two parents, inherited M13 and M14 from ‘Zhoumai 16’, and inherited M3 and M15 from ‘Xinong 4211’.

‘Zhoumai 26’ was bred through the hybridization between ‘Zhoumai 22’ and ‘Zhoumai 24’ (Supplementary Fig. 27). ‘Zhoumai 26’ primarily inherited modules M3, M7, M8, and M14 from its parents while integrating and refining modules M2, M9, M12, and M15. Notably, the completeness of module M12 in the two parents was only 18%, whereas in the progeny ‘Zhoumai 26’, it increased significantly to 61%. Additionally, components of modules M9 and M15 were scarcely present in the parents but showed a marked increase in completeness in ‘Zhoumai 26’.

‘Zhoumai 36’ was bred through the hybridization between ‘Zhoumai 24’ and ‘Aikang 58’ (Supplementary Fig. 28). Module analysis of this pedigree revealed that ‘Zhoumai 32’ inherited modules M3, M7, and M11 from ‘Zhoumai 24’ module M13 from ‘Aikang 58’ and modules M2, M8, M11, M12, and M14 from both parents. Notably, the high-yielding variety, ‘Aikang58’, have a module M2 completeness of 53%.

‘Zhoumai 36’ was bred by crossing ‘Aikang 58’ and ‘Zhoumai 19’, and then hybridizing with ‘Zhoumai 22’ (Supplementary Fig. 29). Module analysis of this pedigree demonstrated that ‘Zhoumai 36’ inherited module M13 from both ‘Aikang 58’ and ‘Zhoumai 19’, module M15 from ‘Aikang 58’, and modules M2, M12, and M14 from multiple parental sources.

Module analysis of these pedigrees suggest that breeding selection has enabled many genetic modules to remain dynamically stable to a significant extent. Epistatic modules can be reshaped and optimized through genetic recombination, while new favorable modules can be generated and selected during the breeding process.

### Supplementary Figures

**Supplementary Fig. 1.**
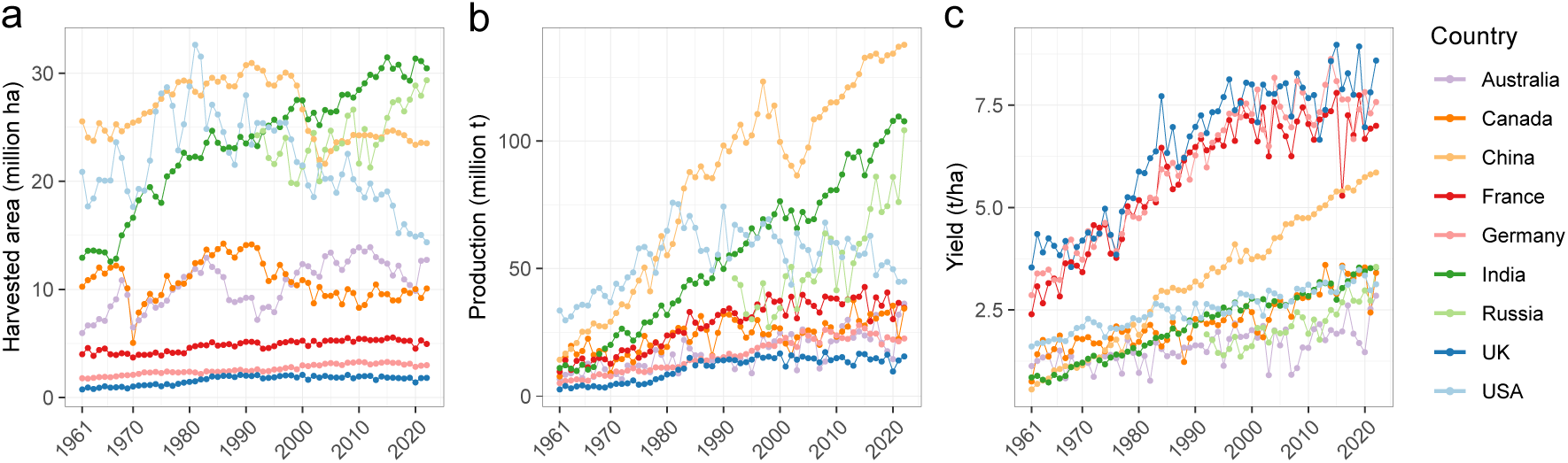
Harvested area (a), production (b) and yield (c) of wheat in nine countries from 1961 to 2022. The statistic data is from Food and Agriculture Organization of the United Nations (https://www.fao.org/home/en).

**Supplementary Fig. 2.**
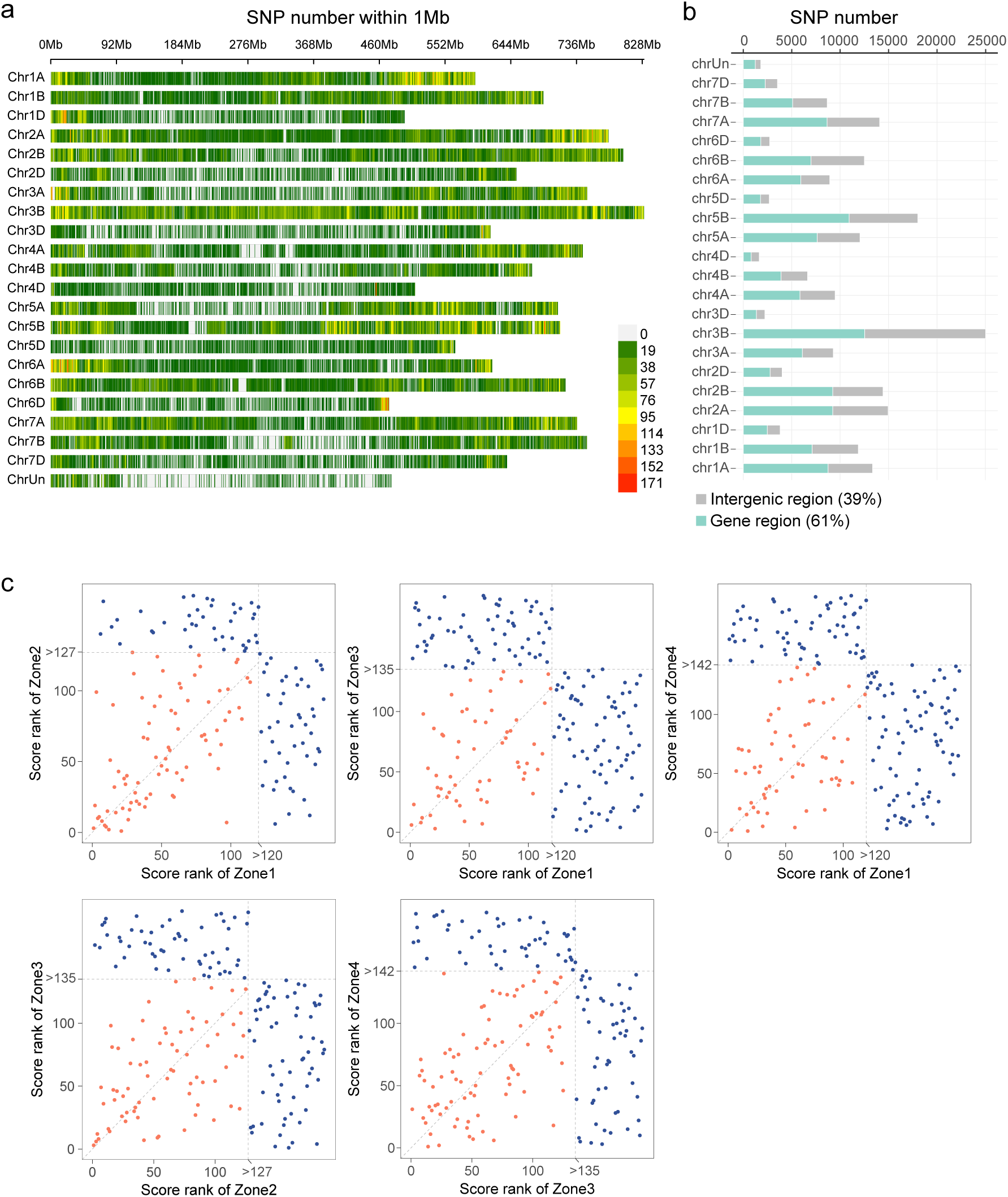
SNP density and XP-CLR analysis. **a**, Density distribution of SNPs. **b**, Number of SNPs of each chromosome. The division of gene region and intergenic region is based on the annotation of SnpEff. **c**, Ranking comparison of XP-CLR scores between different zones. Higher rankings represent stronger selection intensity. The gray dashed line represents the top 1% threshold. Each dot represents a genomic region of 1 Mb. Red dots indicate loci under selection in both ecological zones, whereas dark blue dots indicate loci under selection in only one ecological zone.

**Supplementary Fig. 3.**
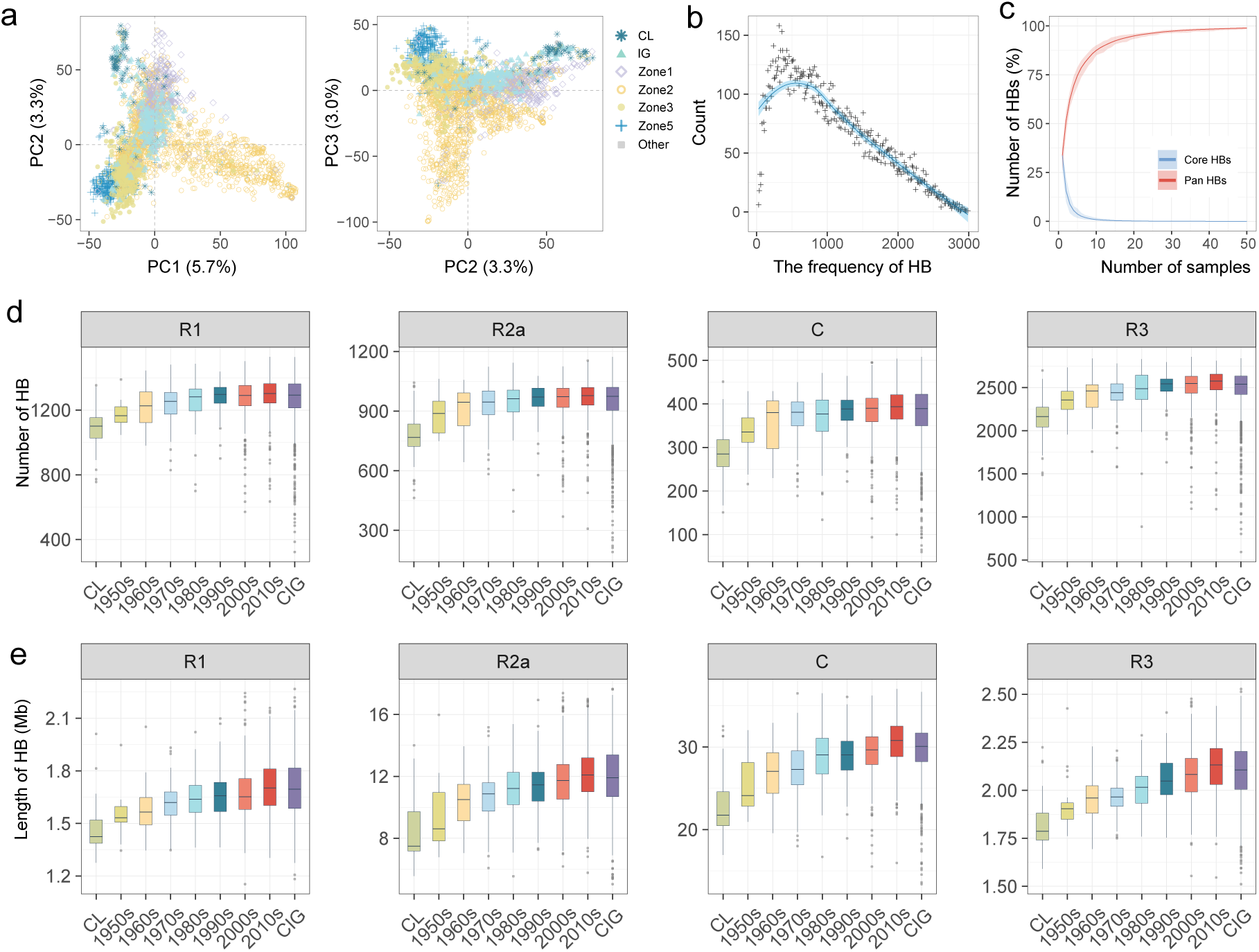
Characteristics of haplotype blocks (HBs). **a,** HB-based principal component analysis (PCA). **b,** Frequency distribution of HBs in the panel. **c,** Changes in the number of pan-HBs and core-HBs as the sample size increases. **d-e,** Number (**d**) and length (Mb) (**e**) of HBs carried by wheat lines from different groups on region R1, R2a, C, and R3. R1 and R3 represent the terminal regions of the short arm and long arm, R2a represents the middle region of the short arm, and C represents the centromere and its adjacent regions. CL: Chinese landraces; CIG: Chinese innovative germplasms.

**Supplementary Fig. 4.**
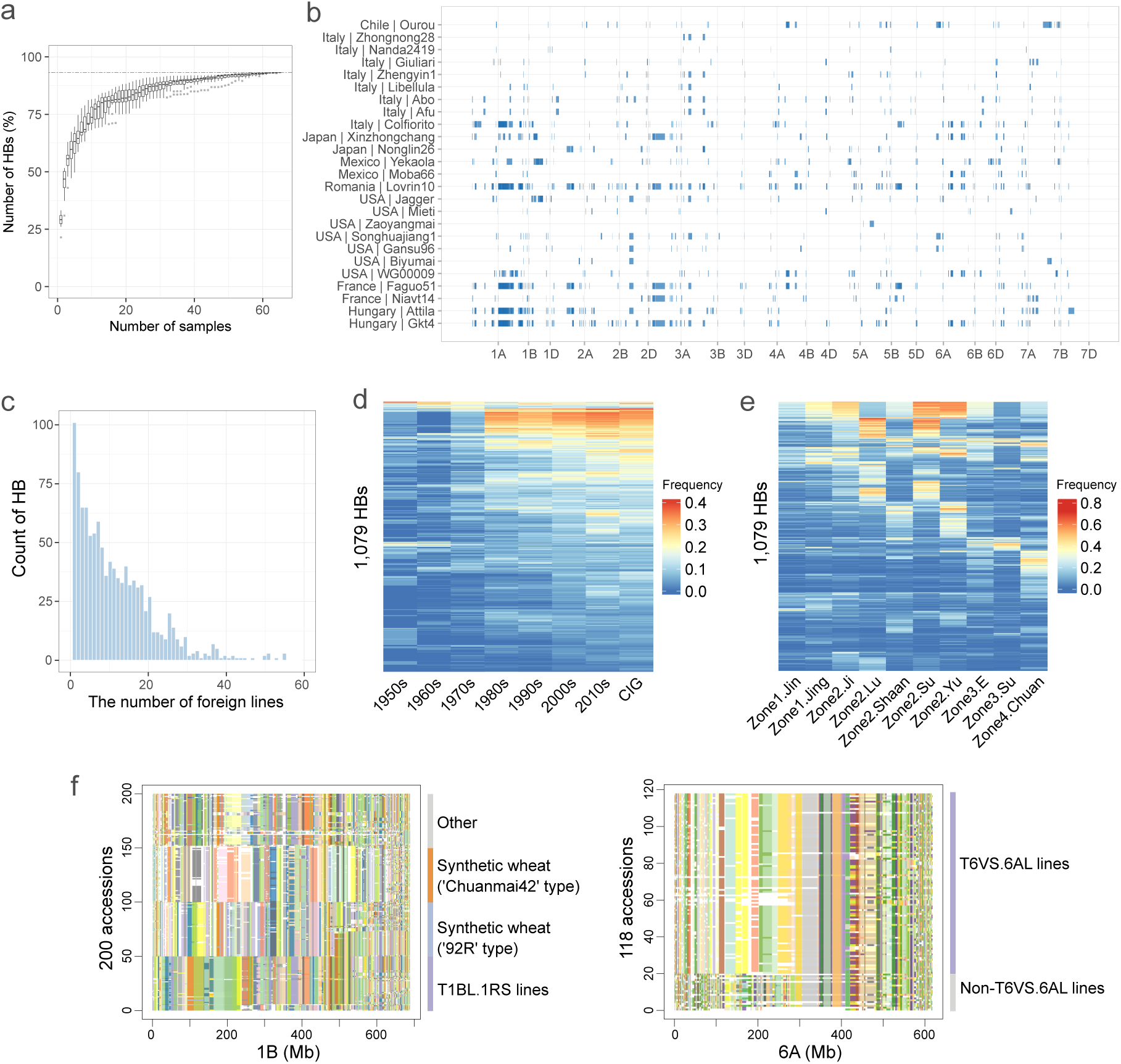
Analysis of HBs originating from Chinese landraces, introduced lines, and wheat relatives. **a,** Changes in the accumulated number of HBs as the sample size of the landraces increases from 1 to 65. For each sample size, perform 10 random samplings and plot the results as a boxplot. When the sample size reaches 65, the cumulative number of non-redundant HBs is 17,498 (blue dashed line). **b,** Frequency distribution of introduced--derived HBs among the population of introduced wheat lines. **c**, 25 representative introduced lines contributed new HBs to the Chinese breeding pool, along with their distribution across chromosomes. **d,** Frequency of introduced-derived HBs populations from different eras. **e,** Frequency of introduced-derived HBs populations from different subzones. **f**, Left: HB heatmap of chromosome 1B across four types of wheat populations, including 50 T1BL.1RS lines, 50 ‘92R’ type synthetic wheat, 50 ‘Chuanmai42’ type synthetic wheat, and 50 other wheat lines. One of the donors of these synthetic wheats is durum wheat. Previous studies have suggested that *Y24/Y26/YrCh42* from durum wheat have multiple sources^6^, and our HB analysis supports this conclusion; Right: HB heatmap of chromosome 6A for 98 T6VS.6AL lines and 20 non-T6VS.6AL lines.

**Supplementary Fig. 5.**
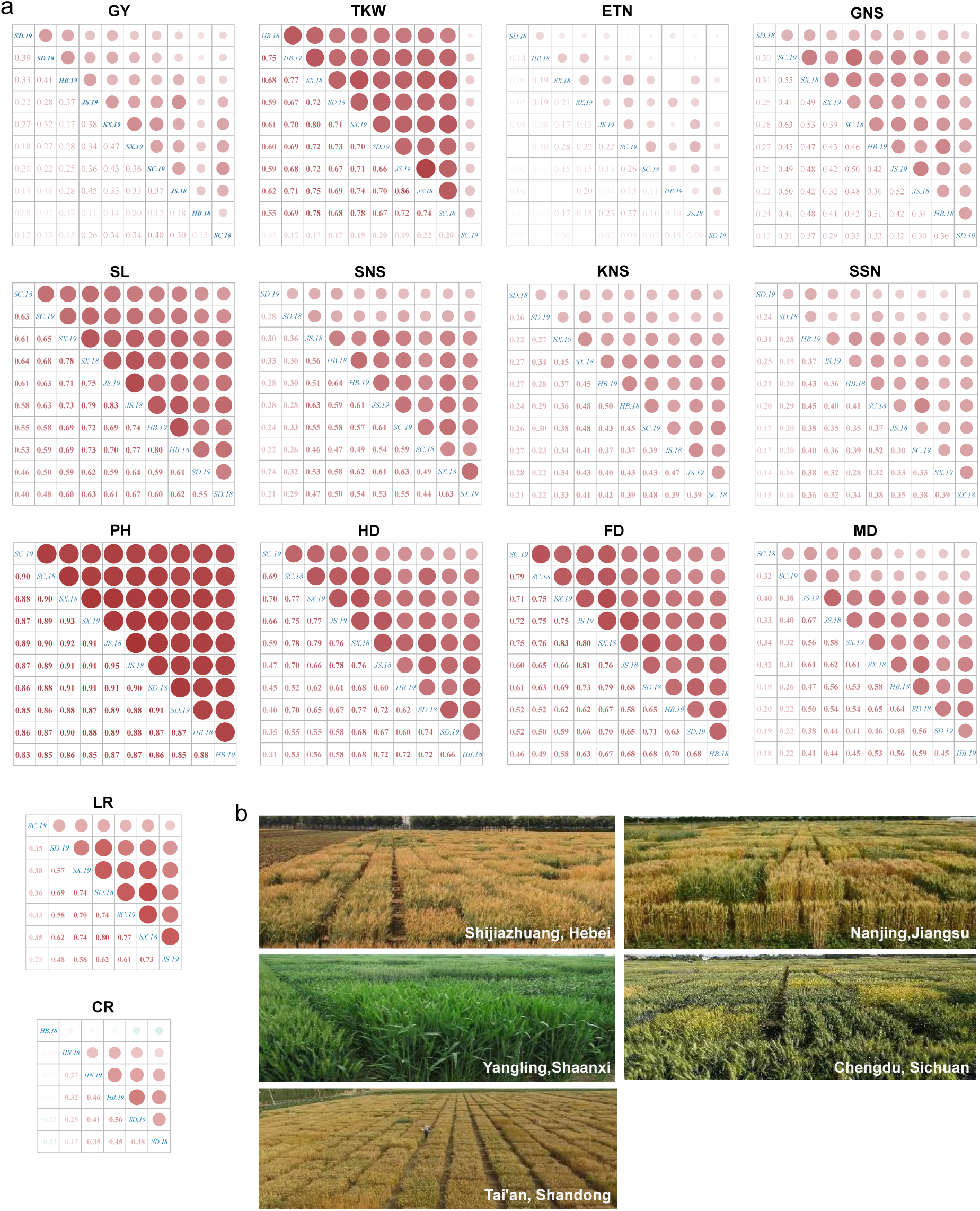
Phenotypic data collection. **a,** Pearson correlation coefficient (PCC) between phenotypic datasets from different environments. GY: grain yield (t ha^−1^); TKW: thousand grain weight; ETN: effective tiller number; GNS: grain number per spike, SL: spike length; SNS: spikelet number per spike; KNS: kernel number per spikelet; SSN: sterile spikelet number per spike; PH: plant height; FD: flowering date; HD: heading date; MD: maturation date; LR: lodging resistance; CR: cold resistance. SD: Shandong; HB: Hebei; SX: Shaanxi; JS: Jiangsu; SC: Sichuan. **b**, Field trials at six ecological sites, including Tai’an (36.20°N, 117.08°E) in Shandong Province, Shijiazhuang (38.05°N, 114.52°E) in Hebei Province, Yangling (34.28°N, 108.07°E) in Shaanxi Province, Nanjing (32.07°N, 118.78°E) in Jiangsu Province, and Chengdu (30.67°N, 104.07°E) in Sichuan Province. Field trials were conducted from 2017 to 2019. Phenotypic data of LR and CR were collected from 7 and 6 environments, respectively.

**Supplementary Fig. 6.**
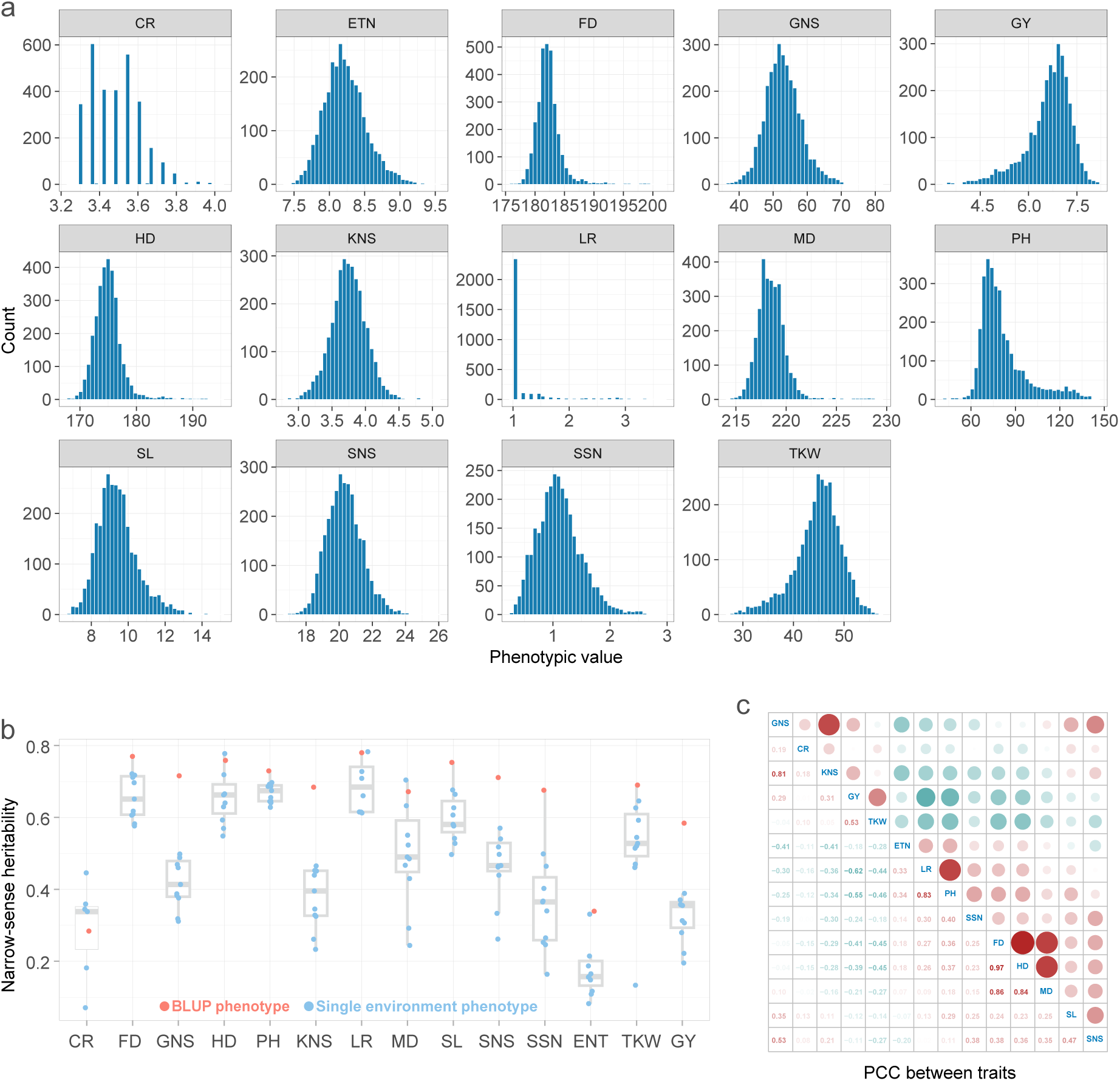
Analyses of phenotypic data. **a,** Distribution of BLUP values for 14 agronomic traits. **b,** Narrow-sense heritability analysis. Blue points represent heritability calculated based on phenotypic data from a single environment, while red points represent heritability calculated based on BLUP values. **c,** Pearson correlation coefficients between different traits.

**Supplementary Fig. 7.**
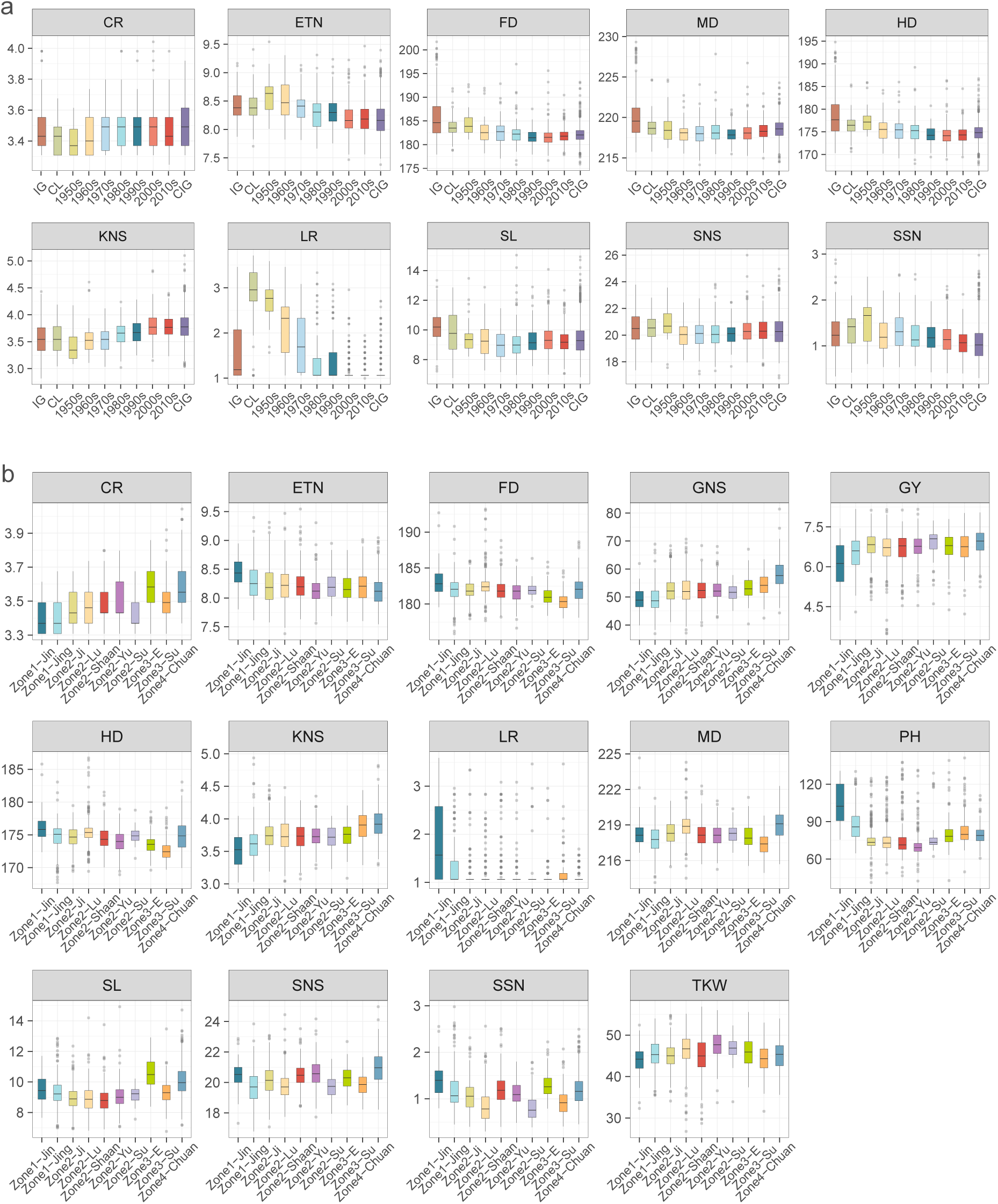
Comparison of phenotypes across different groups. **a,** Comparison of phenotypes across wheat groups from different eras. Each point represents a sample, and boxplots were plotted based on era groupings. **b,** Comparison of phenotypes across wheat groups from different subzones. Each point represents a sample, and boxplots were plotted based on subzone groupings. IG: introduced germplasms.

**Supplementary Fig. 8.**
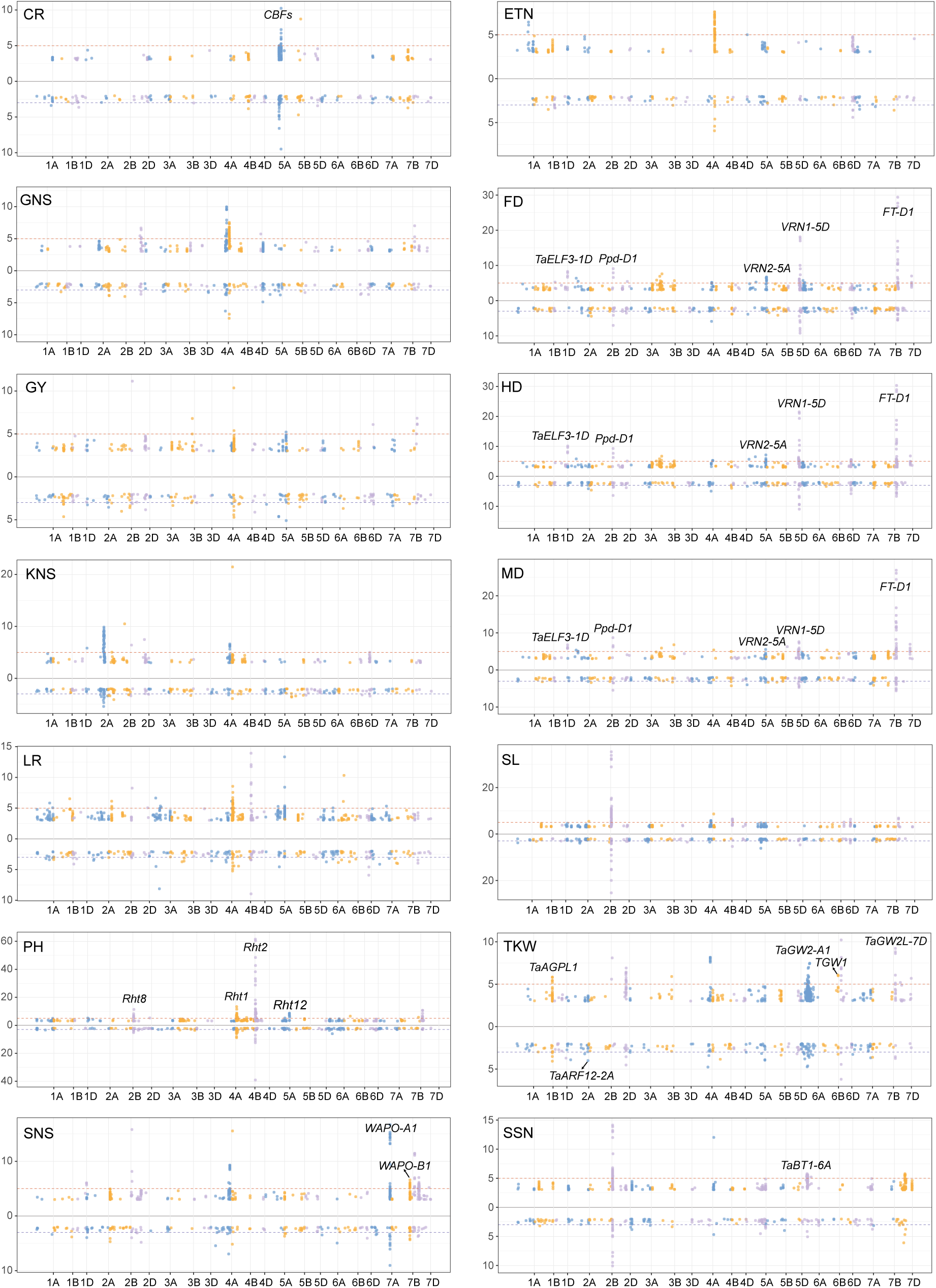
Genome-wide association study (GWAS) of 14 agronomic traits. The upper panel represents SNP-based GWAS, and the lower panel shows HB-based GWAS. For SNP-based GWAS, the multiple-test corrected significance threshold (1×10^−5^) was denoted with red horizontal dashed lines, while for HB-based GWAS, the threshold is set at 1×10^−3^. For SNP-based GWAS, only loci with *P* < 1×10^−3^ are displayed, while for HB-based GWAS, only HBs with *P* < 1×10^−2^ were shown. Some genes with known functions were labeled in the Manhattan plot.

**Supplementary Fig. 9.**
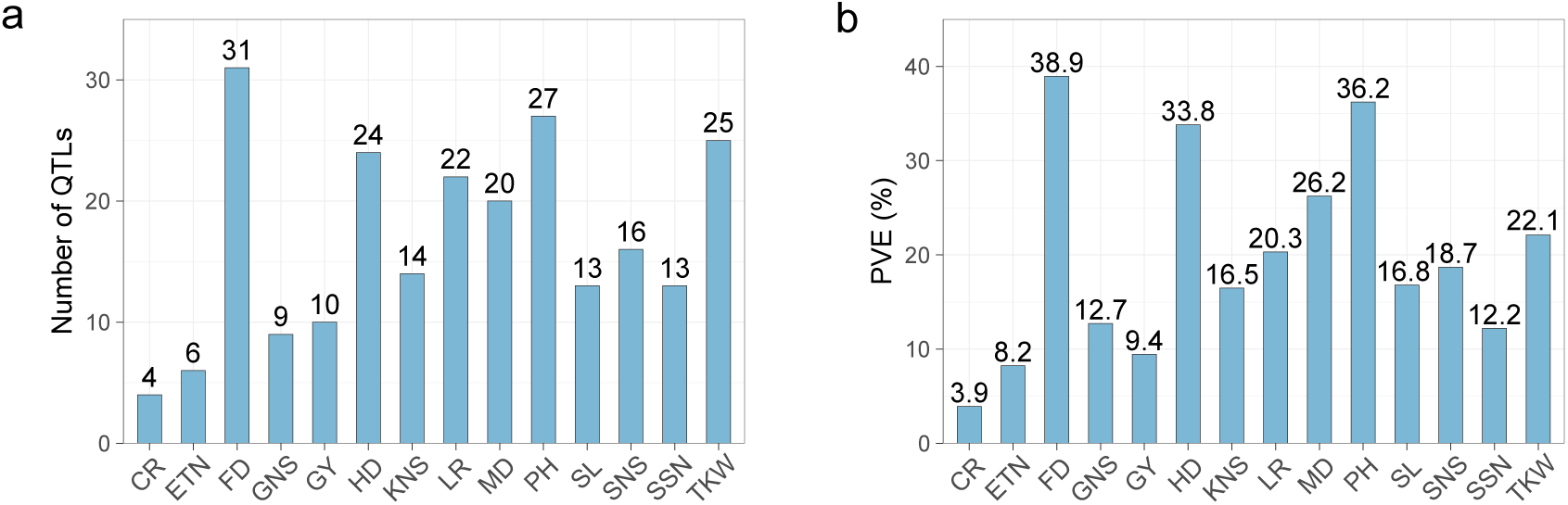
Summary of QTLs. **a**, Number of QTLs identified for each trait. **b**, Total phenotypic variance explained (PVE) by the QTLs identified for each trait.

**Supplementary Fig. 10.**
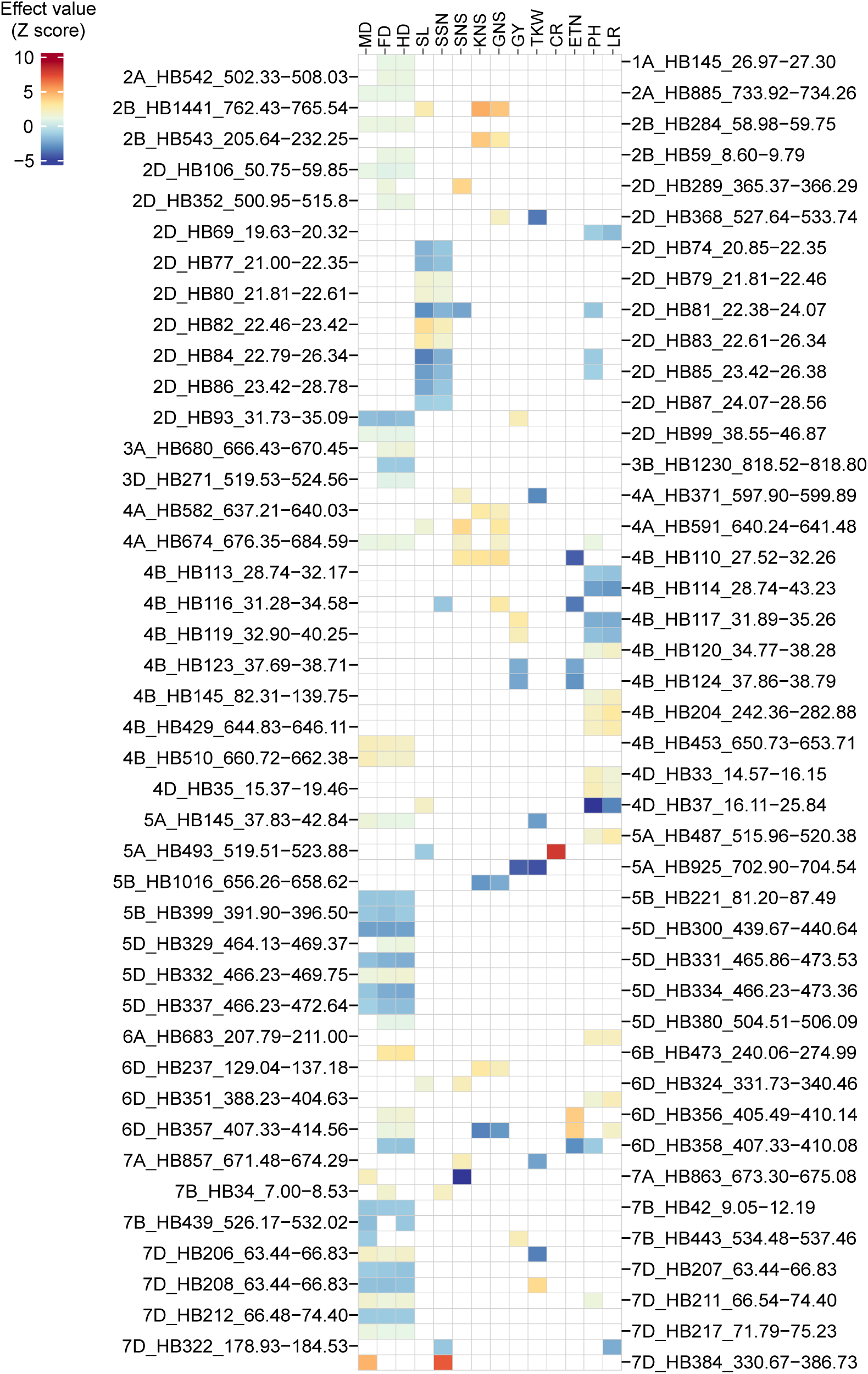
The frequency of favorable HBs across different era and subzone groups. This figure shows some of the results of the analysis, and the complete results are shown in Supplementary Table 17. Favorable HB is defined as follows: for traits such as plant height (PH), sterile spikelet number (SSN), cold resistance (CR), lodging resistance (LR), heading date (HD), flowering date (FD), and maturation date (MD), HB with a negative effect value (beta) indicates that this HB is beneficial. This corresponds to a shortening in plant height, fewer sterile spikelet, enhanced cold and lodging resistance, and earlier heading, flowering, and maturation. On the contrary, for other traits, HB with a positive effect value indicates that this HB is favorable. The position (Mb) of each HB on the chromosome are provided.

**Supplementary Fig. 11.**
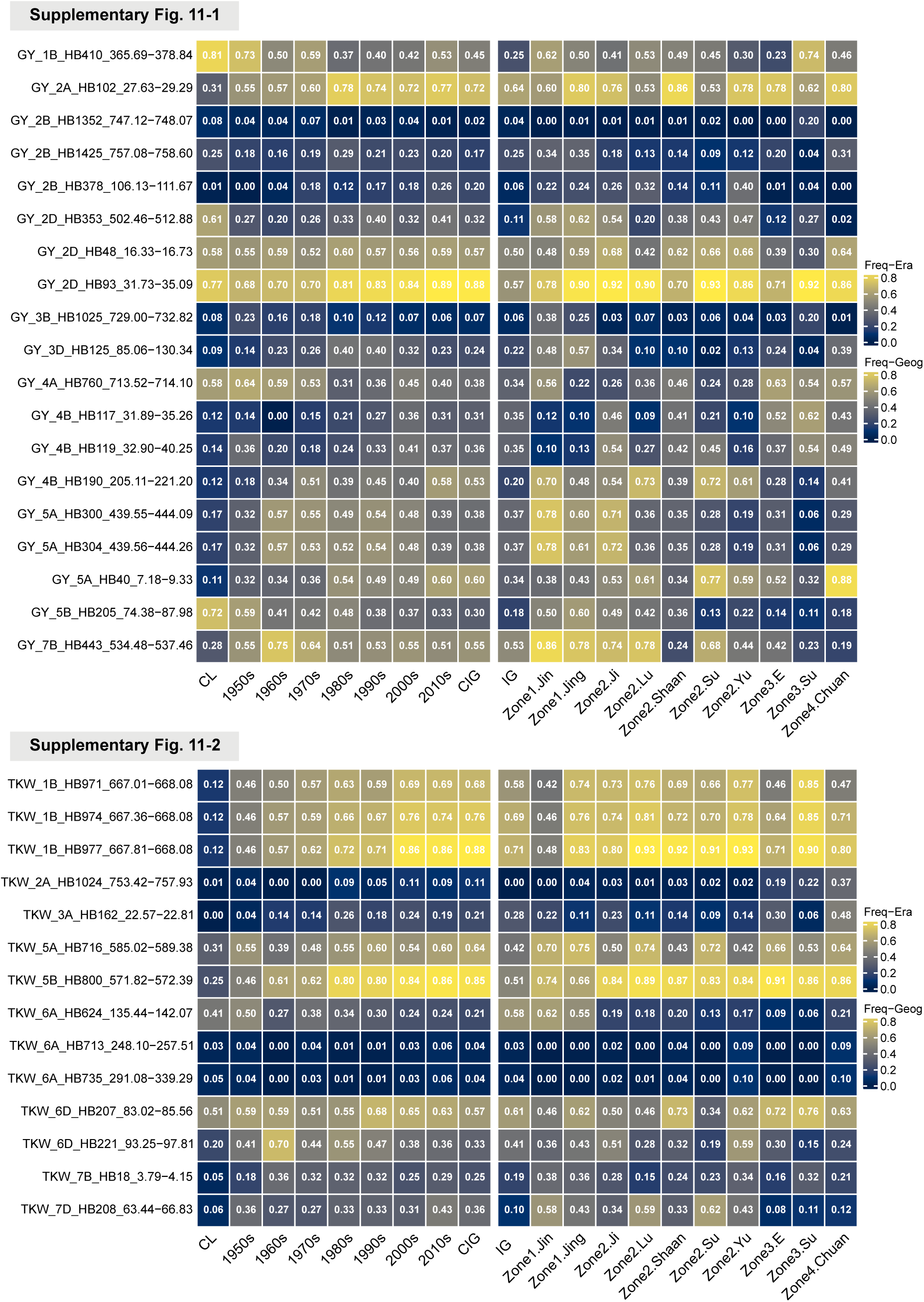
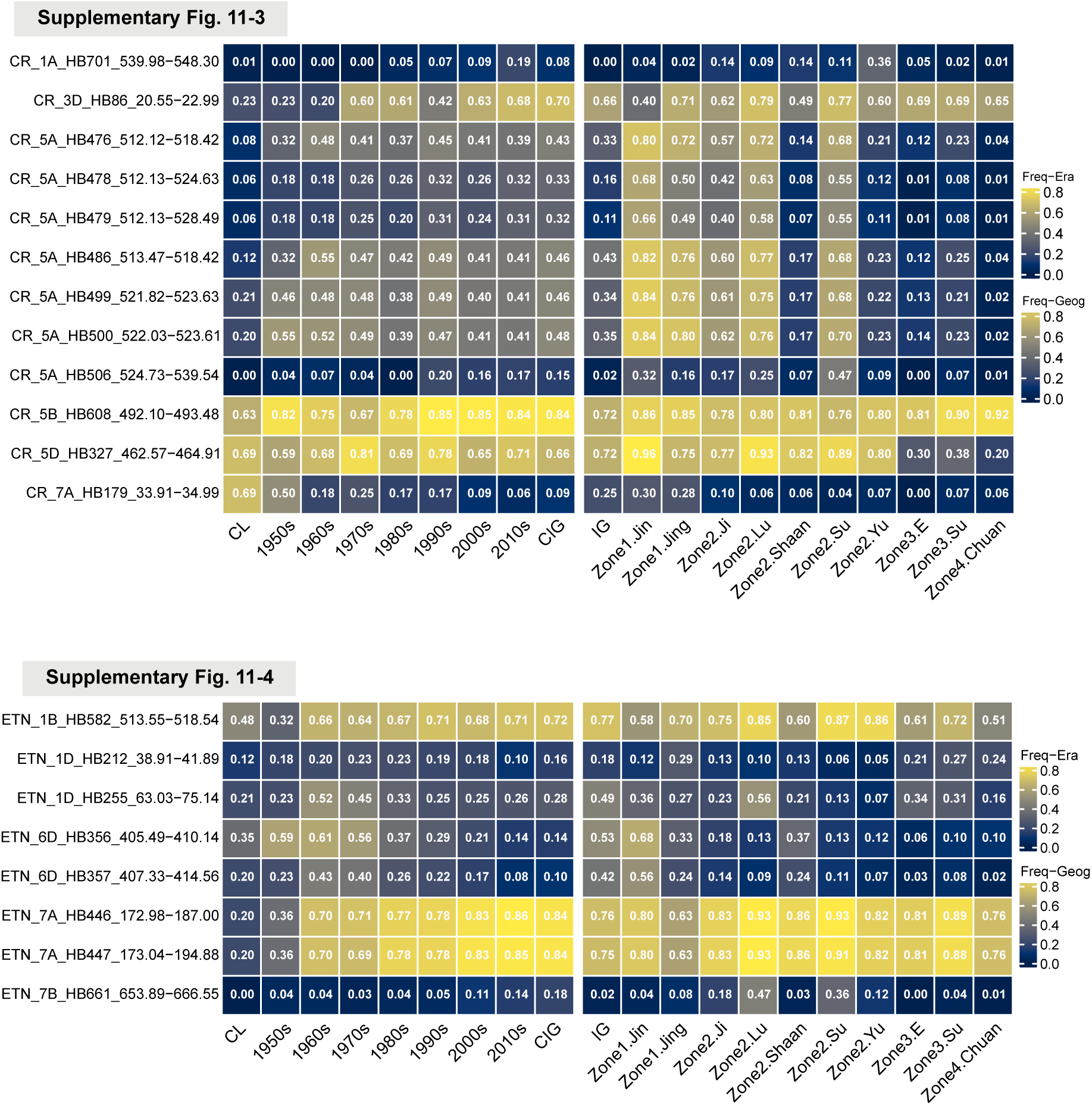
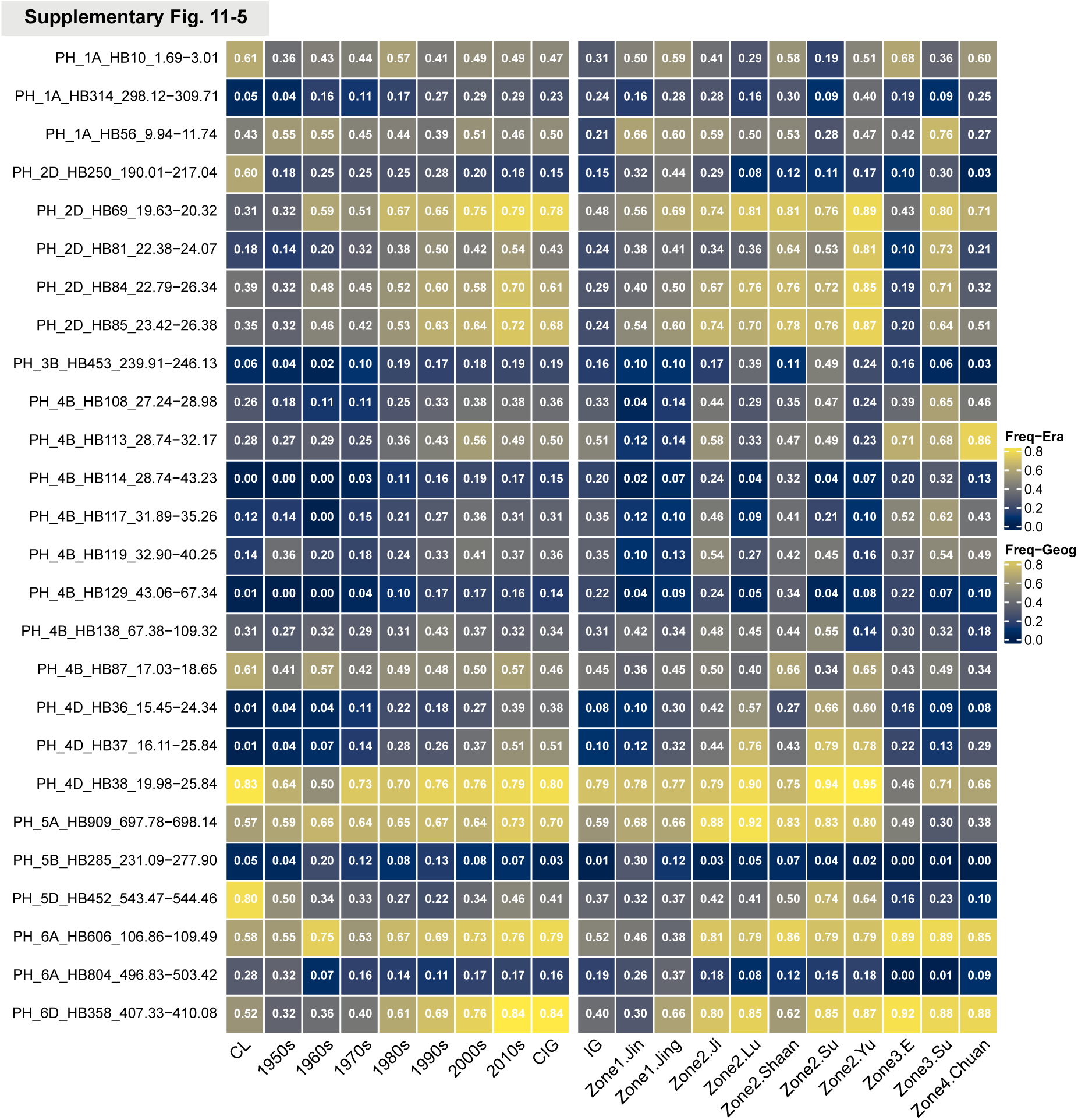
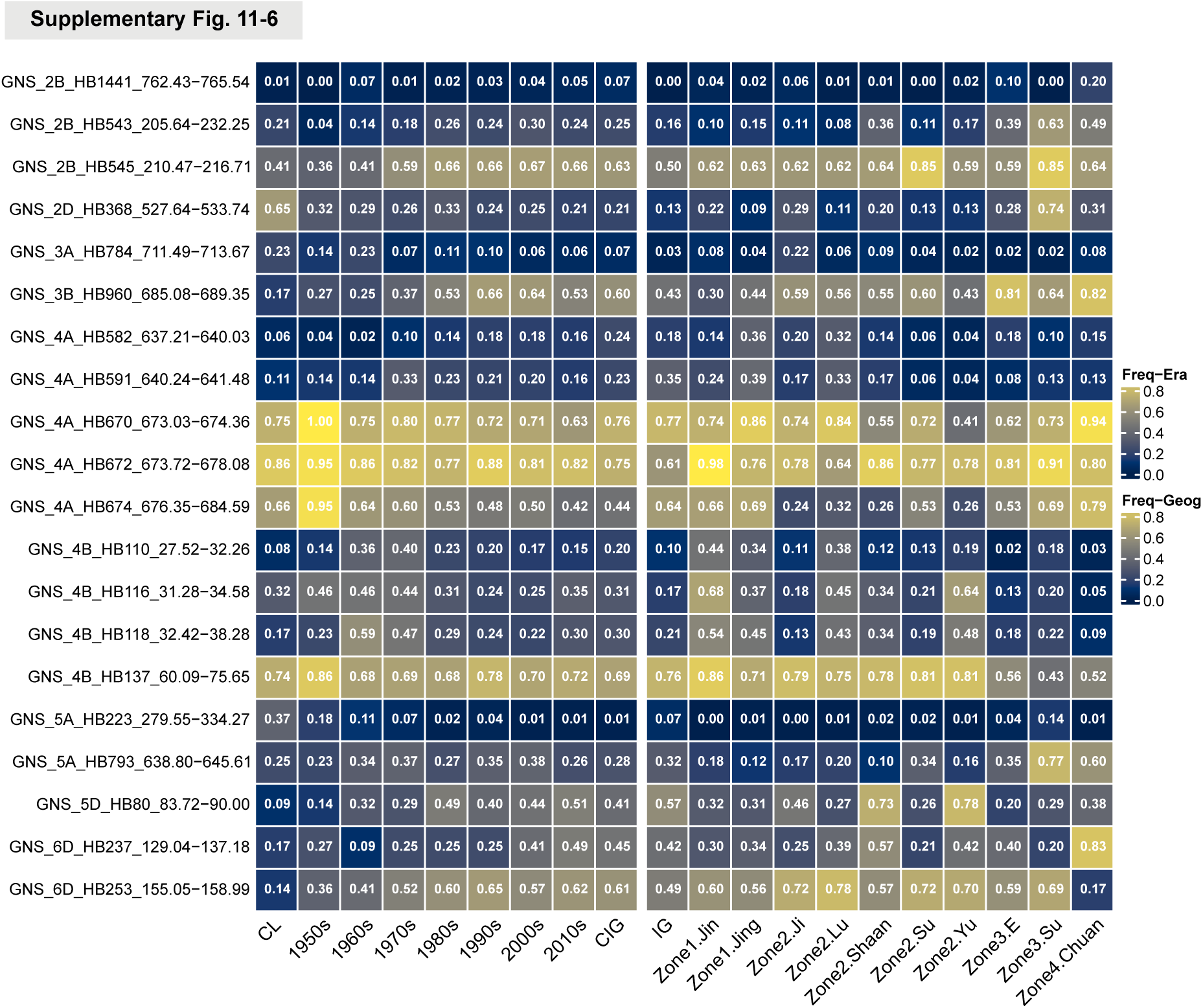
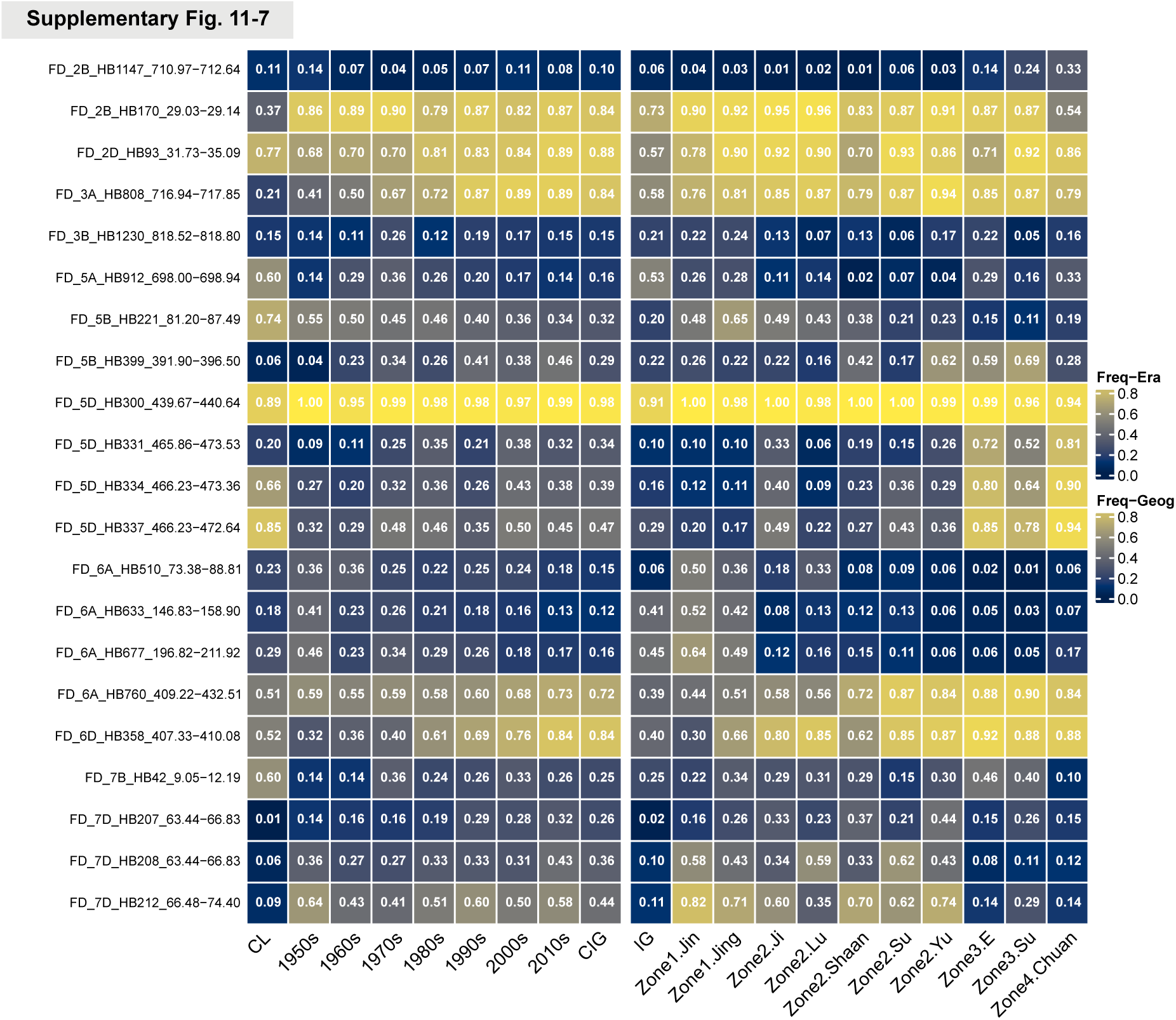
HBs associated with multiple traits. The effects were beta values calculated by GCTA, and the effect values were standardized using Z-scores to facilitate comparison across different traits. The raw values are also provided in Supplementary Table 17.

**Supplementary Fig. 12.**
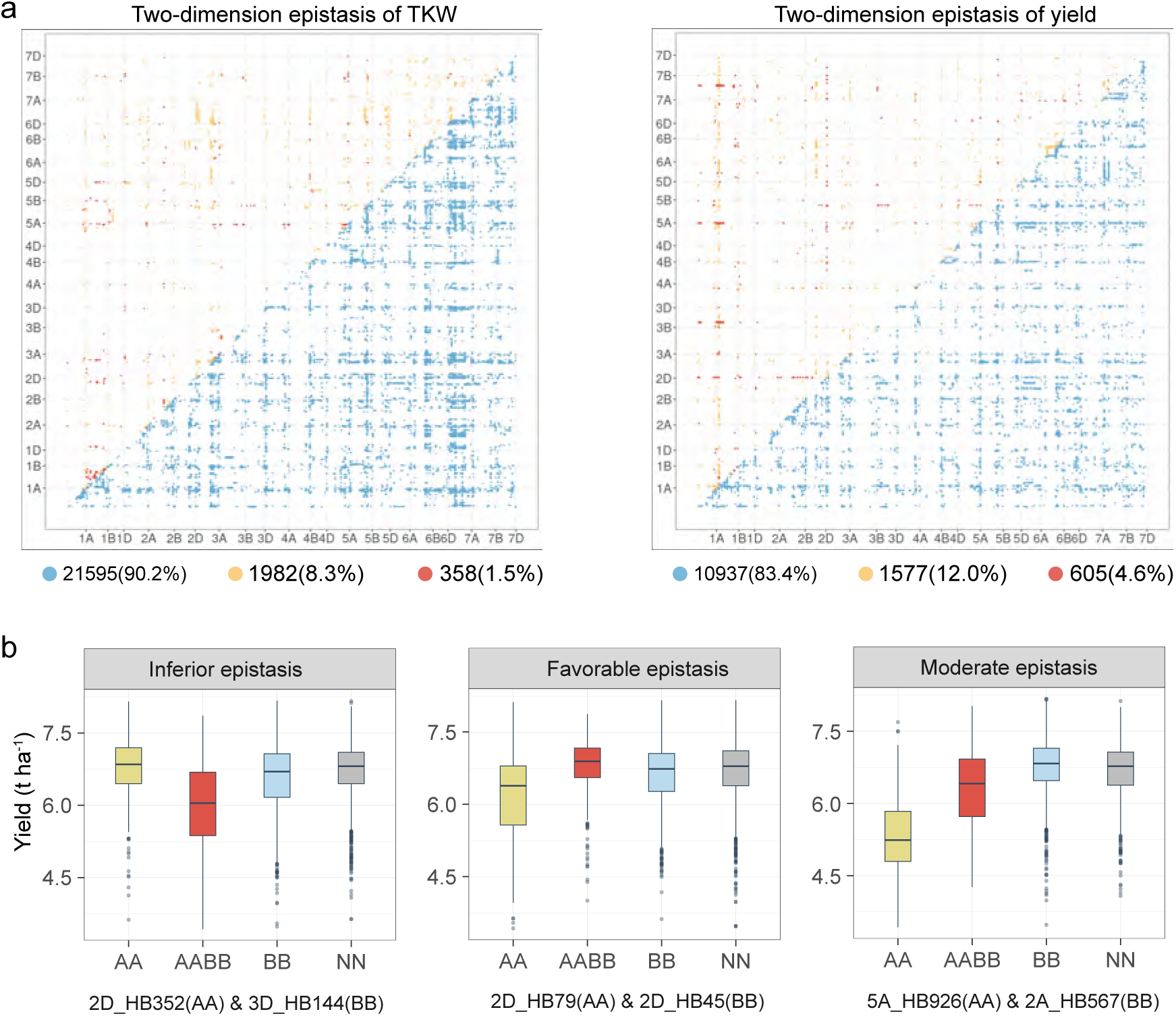
Two-dimension epistatic detection for TKW and grain yield. **a,** HB-HB interactions (epistasis) significantly associated with TKW (left) and grain yield (right) (*P* < 1×10^−16^). **b,** Classification of epistasis base on interaction effects. Three examples of how to classify epistasis based on phenotypic effect were shown. If the yield of accessions carrying a HB-HB interaction (AABB) is significantly higher than that of accessions without the interaction (AA and BB), this HB-HB interaction (AABB) is considered favorable epistasis, and conversely inferior or moderate epistasis.

**Supplementary Fig. 13.**
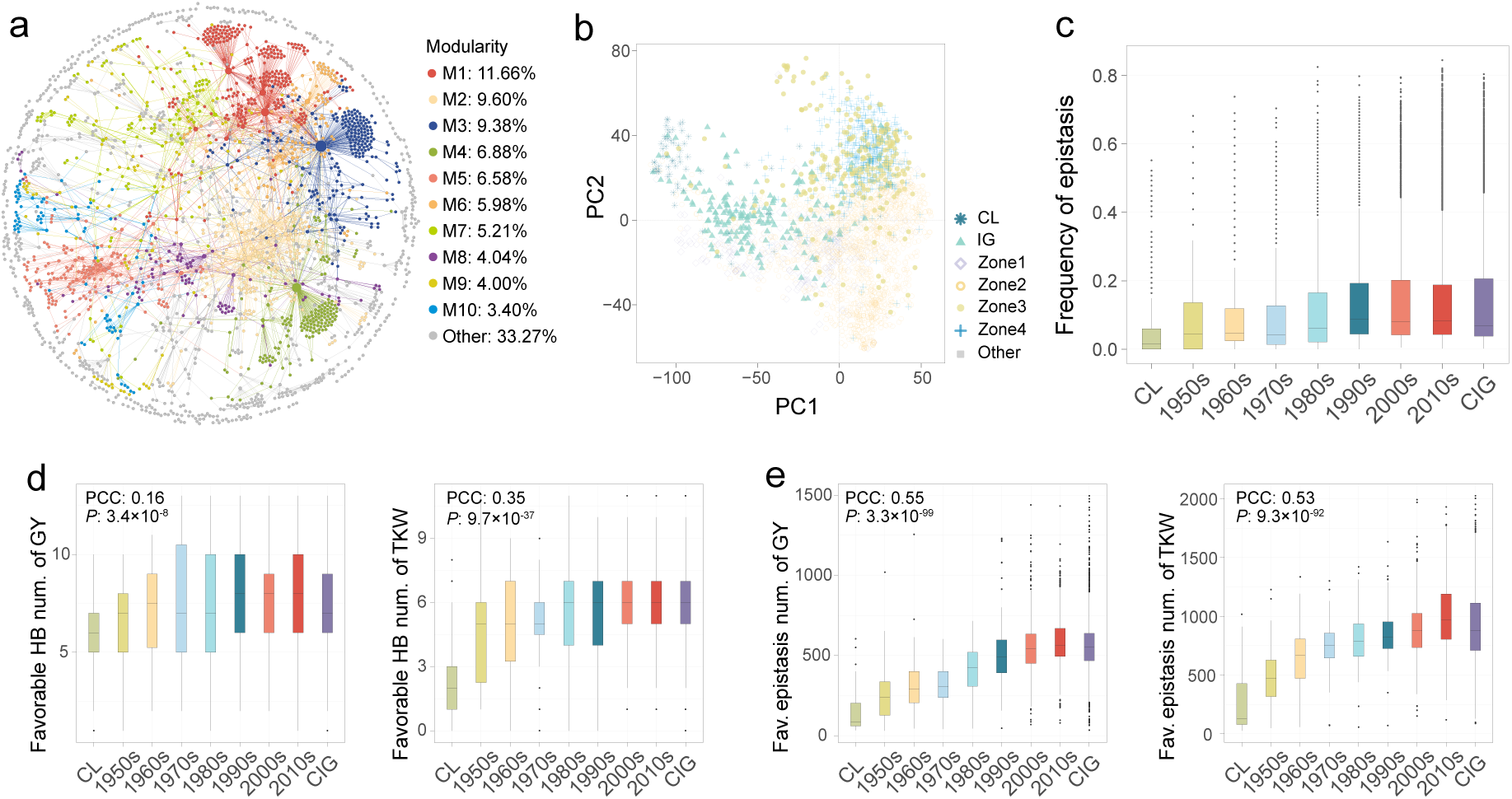
Epistasis network and epistasis patterns across different wheat groups. **a,** The network composed of favorable epistatic interactions. Each point represents an HB, and each edge indicates a favorable epistatic interaction between two HBs. The top 10 largest modules in the network are colored. The percentage of nodes in each module to the total number of nodes in the network is calculated. **b**, PCA based on the epistasis presence-absence matrix. Each point represents a sample and is colored according to its group. **c**, The frequency of favorable epistasis in different era groups. Each point represents an epistasis, and boxplots were plotted based on era groupings. The frequency of these epistasis significantly increased from landraces to early cultivars and current cultivars. **d**-**e**, Number of favorable HB (**d**) and epistasis (**e**) carried by wheat lines from different eras, including grain yield (left) and TKW (right). Each point represents a sample, and boxplots were plotted based on era groupings. Pearson correlation coefficients and *P*-values were calculated. The calculation of correlation excluded the CIG group.

**Supplementary Fig. 14.**
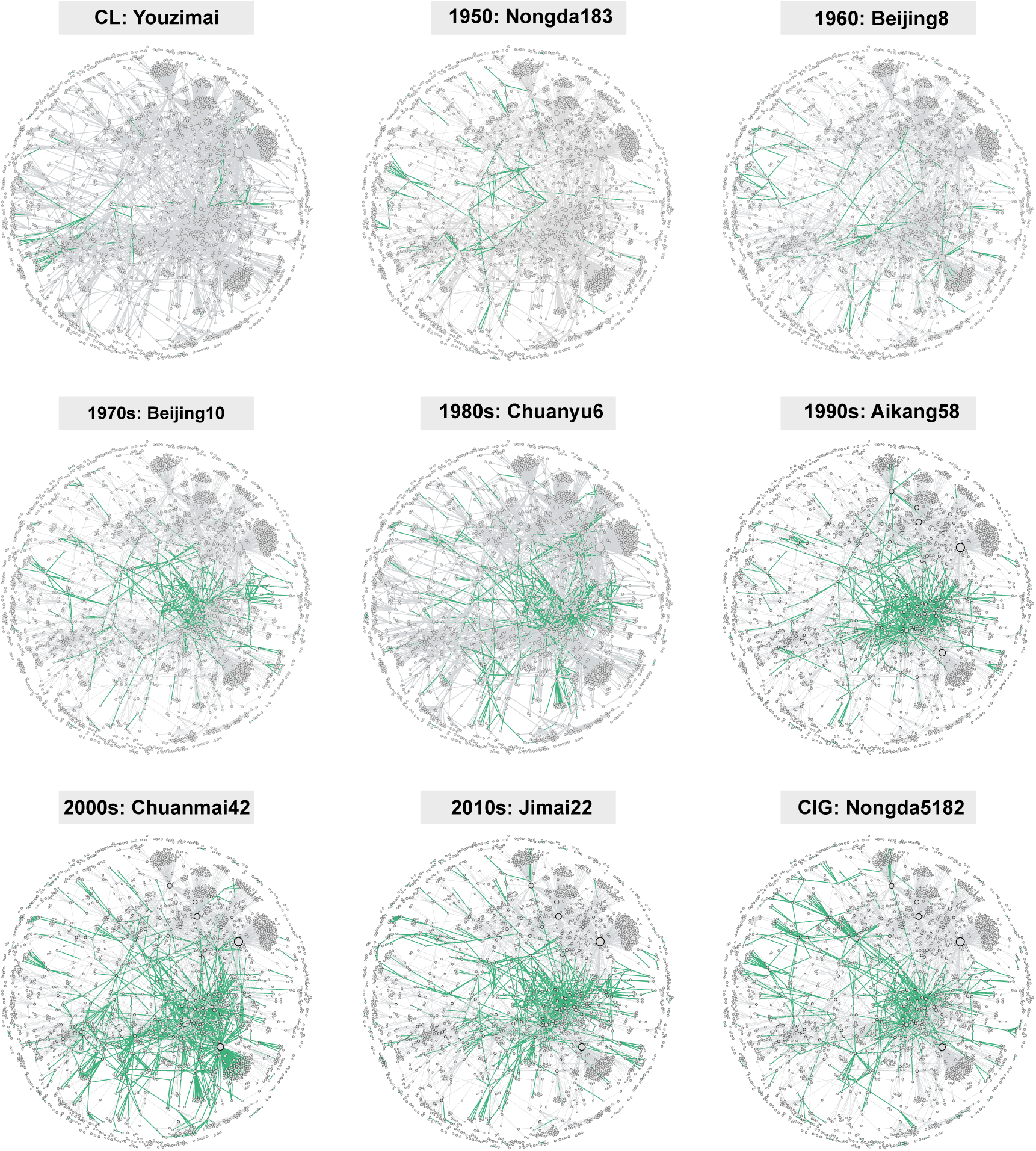
The favorable epistasis carried by wheat lines from different eras. Each point represents an HB, and each edge indicates a favorable epistatic interaction between two HBs. The green edge represents the presence of this epistasis in the wheat line, while the gray edge indicates its absence in the line. A representative wheat line was selected from each era.

**Supplementary Fig. 15.**
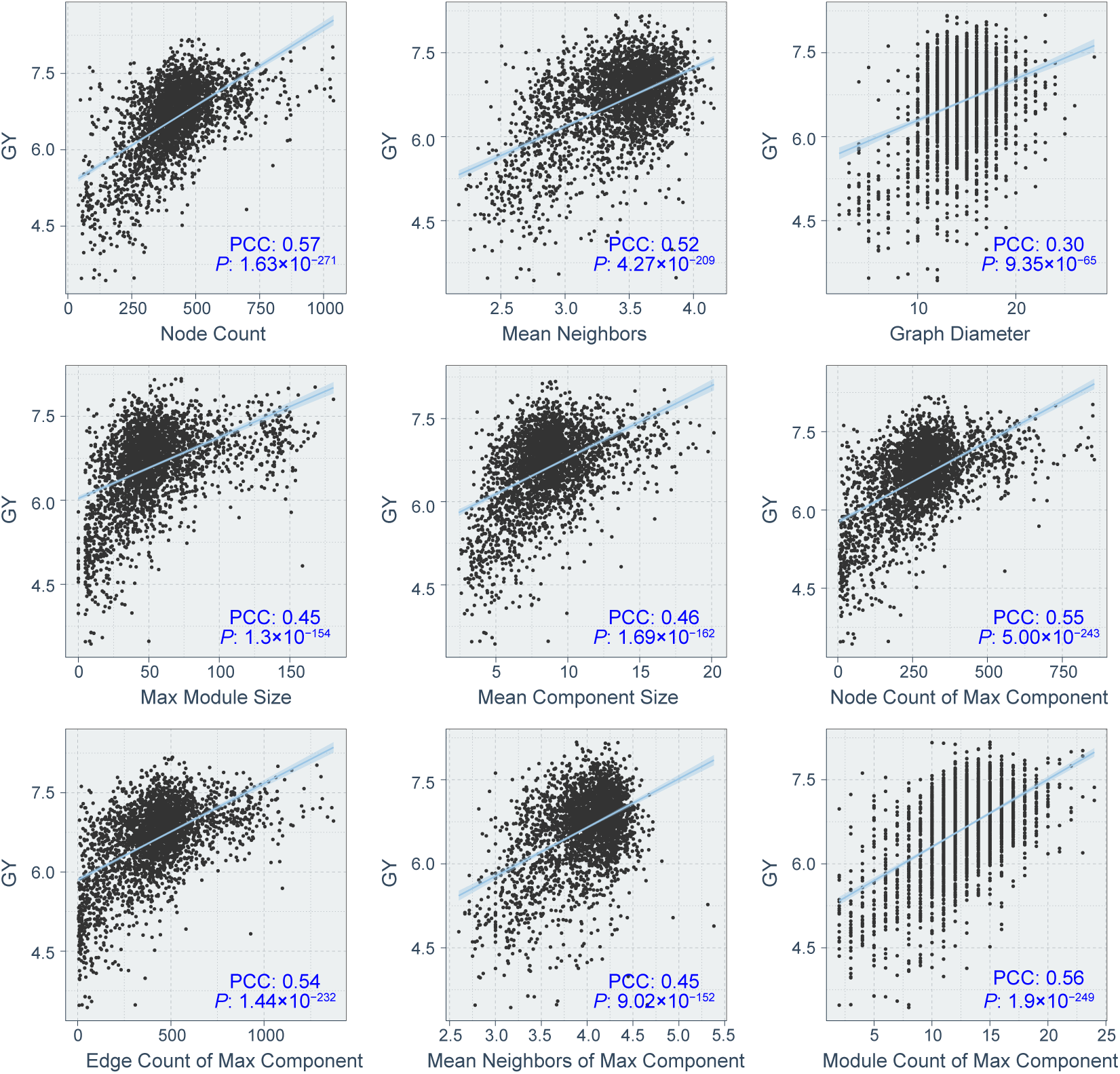
The correlation between the topological properties of epistatic network and yield. Each point represents a sample. Pearson correlation coefficients and *P*-values were calculated. A linear method was used for fitting.

**Supplementary Fig. 16.**
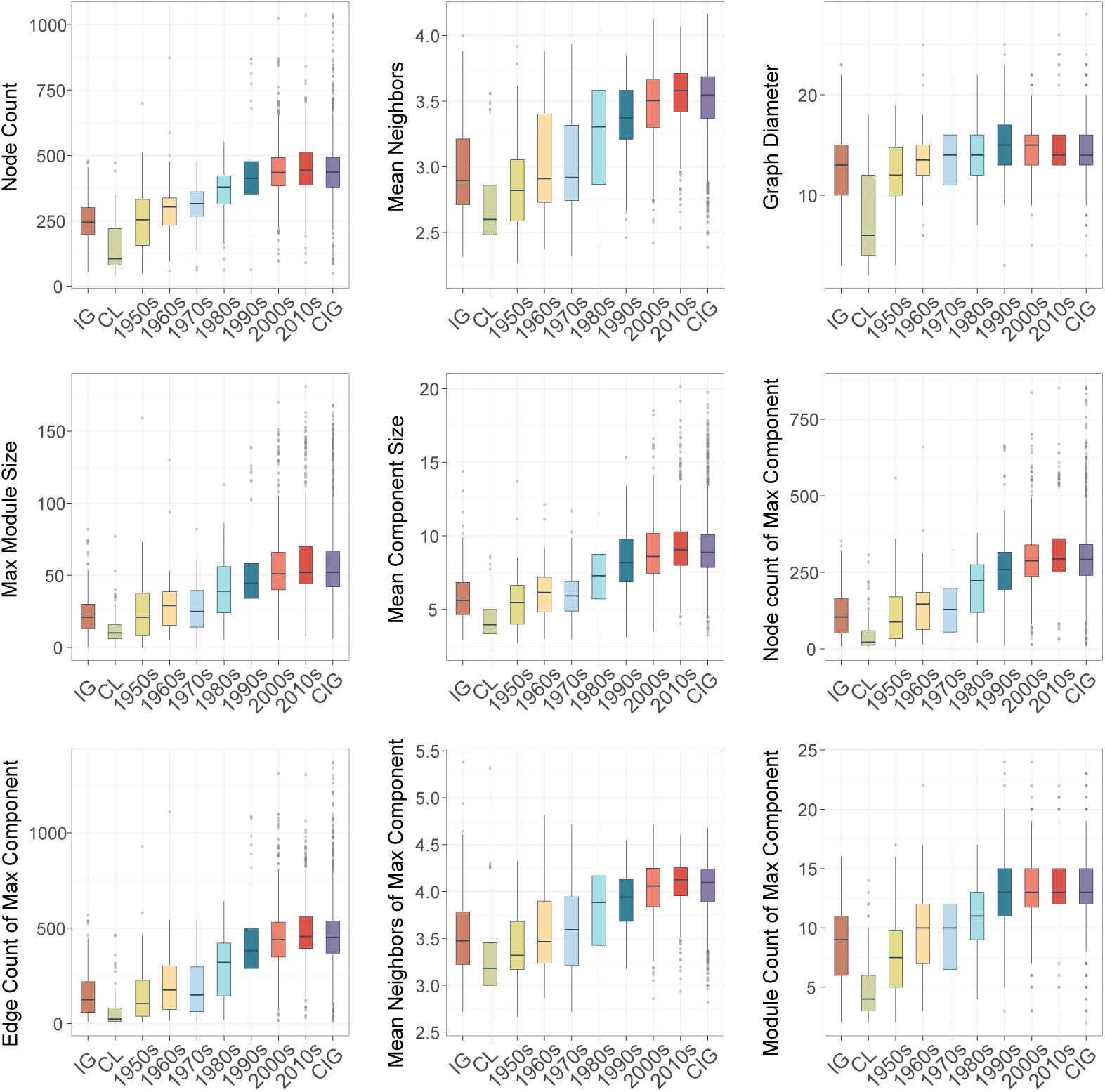
Topological properties of epistatic network of wheat lines across different era groups. Each point represents a sample, and boxplots were plotted based on era groupings.

**Supplementary Fig. 17.**
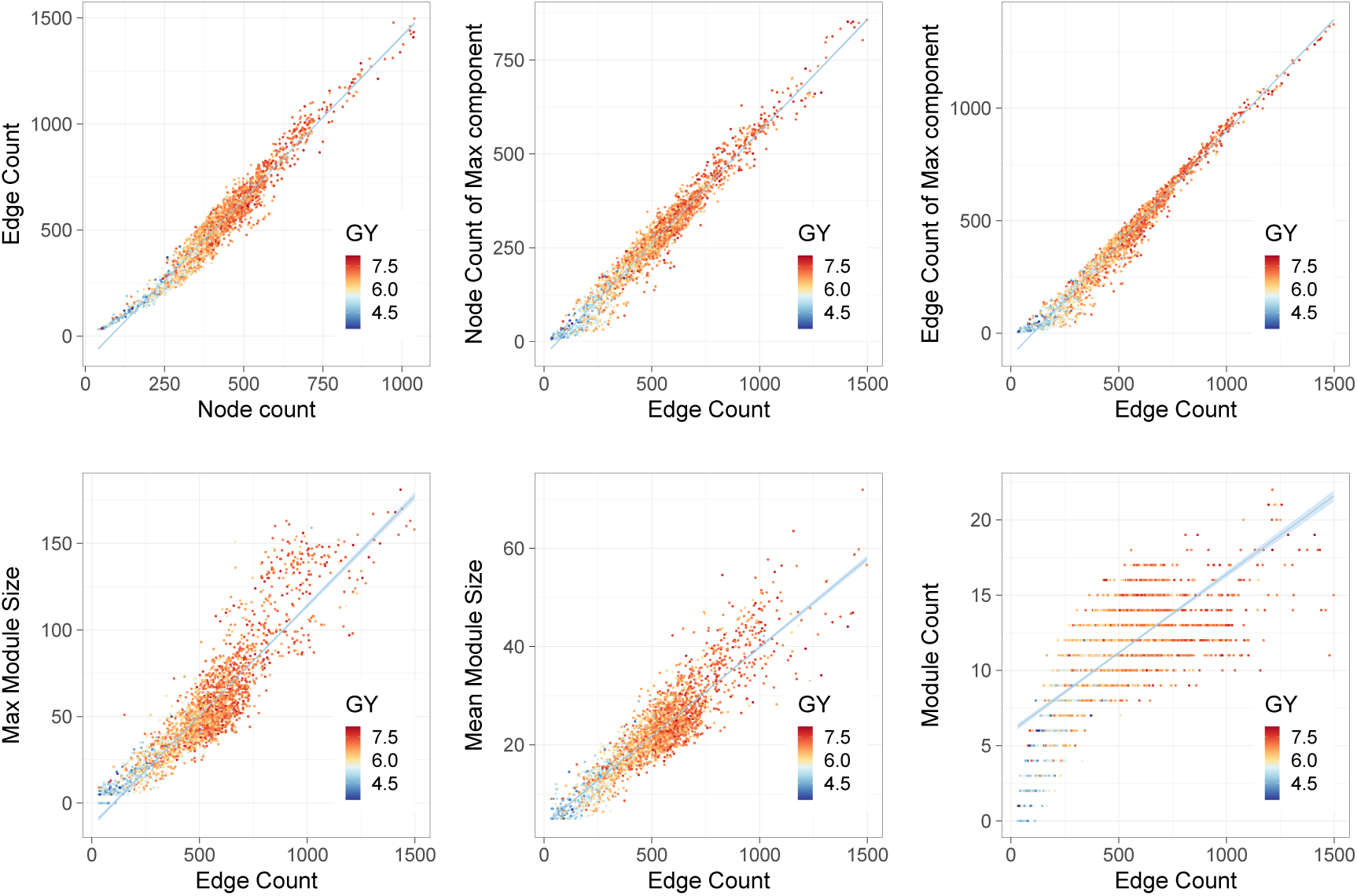
The pattern of network expansion. Each point represents a wheat line, and its yield value is used for coloring. As the nodes or edges of the network expand, properties such as the size and number of modules increase linearly. A linear method was used for fitting. Pearson correlation coefficients and *P*-values were calculated.

**Supplementary Fig. 18.**
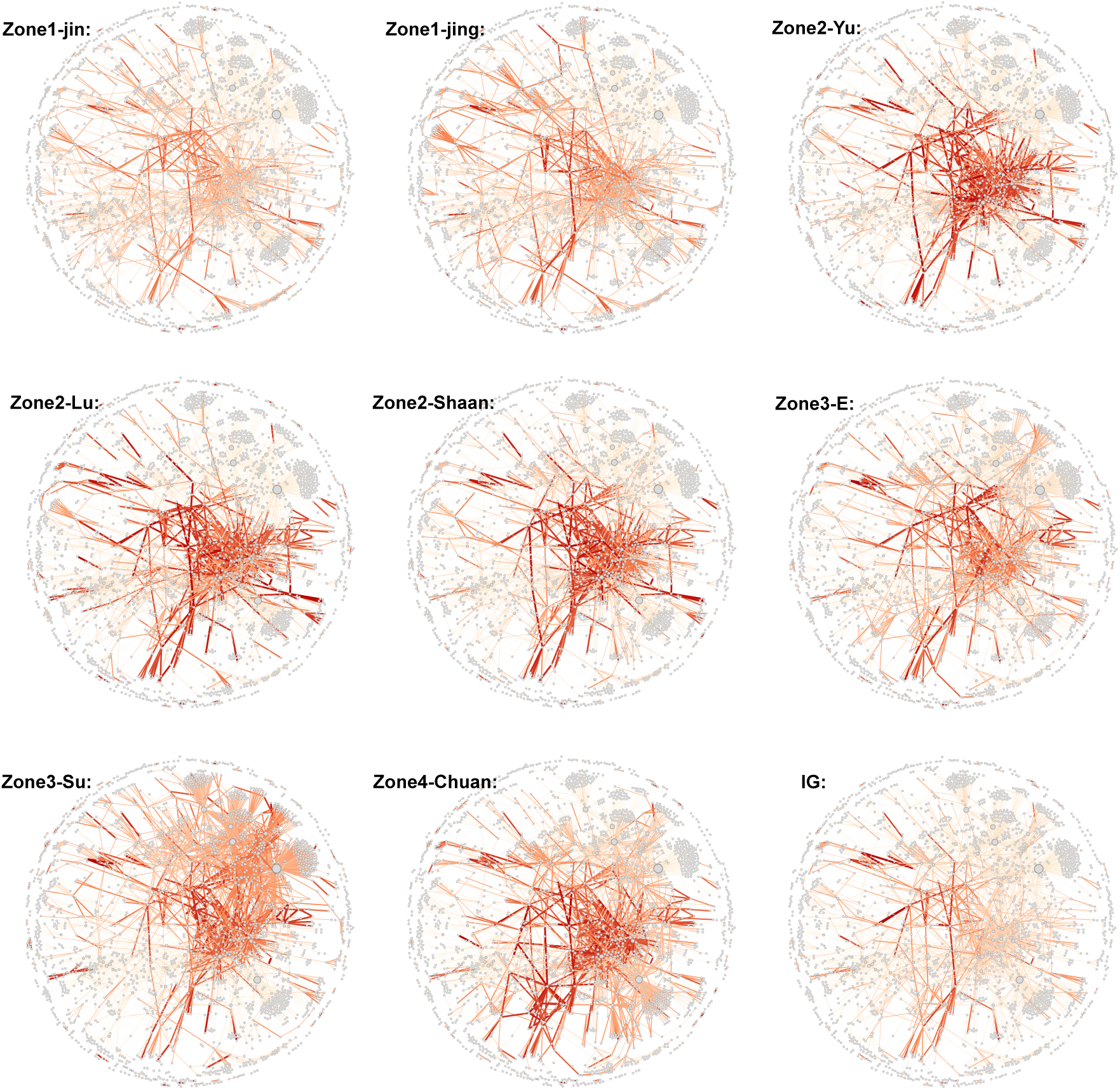
Favorable epistasis networks across different subzone groups. Each node represents an HB, and each edge indicates the epistasis between two HBs. The edge thickness and color intensity reflect the frequency of epistasis in the group, with thicker and redder edges indicating higher frequency.

**Supplementary Fig. 19.**
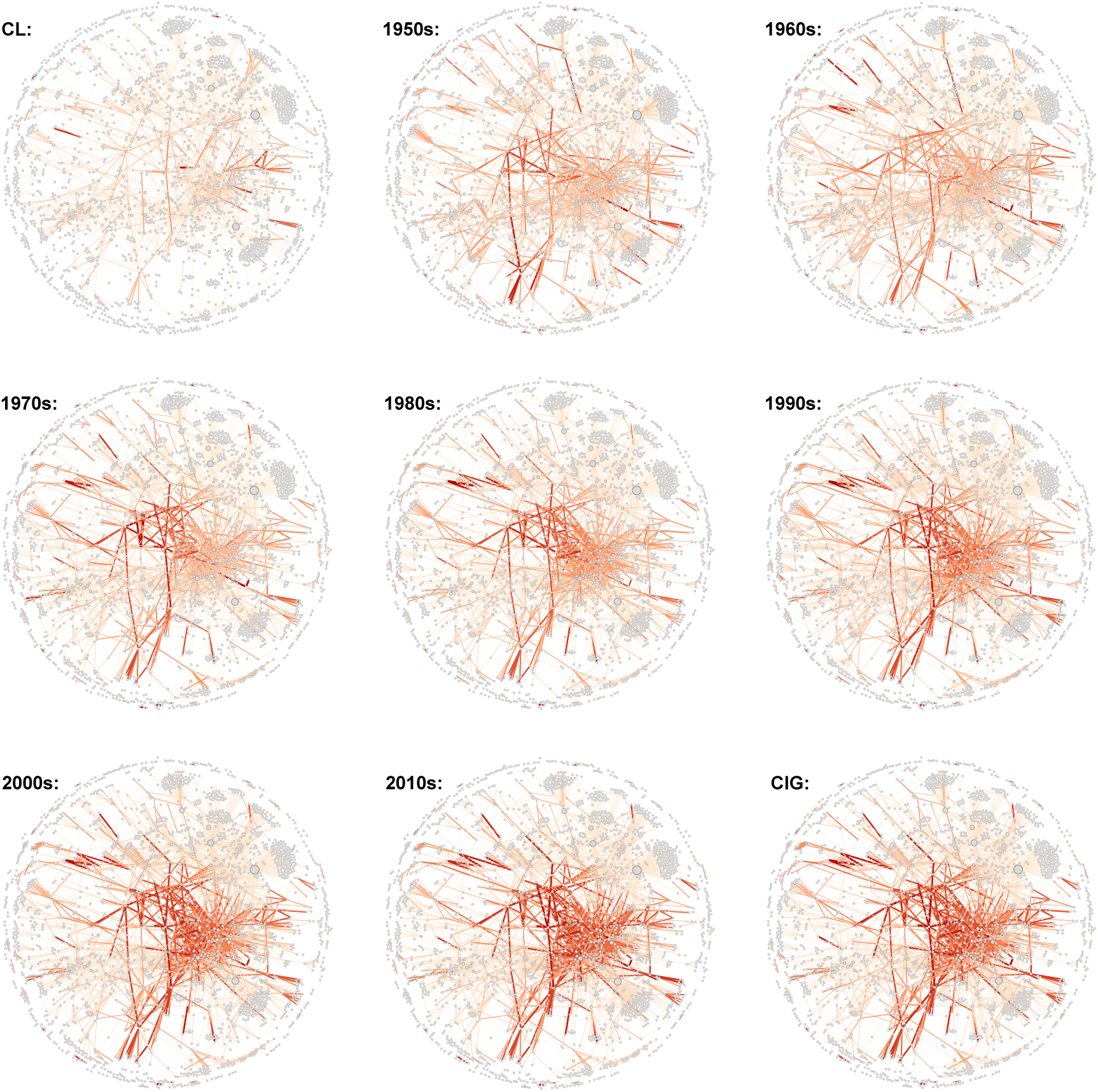
Favorable epistasis networks across different era groups. Each node represents an HB, and each edge indicates the epistasis between two HBs. The edge thickness and color intensity reflect the frequency of epistasis in the group, with thicker and redder edges indicating higher frequency.

**Supplementary Fig. 20.**
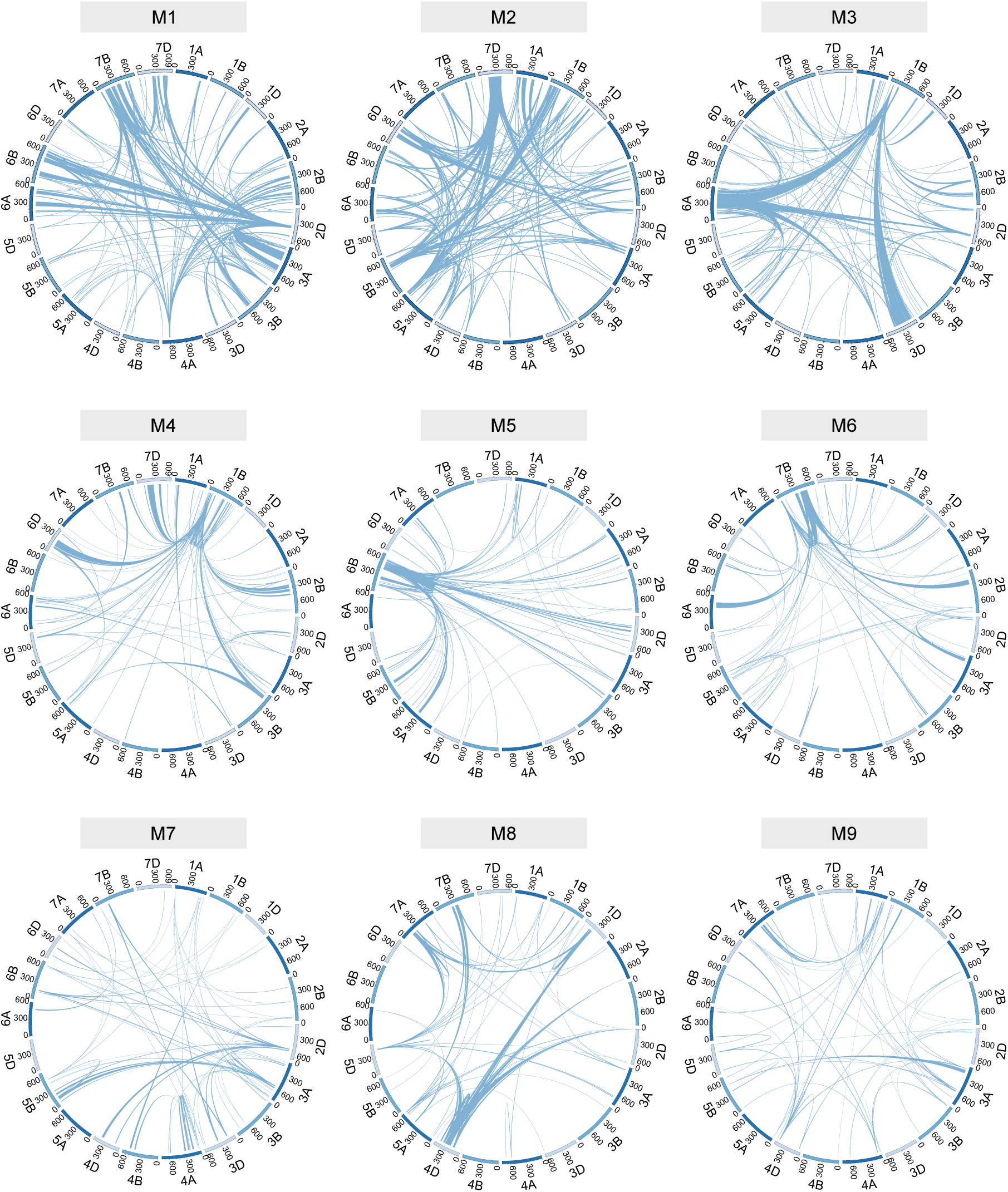
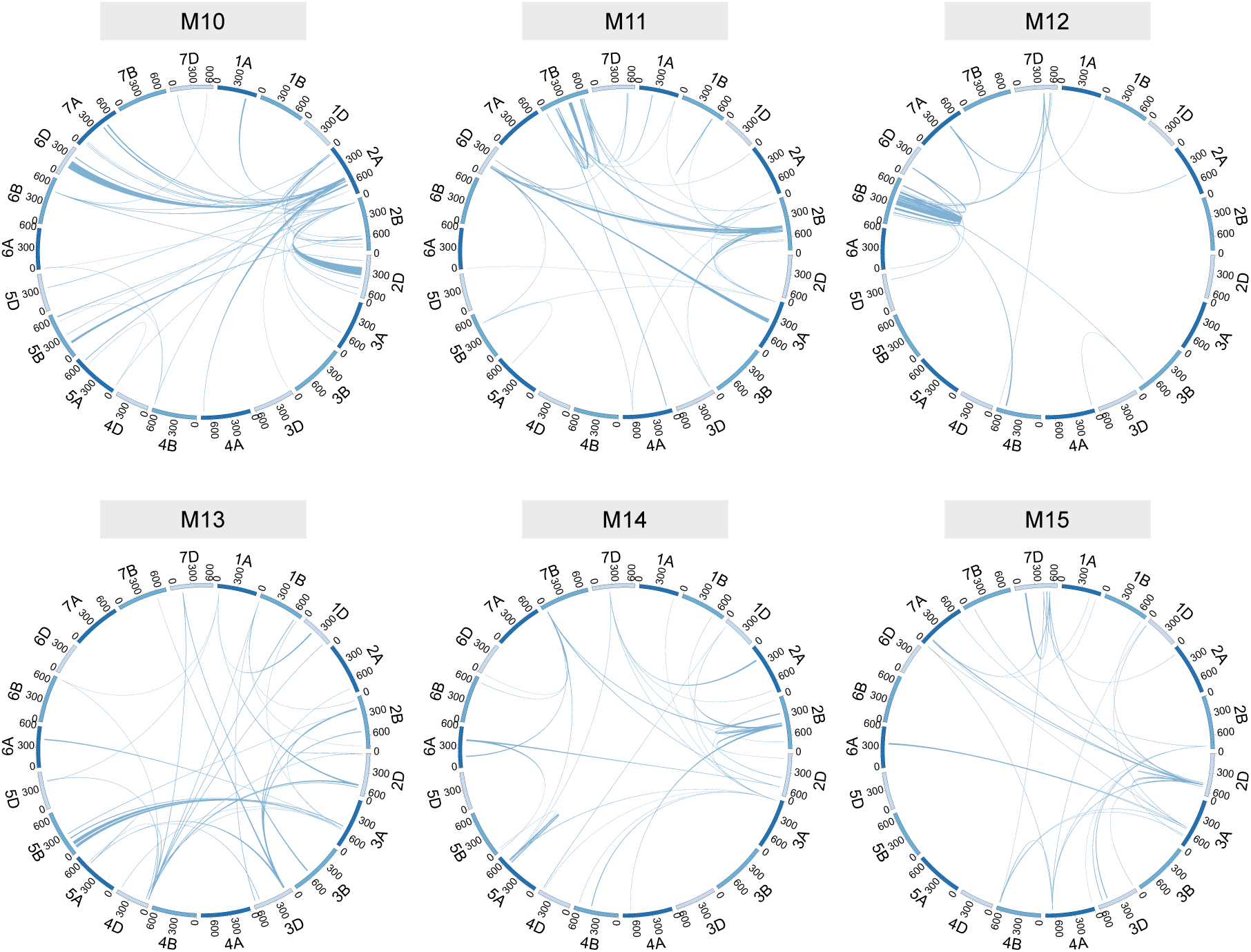
Circular link plots of the top 15 largest modules in the network. Circular link plots reflect the genomic locations of these epistasis and modules. Epistasis between HBs is represented by links. Since an HB corresponds to a genomic segment, all links are shown with a width.

**Supplementary Fig. 21.**
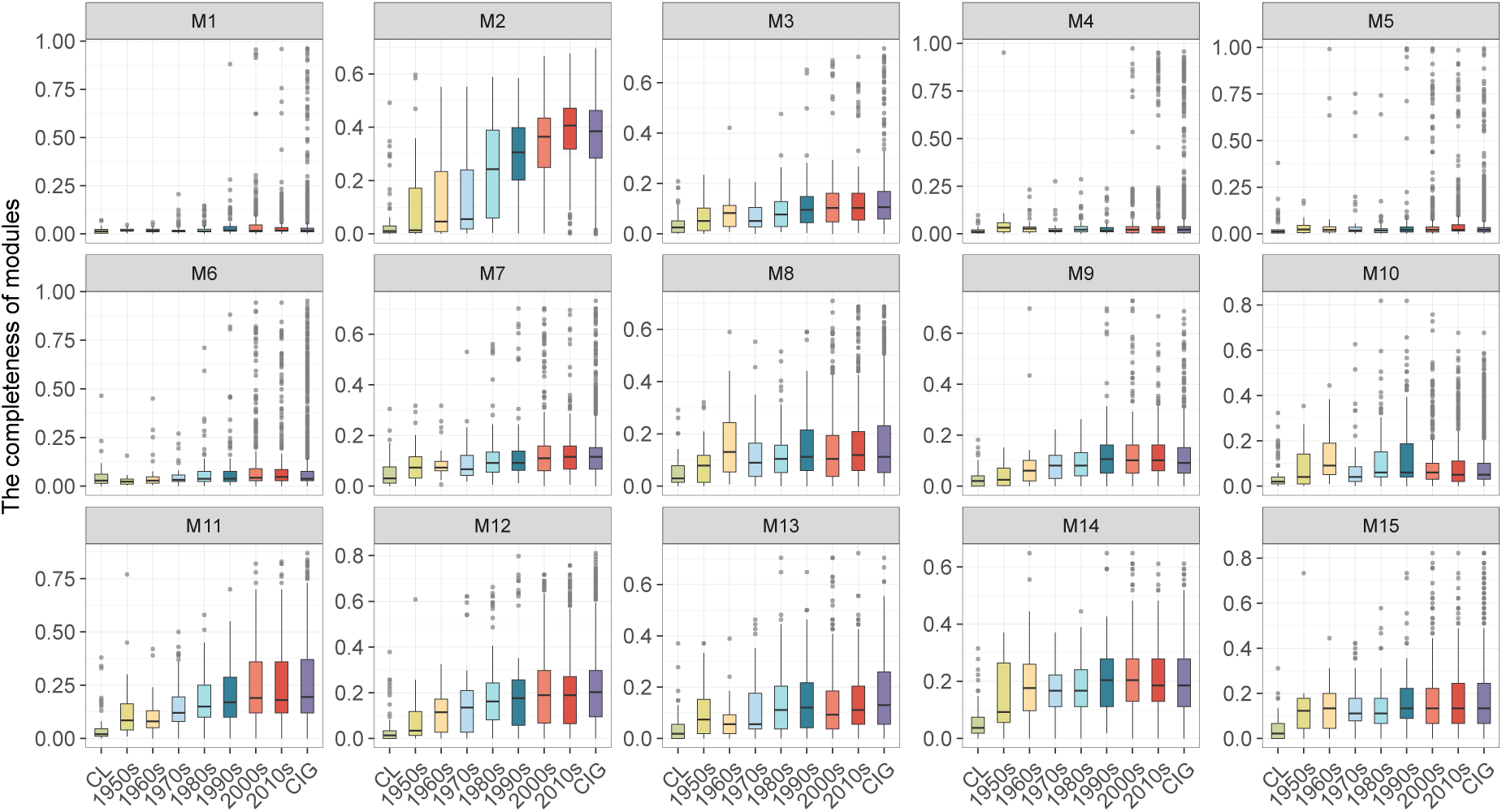
The completeness of the top 15 largest modules in the network changes over time. Each point represents a sample, and boxplots were plotted based on era groupings. The completeness of each module is calculated by dividing the number of edges in the module carried by each wheat line by the total number of edges in the module.

**Supplementary Fig. 22.**
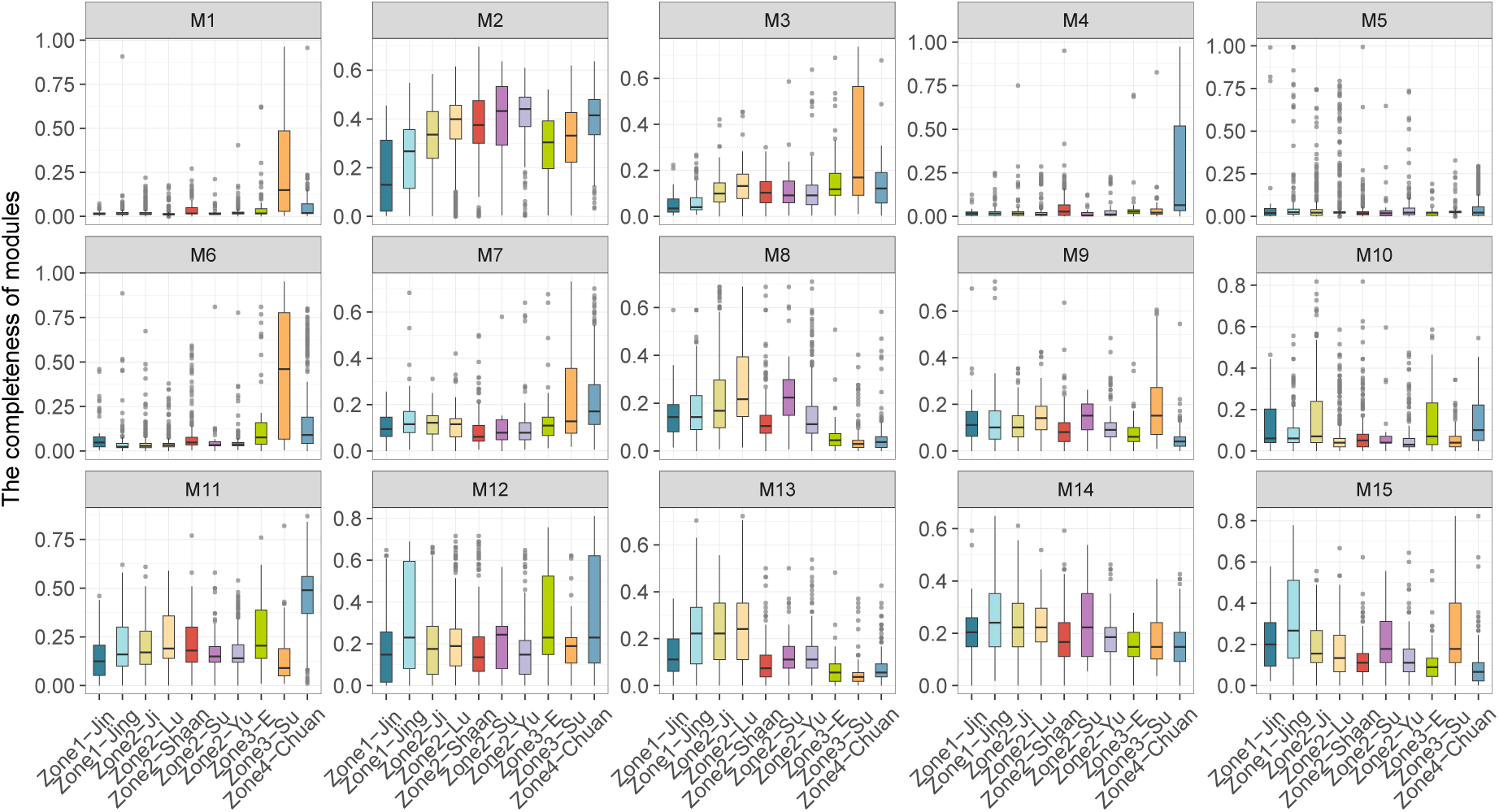
Comparison of module completeness of wheat lines across different subzones. Each point represents a sample, and boxplots were plotted based on subzone groupings. The completeness of each module is calculated by dividing the number of edges in the module carried by each wheat line by the total number of edges in the module.

**Supplementary Fig. 23.**
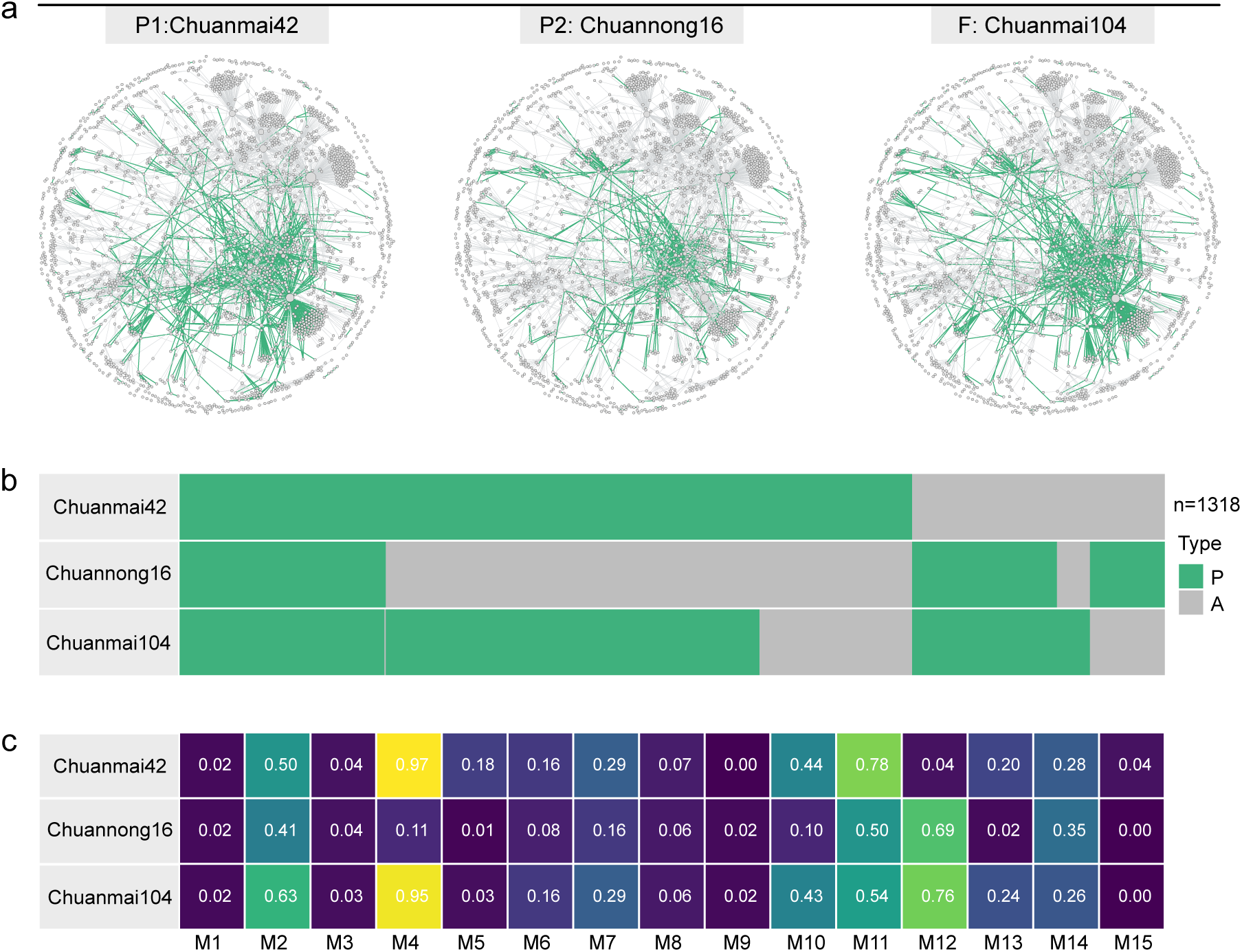
Inheriting and forming of favorable epistatic modules in breeding lineages. **a**, Epistatic networks of pedigree members. The pedigree includes ‘Chuanmai 42’ and ‘Chuannong 16’ as parents, and ‘Chuanmai 104’ as the progeny. Each point represents an HB, and each edge indicates a favorable epistatic interaction between two HBs. The green edges represent the presence of these epistasis in the wheat line, while the gray edges indicate their absence in the line. **b**, The presence-absence matrix of epistasis. Each column represents a favorable epistasis, with a total of 1,318 columns. The green columns represent the presence of these epistasis in the wheat line, while the gray columns indicate their absence in the line. This diagram clearly shows which epistases are shared between parents and progeny, and which are specific to individual genotypes. **c**, Completeness of the top 15 largest modules that were identified based on all accessions. The completeness of each module is calculated by dividing the number of edges in the module carried by each wheat line by the total number of edges in the module.

**Supplementary Fig. 24.**
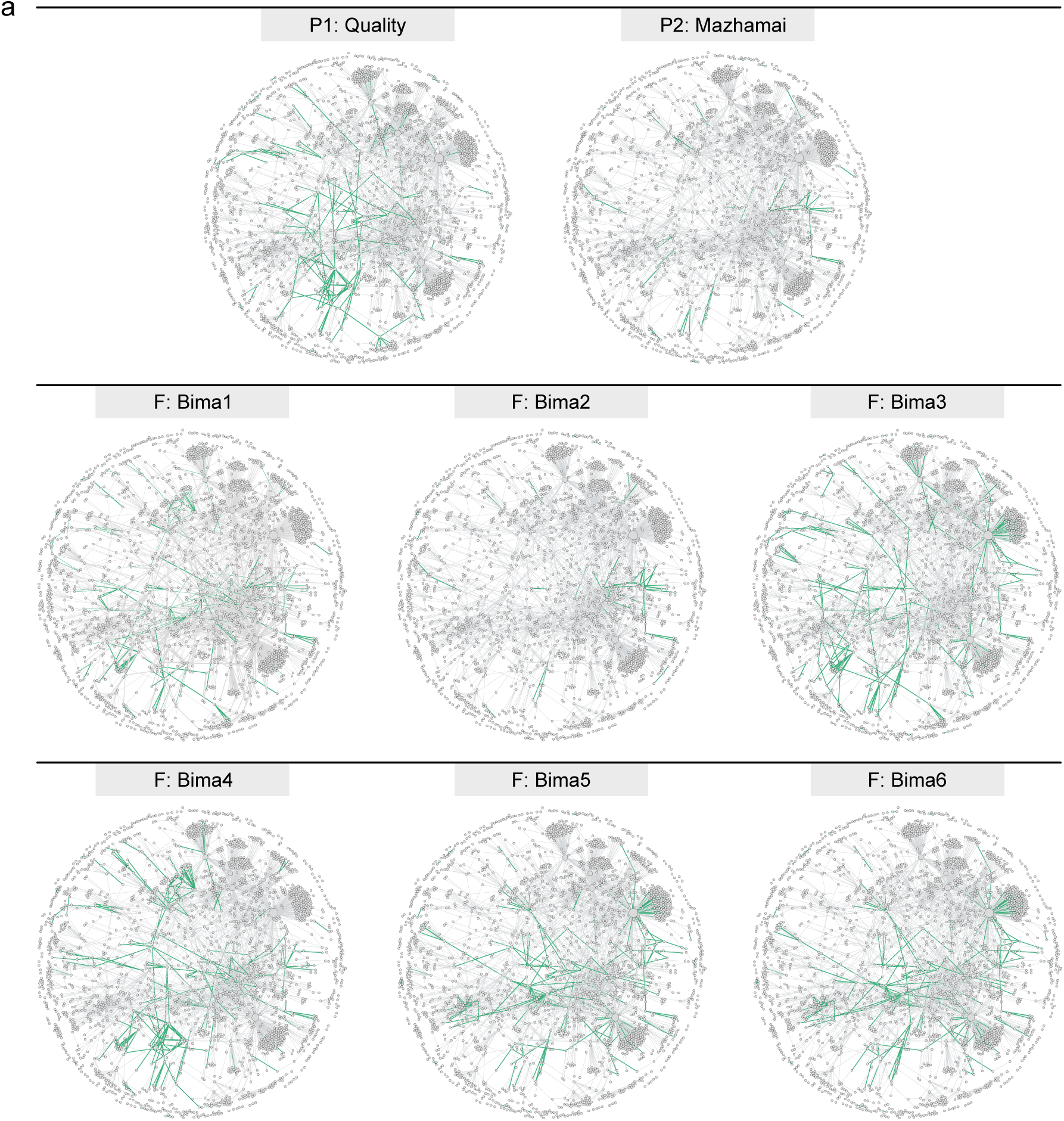
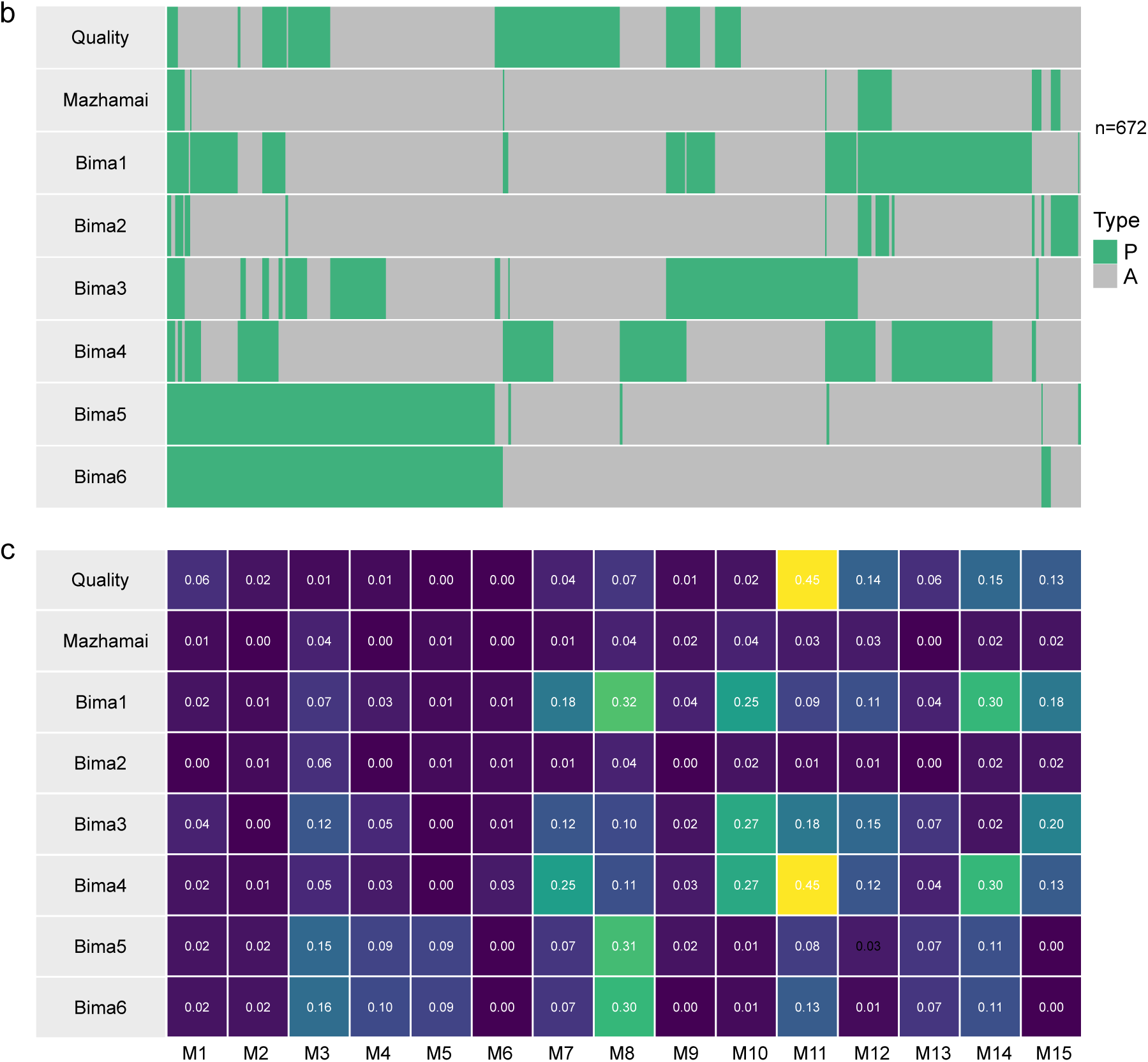
Inheriting and forming of favorable epistatic modules in breeding lineages. **a**, Epistatic networks of pedigree members. The pedigree includes ‘Quality’ and ‘Mazhamai’ as parents, and six lines of ‘Bima 1-6’ as the progenies. Each point represents an HB, and each edge indicates a favorable epistatic interaction between two HBs. The green edges represent the presence of these epistasis in the wheat line, while the gray edges indicate their absence in the line. **b**, The presence-absence matrix of epistasis. Each column represents a favorable epistasis, with a total of 672 columns. The green columns represent the presence of these epistasis in the wheat line, while the gray columns indicate their absence in the line. This diagram clearly shows which epistases are shared between parents and progeny, and which are specific to individual genotypes. **c**, Completeness of the top 15 largest modules that were identified based on all accessions. The completeness of each module is calculated by dividing the number of edges in the module carried by each wheat line by the total number of edges in the module.

**Supplementary Fig. 25.**
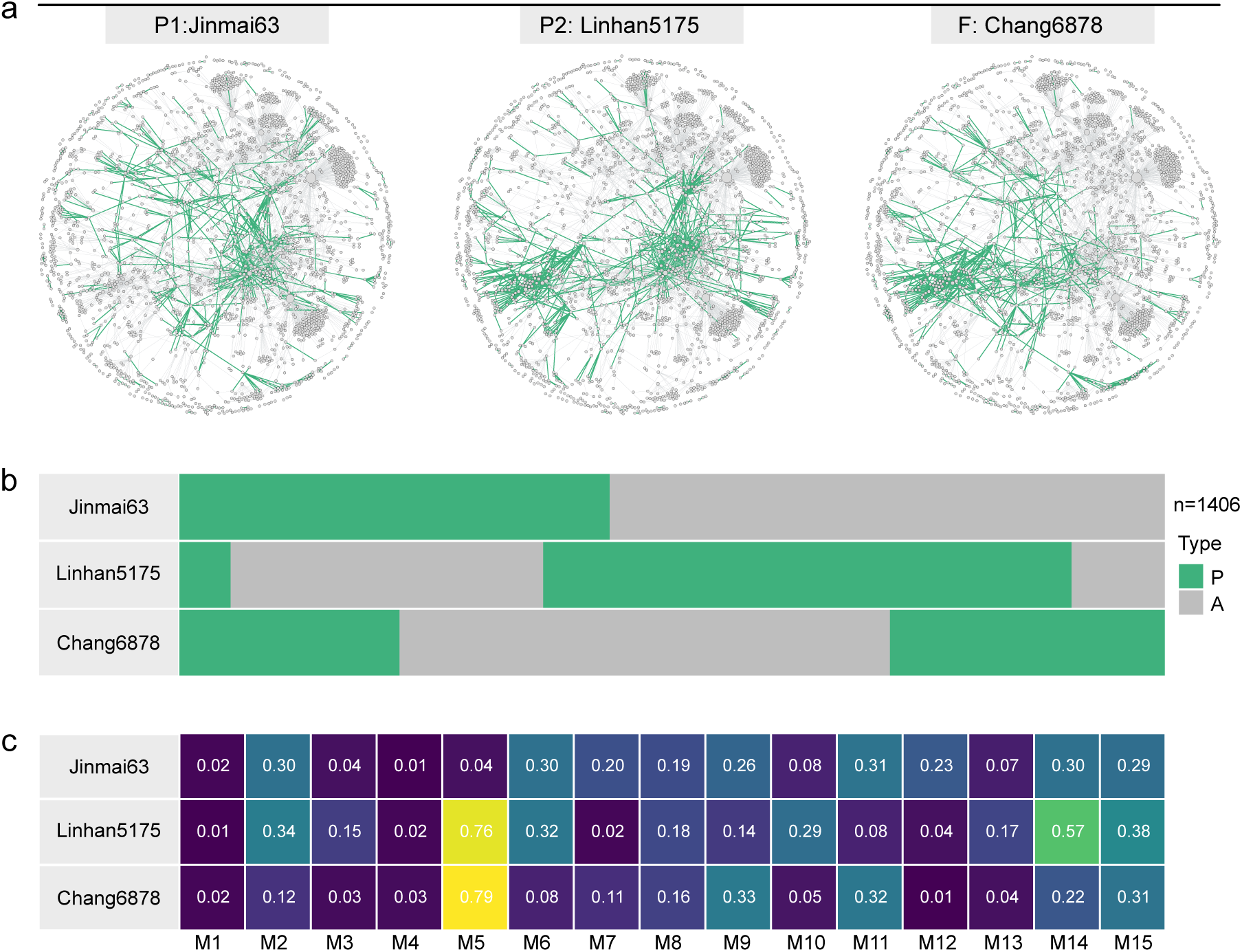
Inheriting and forming of favorable epistatic modules in breeding lineages. **a**, Epistatic networks of pedigree members. The pedigree includes ‘Jinmai 63’ and ‘Linhan 5175’ as parents, and ‘Chang 6878’ as the progenies. Each point represents an HB, and each edge indicates a favorable epistatic interaction between two HBs. The green edges represent the presence of these epistasis in the wheat line, while the gray edges indicate their absence in the line. **b**, The presence-absence matrix of epistasis. Each column represents a favorable epistasis, with a total of 1,406 columns. The green columns represent the presence of these epistasis in the wheat line, while the gray columns indicate their absence in the line. This diagram clearly shows which epistases are shared between parents and progeny, and which are specific to individual genotypes. **c**, Completeness of the top 15 largest modules that were identified based on all accessions. The completeness of each module is calculated by dividing the number of edges in the module carried by each wheat line by the total number of edges in the module.

**Supplementary Fig. 26.**
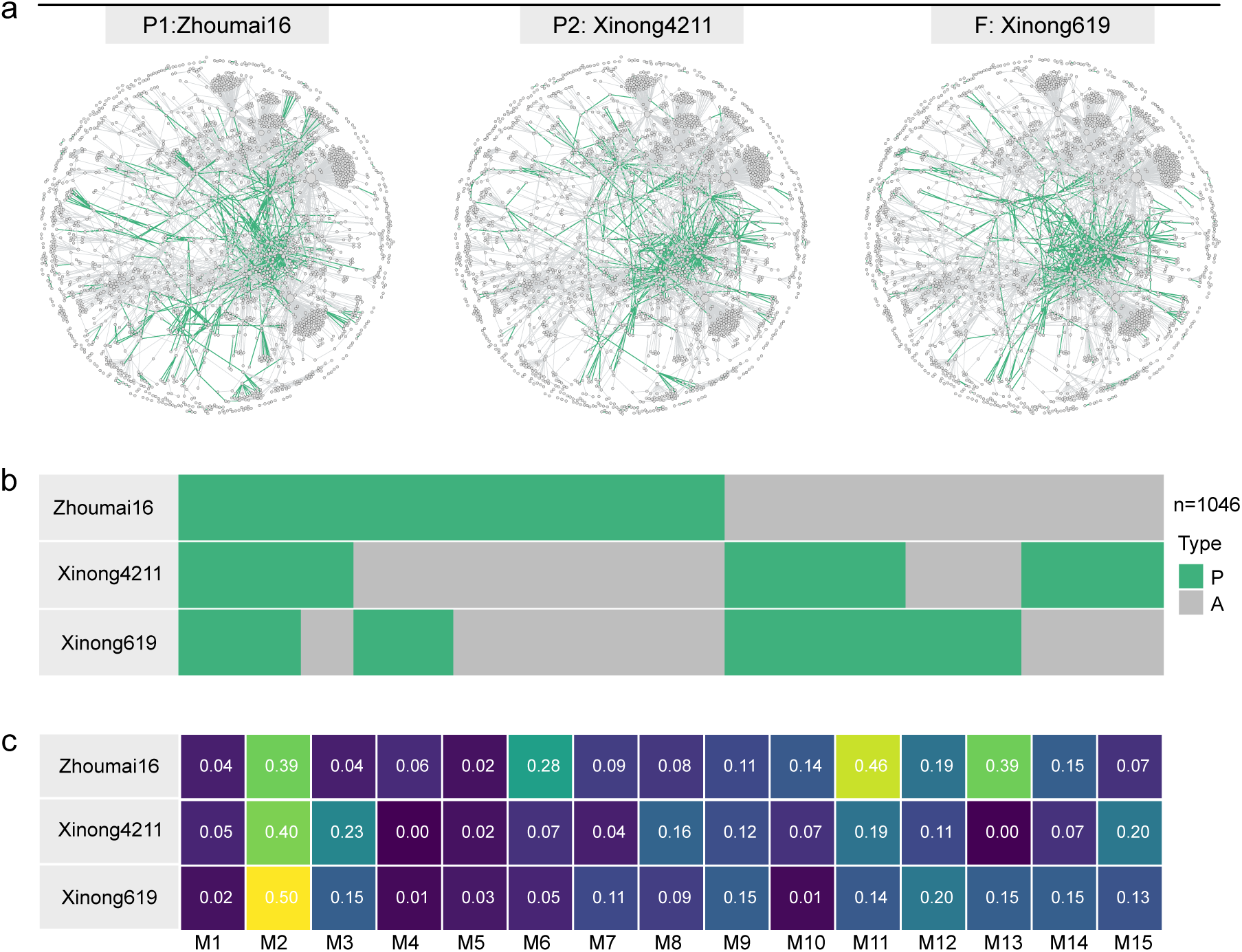
Inheriting and forming of favorable epistatic modules in breeding lineages. **a**, Epistatic networks of pedigree members. The pedigree includes ‘Zhoumai 16’ and ‘Xinong 4211’ as parents, and ‘Xinong 619’ as the progenies. Each point represents an HB, and each edge indicates a favorable epistatic interaction between two HBs. The green edges represent the presence of these epistasis in the wheat line, while the gray edges indicate their absence in the line. **b**, The presence-absence matrix of epistasis. Each column represents a favorable epistasis, with a total of 1,046 columns. The green columns represent the presence of these epistasis in the wheat line, while the gray columns indicate their absence in the line. This diagram clearly shows which epistases are shared between parents and progeny, and which are specific to individual genotypes. **c**, Completeness of the top 15 largest modules that were identified based on all accessions. The completeness of each module is calculated by dividing the number of edges in the module carried by each wheat line by the total number of edges in the module.

**Supplementary Fig. 27.**
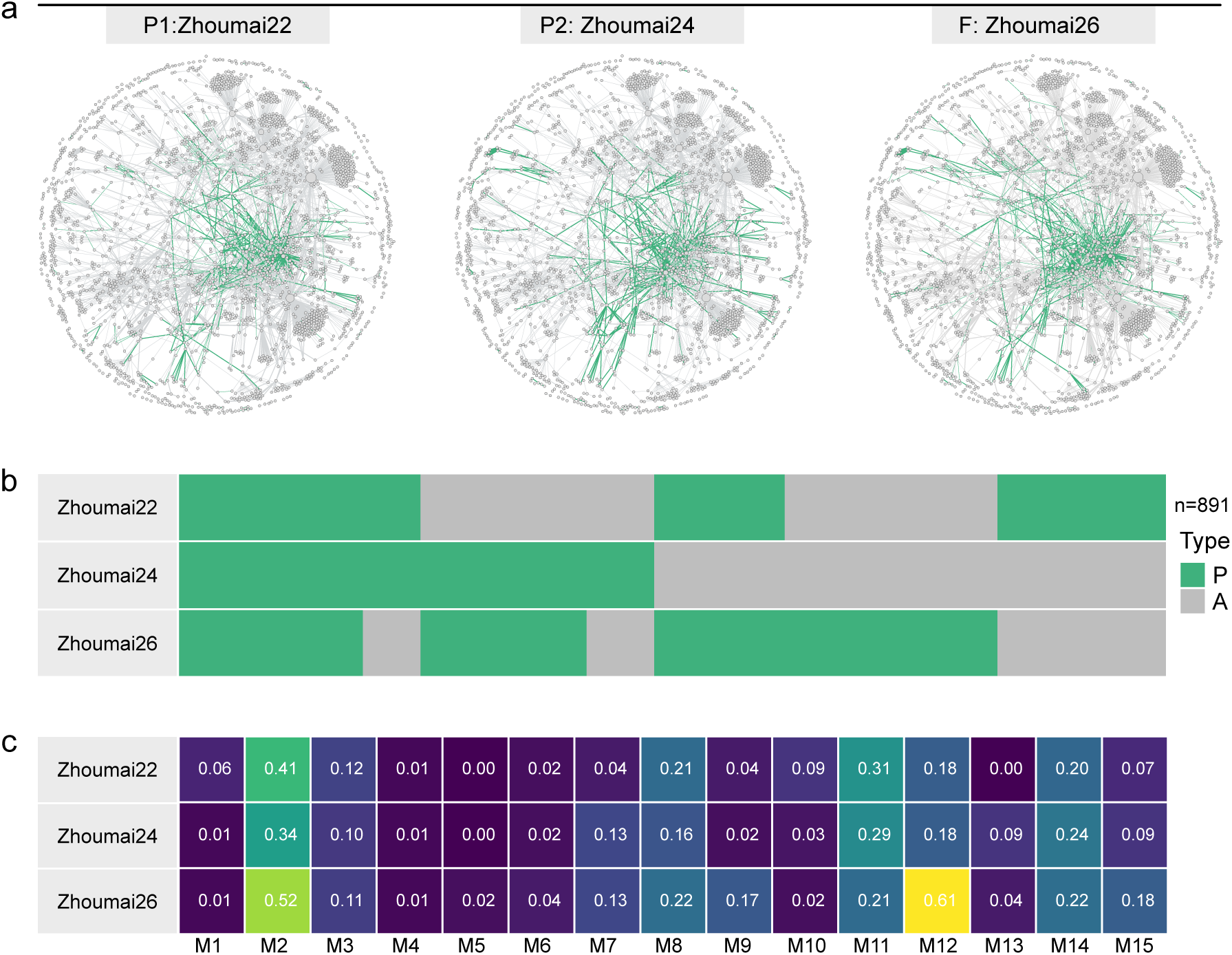
Inheriting and forming of favorable epistatic modules in breeding lineages. **a**, Epistatic networks of pedigree members. The pedigree includes ‘Zhoumai 22’ and ‘Zhoumai 24’ as parents, and ‘Zhoumai 26’ as the progenies. Each point represents an HB, and each edge indicates a favorable epistatic interaction between two HBs. The green edge represents the presence of this epistasis in the wheat line, while the gray edge indicates its absence in the line. **b**, The presence-absence matrix of epistasis. Each column represents a favorable epistasis, with a total of 891 columns. The green columns represent the presence of these epistasis in the wheat line, while the gray columns indicate their absence in the line. This diagram clearly shows which epistases are shared between parents and progeny, and which are specific to individual genotypes. **c**, Completeness of the top 15 largest modules that were identified based on all accessions. The completeness of each module is calculated by dividing the number of edges in the module carried by each wheat line by the total number of edges in the module.

**Supplementary Fig. 28.**
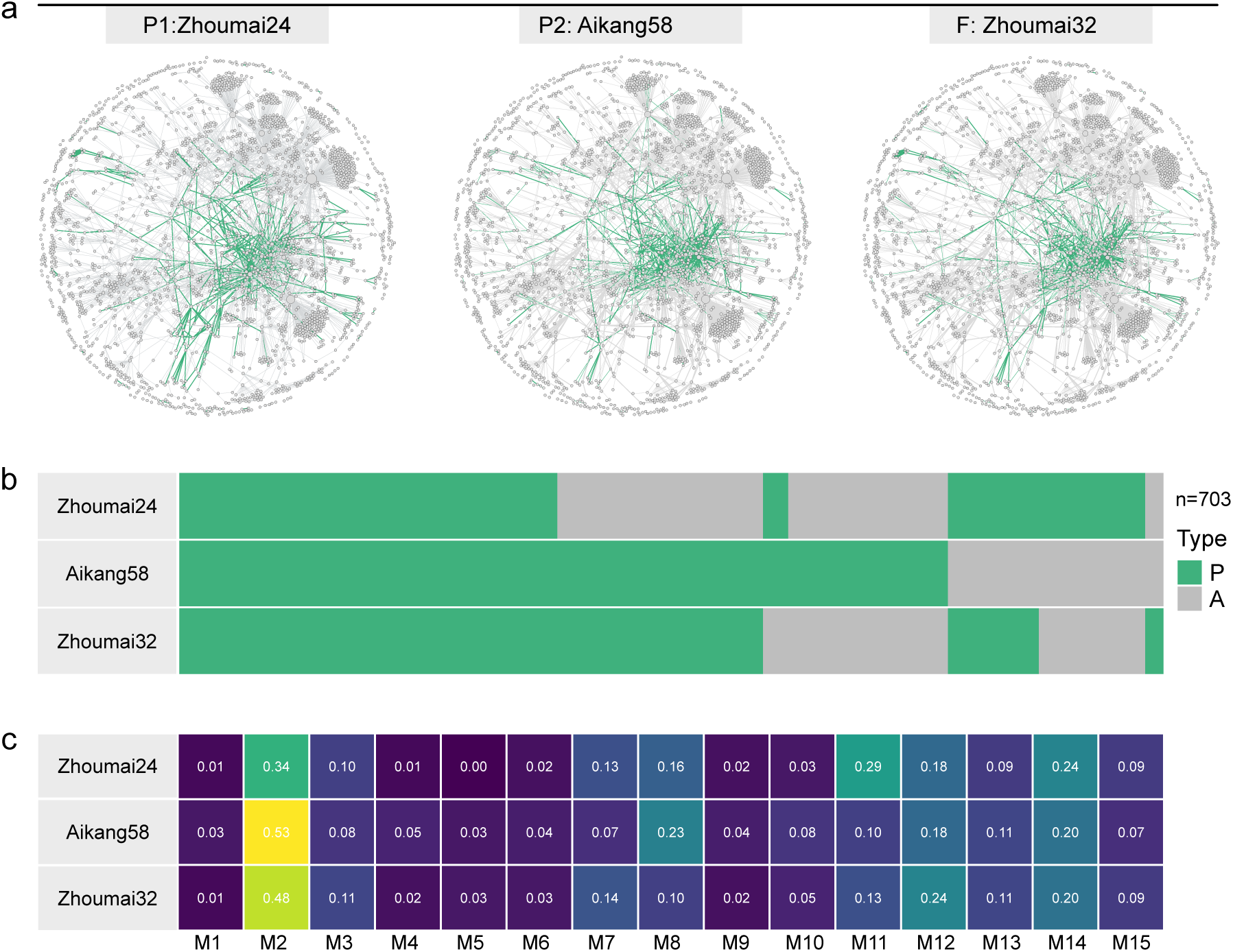
Inheriting and forming of favorable epistatic modules in breeding lineages. **a**, Epistatic networks of pedigree members. The pedigree includes ‘Zhoumai 24’ and ‘Aikang 58’ as parents, and six lines of ‘Zhoumai 32’ as the progenies. Each point represents an HB, and each edge indicates a favorable epistatic interaction between two HBs. The green edge represents the presence of this epistasis in the wheat line, while the gray edge indicates its absence in the line. **b**, The presence-absence matrix of epistasis. Each column represents a favorable epistasis, with a total of 703 columns. The green columns represent the presence of these epistasis in the wheat line, while the gray columns indicate their absence in the line. This diagram clearly shows which epistases are shared between parents and progeny, and which are specific to individual genotypes. **c**, Completeness of the top 15 largest modules that were identified based on all accessions. The completeness of each module is calculated by dividing the number of edges in the module carried by each wheat line by the total number of edges in the module.

**Supplementary Fig. 29.**
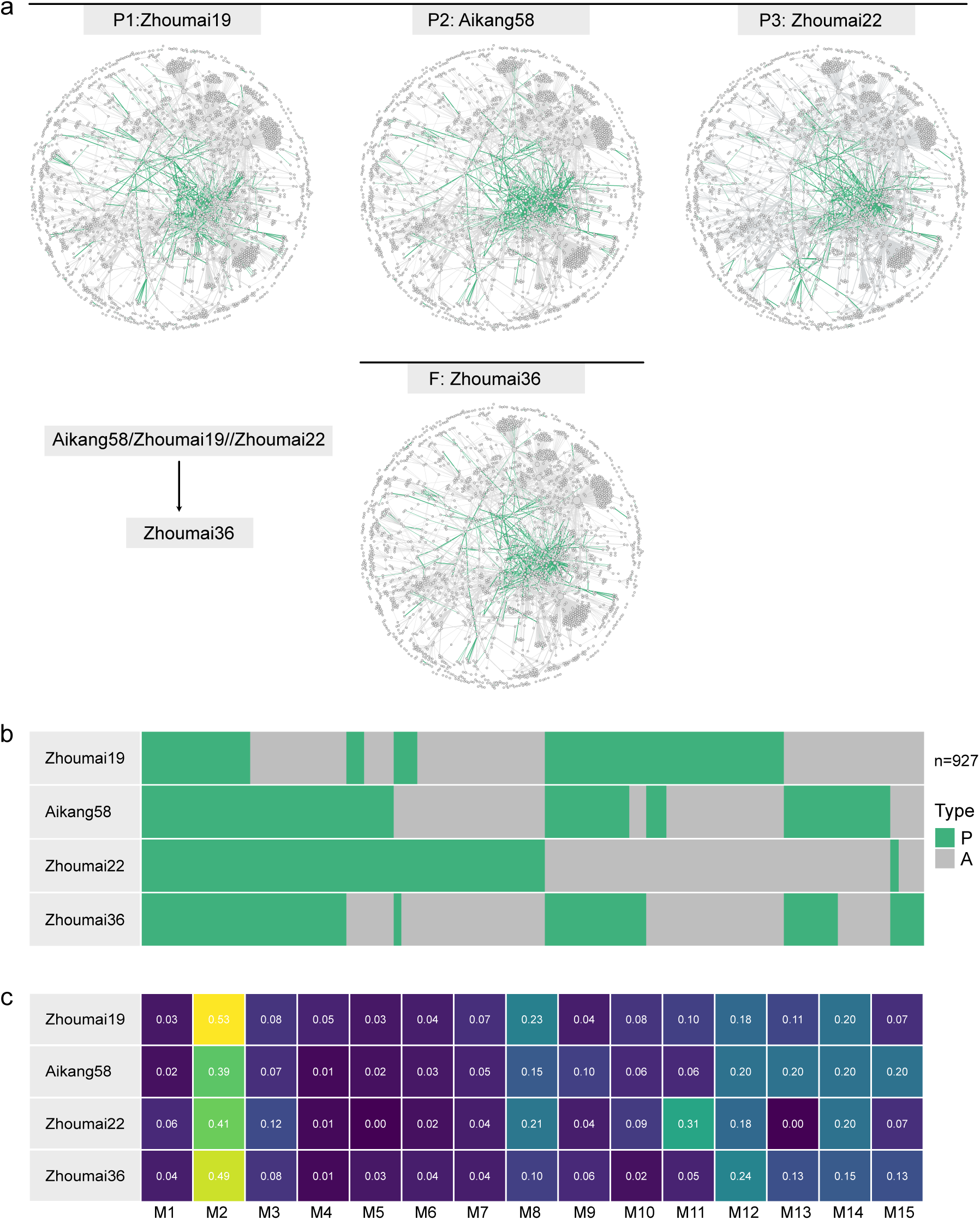
Inheriting and forming of favorable epistatic modules in breeding lineages. **a**, Epistatic networks of pedigree members. The pedigree includes ‘Zhoumai 19’, ‘Aikang 58’ and ‘Zhoumai 22’ as parents, and six lines of ‘Zhoumai 36’ as the progenies. ‘Zhoumai 36’ is a wheat variety developed by first crossing ‘Aikang 58’ as the female parent with ‘Zhoumai 19’ as the male parent, and then further crossing the resulting hybrid with ‘Zhoumai 22’. Each point represents an HB, and each edge indicates a favorable epistatic interaction between two HBs. The green edge represents the presence of this epistasis in the wheat line, while the gray edge indicates its absence in the line. **b**, The presence-absence matrix of epistasis. Each column represents a favorable epistasis, with a total of 927 columns. The green represents the presence of these epistasis in the wheat line, while the gray indicates their absence in the line. This diagram clearly shows which epistases are shared between parents and progeny and which are specific to individual genotypes. **c**, Completeness of the top 15 largest modules that were identified based on all accessions. The completeness of each module is calculated by dividing the number of edges in the module carried by each wheat line by the total number of edges in the module.

**Supplementary Fig. 30.**
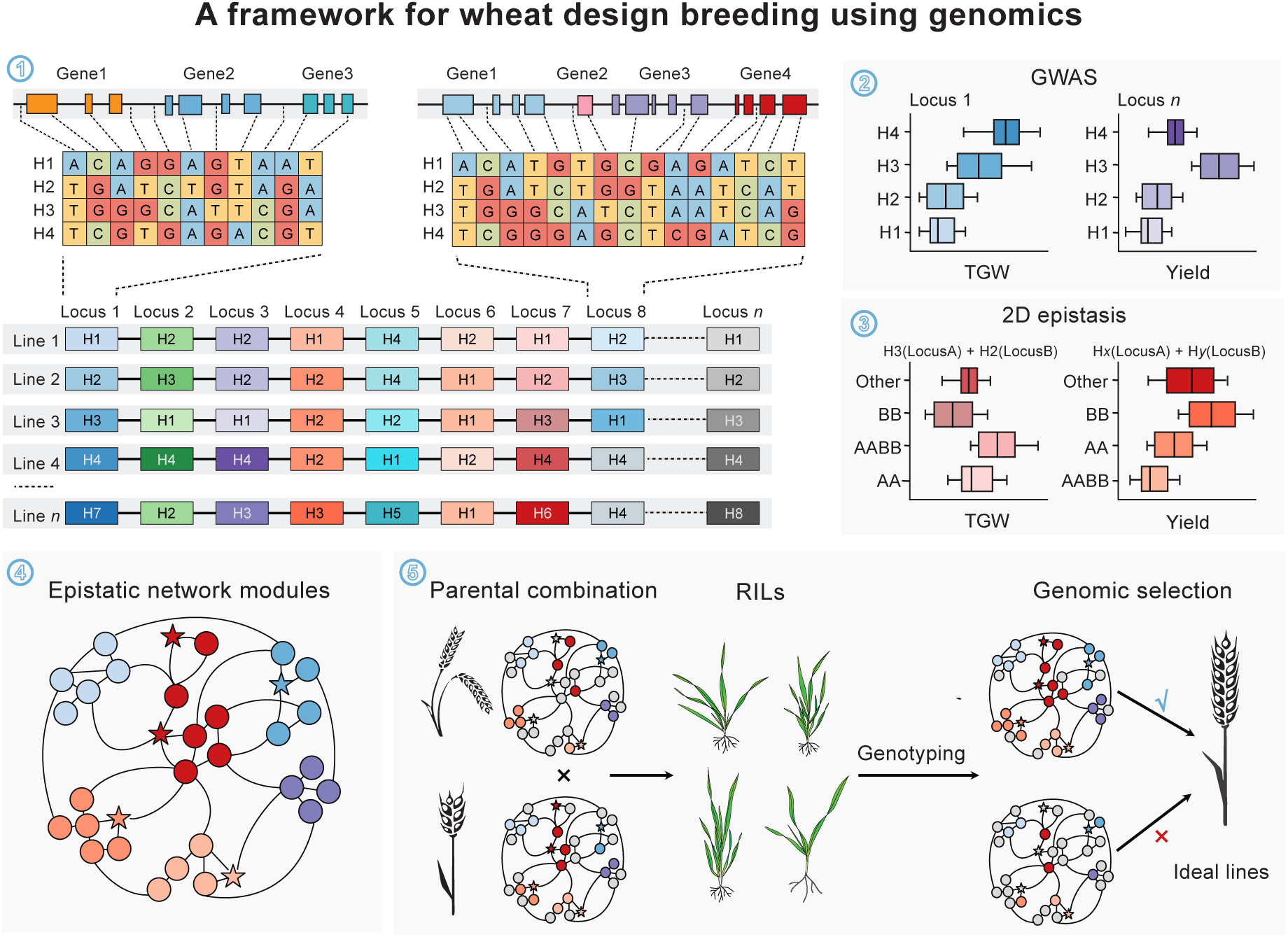
A framework for wheat design breeding through identification of epistasis modules. This figure references the work of Bevan et al^7^ in Section 1 and 2. This figure consists of five sections. Section 1 is the visualization of haplotype blocks. Section 2 represents haplotype-based GWAS analysis. Section 3 represents haplotype-based epistatic analysis. Section 4 represents the construction of the epistatic network and module identification. Section 5 includes the selection of appropriate parental combinations based on epistatic network modules, the construction of recombinant inbred lines, and genomic prediction and selection through sequencing and epistatic network module analysis.

### Supplementary Tables

**Supplementary Table 1 Information of 3030 accessions and 7 control varieties (CK 1-7)**.

**Supplementary Table 2 Statistics of SNP annotatio.**

**Supplementary Table 3 The mean nucleotide diversity (π) of different groups.**

**Supplementary Table 4 The mean divergence index (*F*_ST_) between groups.**

**Supplementary Table 5 The summary of the top 1% of XPCLR analysis.**

**Supplementary Table 6 GO enrichment of candidate genomic regions scanned by XP-CLR.**

**Supplementary Table 7 The information of inferred haplotype blocks.**

**Supplementary Table 8 The statistics of haplotype blocks in number and length.**

**Supplementary Table 9 The number of haplotype blocks along chromosomes.**

**Supplementary Table 10 The genomic contribution of Chinese landraces and introduced lines.**

**Supplementary Table 11 The alien chromosome segments inferred by SNP missing ratio.**

**Supplementary Table 12 The phenotypic BLUP of 14 traits based on 5 environment data over two consecutive years.**

**Supplementary Table 13 Phenotypic changes every 10 years.**

**Supplementary Table 14 The SEM analysis to quantify direct and indirect effects of yield components on grain yield.**

**Supplementary Table 15 Summary of quantitative trait loci (QTLs) of 14 traits.**

**Supplementary Table 16 QTLs associated with multiple traits.**

**Supplementary Table 17 HBs associated with 14 traits and their frequencies across era or geographical groups.**

**Supplementary Table 18 HBs associated with multiple traits.**

**Supplementary Table 19 Statistics and classification of epistasis.**

**Supplementary Table 20 The narrow-sense heritability of grain yield calculated by SNPs, HBs, GWAS hits, and epistasis.**

**Supplementary Table 21 The topological properties of favorable epistasis network for 3037 accessions.**

### Supplementary Data

**Supplementary Data 1 The presence-absence matrix of HBs across 3037 accessions.**

**Supplementary Data 2 The presence-absence matrix of epistasis across 3037 accessions.**

